# Activity-dependent myelination in prefrontal circuits signals the offset of infantile amnesia

**DOI:** 10.64898/2026.09.03.749197

**Authors:** A. Golbabaei, A.I. Ramsaran, A. Rashid, M.L. de Snoo, D. Karamboulas, J. Wo, B. Shahrbandi, Y. Li, B.S. Wang, A. Willis, M. Fukuchi, S. Kida, D.R. Kaplan, F.D. Miller, S.A. Josselyn, P.W. Frankland

## Abstract

Human and non-human infants can form memories for events, but these memories are not successfully consolidated into remote memory. While the neurobiological basis of this phenomenon—known as infantile amnesia—remains unclear, it is hypothesized that the neural circuits required for successful consolidation are insufficiently mature. Here we find that heightened activity in the prelimbic cortex of developing mice triggers a sequalae of maturational steps that culminates in adult-like memory persistence: Activity-dependent increases in brain-derived neurotrophic factor (BDNF) promote myelination of prelimbic circuits via activation of tyrosine kinase receptor B (TrkB) receptors on oligodendrocyte precursor cells (OPCs). Inhibiting any of these steps within a critical developmental window delays the offset of infantile amnesia, whereas promoting this sequalae results in the precocial emergence of memory persistence. Similar to critical periods in sensory cortices, our results indicate that developmental myelination is required for proper circuit maturation and emergence of adult-like memory function.

## INTRODUCTION

The persistence of event-based (or episodic^1^) memories differs in children compared to adults. While episodic memories acquired in adulthood may persist for months or years, those acquired during infancy fade quickly^2–4^. This loss of episodic memories from our earliest childhood years is known as infantile amnesia^5^ and typically spans the first 3-4 years of a person’s life^2,6^. Cultural factors^7^, language development^8^ and the emergence of self-identity^9^ influence the timing and nature of infantile amnesia. However, similar loss of event-based memories is observed in non-human animals^10,11^, including mice^12,13^, indicating that human-centered accounts of infantile amnesia are incomplete. Instead, neurobiologically-motivated accounts of infantile amnesia link the immaturity of the infant brain to memory loss^14–18^.

Infantile amnesia does not appear to be due to an inability to encode event-based memories. Human infants can form episodic memory (i.e., bind the ‘what-where-when’ details of an event) by ∼2 years of age^18–20^, with rudimentary aspects of episodic memory encoding even detectable in 12 month-old human infants^21^. Likewise, infant mice can form event-based memories by the end of the second post-natal week (i.e., P14) ^22^. Instead, the subsequent loss of these infant-acquired memories in human and non-human animals suggests that infantile amnesia reflects deficits in post-encoding processes, such as consolidation. In particular, the neural systems required for successful consolidation of event-based memories may be insufficiently mature for the generation of enduring, or remote, memories^6,14,15,19,23^.

In adult humans and rodents, the transformation of initially hippocampus-dependent event-based memories into remote memory (i.e., systems consolidation) depends on medial prefrontal structures, including the prelimbic (PrL) cortex^24,25^. Here, we causally link the maturational state of the PrL to the offset of infantile amnesia in mice, identifying a developmental program that is necessary for the emergence of adult-like memory persistence. Our results indicate that heightened PrL activity during infancy promotes BDNF release, activation of TrkB receptors on OPCs, oligodendrogenesis and subsequent myelination of PrL circuits. Inhibiting any step within a defined developmental window delays the offset of infantile amnesia. By contrast, promoting this cascade within the same critical developmental window leads to earlier infantile amnesia offset, allowing infant mice to precociously exhibit adult-like memory persistence.

## RESULTS

### Infantile amnesia offset occurs between P20 and P25

The binding of events to their surrounding spatial context is a core feature of episodic memory and may be studied in mice using contextual fear conditioning^26^. We evaluated retention of contextual fear memory in developing (postnatal day 15-30 [P15-30]) and adult (P60) mice. Mice were trained and memory was assessed at recent (2 days) and remote (15 days) delays by placing mice back into the conditioned context and measuring freezing behavior^27^ (Fig. 1a). Younger mice (P15-P20 or ≤P20) froze more in the recent compared to the remote test, indicating that they do not form remote memories for the conditioning experience. By contrast, older mice (≥P25) froze equivalently in the recent and the remote tests, reflecting successful remote memory consolidation (Fig. 1b and Extended Data Fig. 1a-e). To compare memory consolidation across development, we normalized freezing levels in the remote test to those in the recent test for each mouse. This analysis revealed that the offset of infantile amnesia occurs between P20 and P25, with mice ≥P25 able to successfully convert recent memory into remote memory (Fig. 1c).

**Fig. 1.**
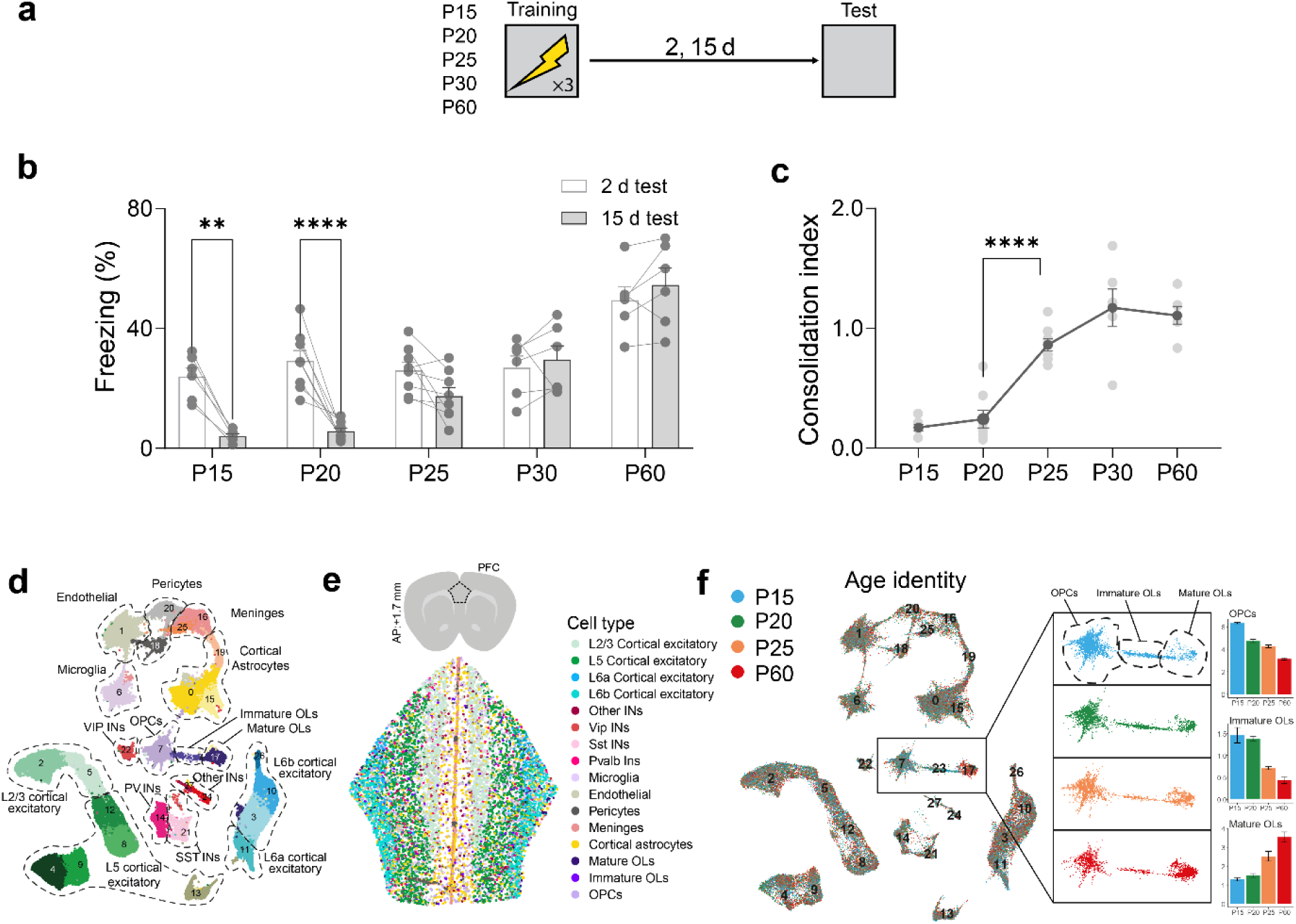
Infantile amnesia offset occurs between P20 and P25. **a,**Contextual fear conditioning was used to assess memory persistence across development. **b,** Younger mice (≤P20) froze more in recent than remote tests, whereas older mice (≥P25) showed comparable freezing in both tests (ANOVA, Age × Delay interaction: *F*_4,58_ = 6.84, *P* < 0.001). **c,** Consolidation index (remote/recent freezing) increased between P20 and P25, indicating infantile amnesia offset (ANOVA, main effect of Age: *F*_4,29_ = 31.34, *P* < 0.0001). **d-f,** Xenium spatial transcriptomics was performed on PrL sections from mice at P15, P20, P25, and P60 (n = 3/age). Cell transcriptomes were merged and annotated. UMAPs showing (**d**) cell-type annotations, (**e**) spatial plots of cell types in a sample section of PrL, and (**f**) age-coded transcriptomes (P15 blue, P20 green, P25 orange, P60 red). Bar plots show proportions of OPCs, immature OLs, and mature OLs per sample.

In human and non-human adults, subregions of the medial prefrontal cortex, including the PrL, are important for conversion of recent into remote memory^28–32^. Using post-training chemogenetic inhibition of the PrL^33^, we confirmed that PrL activity during the post-training period is necessary for successful remote memory consolidation in developing (P25) and adult (P60) mice (but not at younger ages prior to the offset of infantile amnesia) (Extended Data Fig. 1f–i).

### Developmental emergence of a myelination-promoting transcriptional program in PrL

To identify molecular changes in the PrL that span the offset of infantile amnesia, we performed spatial single-cell transcriptomic profiling of PrL across four developmental ages (P15, P20, P25, and P60). Using the Xenium platform with a custom 347-gene probe set (Table S1) enriched for developmental regulators, we profiled 103,878 cells, ∼57% of which were neurons^34^. UMAP visualization and spatial mapping revealed clear clustering, and canonical marker genes identified the expected neuronal and glial cell classes (Fig. 1d,e and Extended Data Fig. 2a-c, Extended Data Table 2). While most cell-type proportions remained stable across ages, we observed marked developmental shifts in the oligodendrocyte lineage: OPCs and immature oligodendrocytes were abundant at early ages, whereas mature oligodendrocytes increased by adulthood (Fig. 1f; for other cell-types see Extended Data Table 3), consistent with known postnatal cortical myelination trajectories^35^.

To explore these changes in greater detail, we subsetted and reclustered oligodendrocyte-lineage cells (Fig. 2a,b, and Extended Data Fig. 2d,e, and Extended Data Table 4). This analysis confirmed progressive transitions from OPC → immature → mature oligodendrocytes across development (Fig. 2c,d). Spatial mapping further revealed that OPCs and immature oligodendrocytes were broadly distributed throughout the PrL, whereas mature oligodendrocytes were concentrated in deep layers adjacent to the corpus callosum (Fig. 2e), consistent with previous single cell and spatial transcriptomic studies^36^. To identify molecular pathways linked to these developmental shifts, we performed differential gene expression analyses comparing juvenile (P15–P25) and adult (P60) oligodendrocyte-lineage cells. Across these comparisons, we identified many developmentally regulated genes (105 upregulated [Extended Data Table 5], 65 downregulated [Extended Data Table 6]), including genes that were most enriched from P15– P20, a window that immediately precedes the emergence of adult-like remote memory consolidation (Fig. 2f-i and Extended Data Fig. 3).

**Fig. 2.**
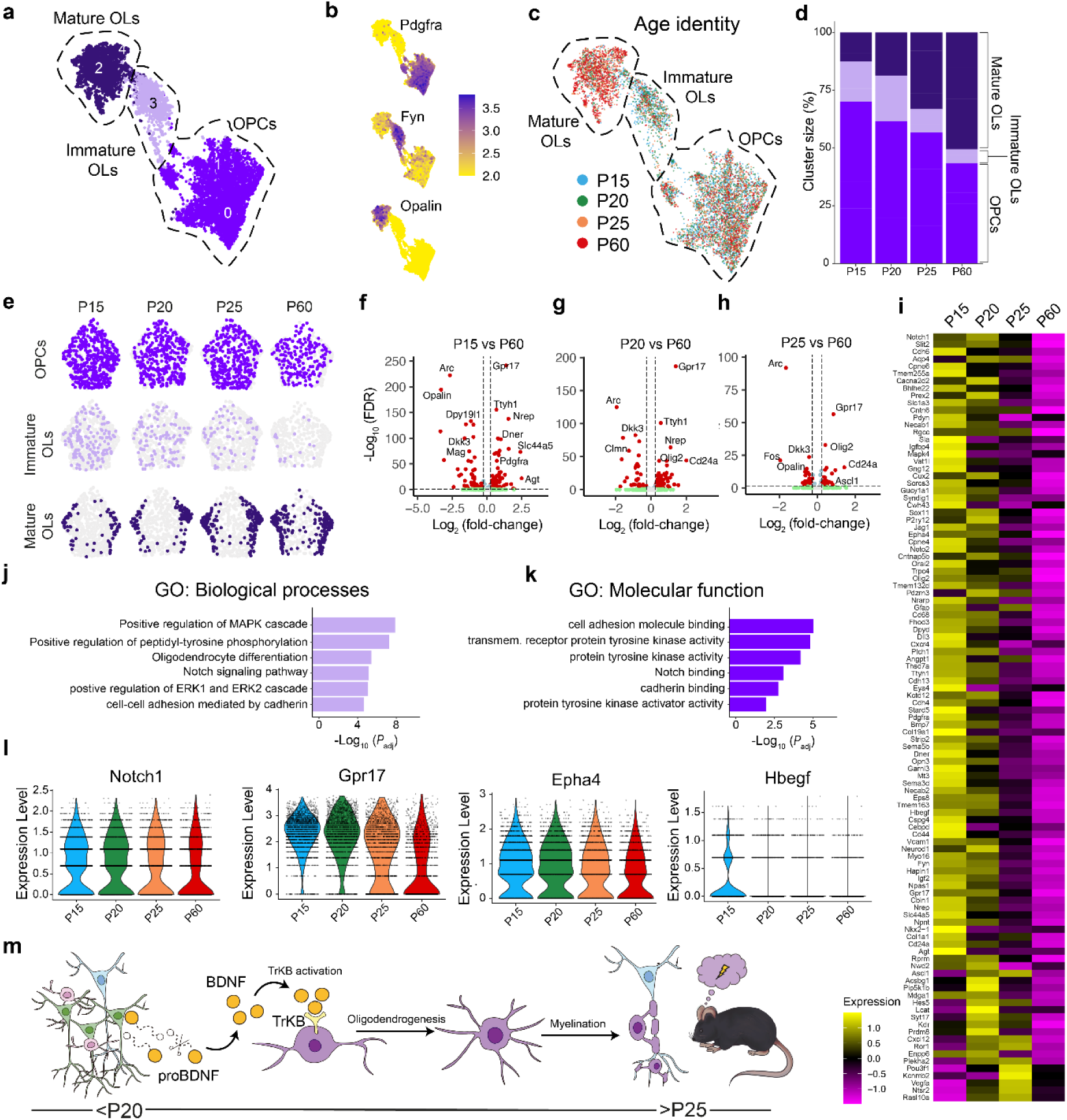
Developmental shifts in PrL oligodendrocyte-lineage composition. **a-c**, UMAPs of oligodendrocyte-lineage cells only, displaying (**a**) subtype annotations, (**b**) marker expression (Pdgfra, Fyn, Opalin), and (**c**) age-coded cells. **d**, Lineage composition (% OPCs/immature oligodendrocytes/mature oligodendrocytes) across ages. **e**, Spatial maps of lineage subtypes within representative PrL sections at each age. **f-h**, Volcano plots of differentially expressed genes (red; adj *P* value < 0.05, and log_2_(FC) > 0.25) for P15, P20, and P25 versus P60 (x-axis indicates fold-change [log _2_], y-axis represents -log_10_ [FDR]). **i**, Average expression heatmap showing genes upregulated at P20–P25 relative to P60. **j**,**k**, GO terms enriched among developmentally upregulated genes (molecular functions; biological processes). **l**, Violin plots of selected developmentally regulated genes (Notch1, Ascl1, Epha4, Hbegf). **m**, Working model for developmental sequalae of events in PrL leading to infantile amnesia offset. Neural activity → proBDNF to mature BDNF conversion → TrkB activation on OPCs → oligodendrogenesis and myelination.

Gene ontology analyses revealed that P15-P20 oligodendrocyte-lineage cells were enriched for pathways associated with oligodendrocyte differentiation, Notch signaling, and tyrosine kinase receptor activity (Fig. 2j-l, and Extended Data Tables 7 and 8), all of which contribute to regulating the timing of oligodendrocyte maturation^37–40^. In particular, increased tyrosine kinase receptor (Trk) activity raises the possibility that TrkB-dependent progression of OPCs toward mature myelinating oligodendrocytes^40–42^ may be the key developmental milestone signaling the offset of infantile amnesia. According to this hypothesis (Fig. 2m), infantile amnesia offset depends on a temporally ordered sequence of developmental events. Neural activity in the PrL first triggers BDNF release, which promotes TrkB-dependent differentiation of OPCs into mature oligodendrocytes and subsequent myelination of prefrontal circuits. The resulting myelination of prefrontal circuits then permits the conversion of recent into remote memory, supporting the emergence of adult-like memory function.

### Coordinated PrL developmental changes coincide with infantile amnesia offset

We next asked how neural activity, trophic signaling, and oligodendrocyte maturation in PrL change across the developmental period in which remote memory first emerges. To track developmental changes in neural activity in the PrL, we quantified expression of the activity-dependent immediate early gene, c-Fos, in mice from P15-P60. Similar to other activity-regulated genes^43^, c-Fos levels were elevated in younger mice (≤P20) taken from the home cage (Fig. 3a-c and Extended Data Fig. 4a,b). Coinciding with heightened neural activation, western blot analysis revealed a pronounced peak in mature BDNF (and concomitant decrease in proBDNF) levels in PrL in home cage mice at ages P19–P20 (Fig. 3d-f and Extended Data Fig. 4c-e), consistent with an activity-dependent conversion of BDNF from its immature, uncleaved form^44^ (for converging results using a BDNF-luciferase reporter mouse see Extended Data Fig. 4f-h). Given that mature BDNF is the ligand for TrkB^45^, we next assessed TrkB activation using phospho-TrkB (pTrkB) immunohistochemistry (Fig. 3g-j). pTrkB signal closely mirrored the temporal profile of mature BDNF, with elevated staining in OPCs at P20, indicating that the transient increase in BDNF coincides with activation of TrkB signaling in the developing PrL.

**Fig. 3.**
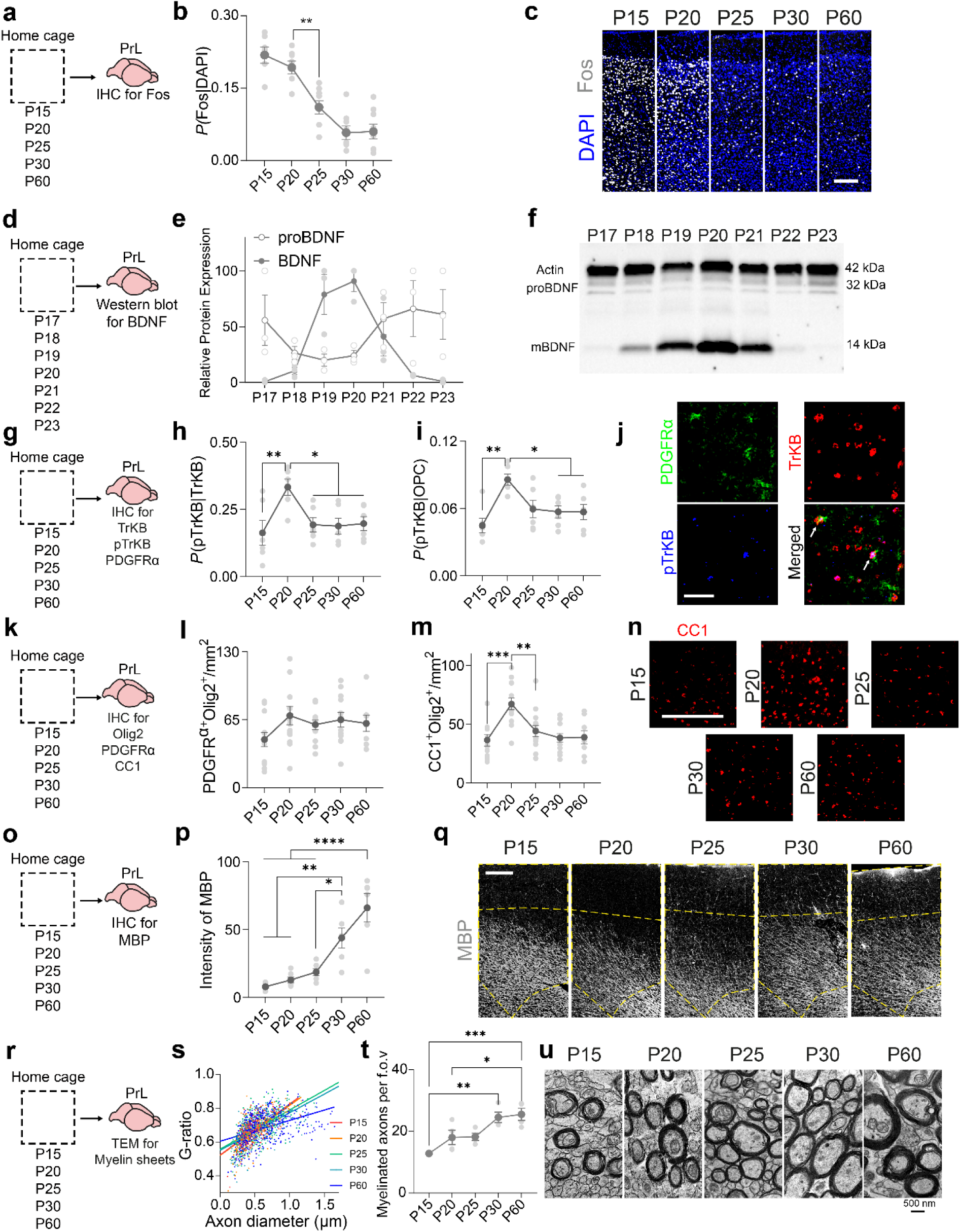
Developmental changes in activity, BDNF-TrkB signaling, and oligodendrocyte maturation in PrL. **a-c**, Fos staining in home-cage mice shows elevated PrL activity at P15– P20 (ANOVA, effect of Age: *F*_4,32_ = 24.29, *P* < 0.0001). Scale bar: 200 µm. **d-f**, Western blots from PrL (P17–P23) show a peak in BDNF at P20, while proBDNF remains stable. Actin served as loading control (BDNF ANOVA, effect of Age: *F*_6,14_ = 14.18, *P* < 0.0001. proBDNF ANOVA, effect of Age: *F*_6,14_ = 1.44, *P* = 0.27). **g-j**, PrL sections stained for TrkB, pTrkB, and PDGFRα. pTrkB levels peaked at P20 in both TrkB⁺ cells (ANOVA, effect of Age: *F*_4,25_ = 4.36, *P* < 0.01) and OPCs (ANOVA, effect of Age: *F*_4,25_ = 5.59, *P* < 0.01). Scale bar: 50 µm. **k-n**, Oligodendrocyte lineage markers (Olig2, PDGFRα, CC1) reveal stable OPC numbers (ANOVA, effect of Age: *F*_4,53_ = 1.56, *P* = 0.20) but a transient increase in mature oligodendrocytes at P20 (ANOVA, effect of Age: *F*_4,53_ = 7.41, *P* < 0.0001). Scale bar: 200 µm. **o-q**, MBP expression increased after P25 (ANOVA, effect of Age: *F*_4,25_ = 16.94, *P* < 0.0001). Scale bar: 200 µm. **r-u**, Transmission electron microscopy (TEM) analyses revealed increased myelin sheath thickness (G-ratio) (Linear regression: P15 slope = 0.30, P20 slope = 0.29, P25 slope = 0.23, P30 slope = 0.22, P60 slope = 0.12; ANOVA, effect of Age: *F*_4,1576_ = 18.04, *P* < 0.001) and more myelinated axons with age (ANOVA, effect of Age: *F*_4,15_ = 9.87, *P* < 0.001). Scale bar: 500 nm.

To determine whether this transient surge in BDNF-TrkB signaling influences oligodendrogenesis, we performed immunohistochemistry for general oligodendrocyte-lineage marker (Olig2), OPCs (PDGFRα) and differentiated, mature oligodendrocytes (CC1) (Fig. 3k). While the number of OPCs (i.e., Olig2⁺PDGFRα⁺) remained stable, we observed an increase in differentiated oligodendrocytes (Olig2⁺CC1⁺) at P20, coinciding with the window of elevated neural activity and TrkB activation (Fig. 3l–n and Extended Data Fig. 4i–m). Similar transient increases in differentiated oligodendrocytes have been reported during postnatal development in other CNS regions, including motor cortex^46^, corpus callosum^47,48^ and spinal cord^49^, although the precise developmental timing differs across structures. These transient peaks are thought to reflect periods of heightened oligodendrocyte differentiation, during which newly generated OPCs transition into differentiated oligodendrocytes. Many of these newly differentiated oligodendrocytes are subsequently eliminated during normal cortical development^50,51^, potentially contributing to the reduction in Olig2⁺CC1⁺ cells after P20. Consistent with this, thymidine labeling using EdU revealed increased oligodendrogenesis between P15 and P20 (Extended Data Fig. 4n-q), confirming that newly generated oligodendrocytes mature during this developmental period.

These developmental changes in BDNF-TrkB signaling preceded myelination in PrL: Immunohistochemical staining for myelin basic protein (MBP) revealed an age-dependent increase in MBP staining in mice ≥P20 for both inhibitory and excitatory neurons (Fig. 3o-q and Extended Data Fig. 4r-u). Consistent with this, transmission electron microscopy revealed age-dependent increases in axon diameter and the proportion of myelinated axons, indicating enhanced myelination during this developmental window (Fig. 3r-u).

### Activity-dependent myelination in PrL is required for the developmental emergence of remote memory

Our results support a model whereby activity-dependent BDNF release drives TrkB-mediated differentiation of OPCs into mature, myelinating oligodendrocytes in the PrL, permitting adult-like remote memory. To address whether these events are necessary for infantile amnesia offset, we asked whether inhibiting these upstream regulators of PrL myelination would extend the period of infantile amnesia and delay the emergence of adult-like memory persistence.

Previous studies have shown that chemogenetic silencing of neural activity during development can delay maturation of entorhinal–hippocampal circuits^52^. Using a similar approach, we asked whether suppressing PrL activity during development would interfere with activity-dependent processes required for normal oligodendrocyte maturation and myelination. To do this, we expressed the inhibitory DREADD hM4Di in PrL excitatory neurons and administered C21 during discrete 5-day developmental windows. We varied the timing of inhibition (P15–P19, P20–P24 or P25–P29) such that each manipulation preceded training at P20, P25 or P30, respectively (Fig. 4a–c). In VEH-treated controls, the developmental increase in differentiated oligodendrocytes emerged between P20 and P25, alongside the successful conversion of recent into remote memory at P25, consistent with previously observed developmental timing. By contrast, suppressing PrL activity delayed the developmental increase in differentiated oligodendrocytes and infantile amnesia offset (Fig. 4d,e and Extended Data Fig. 5a–g).

**Fig. 4.**
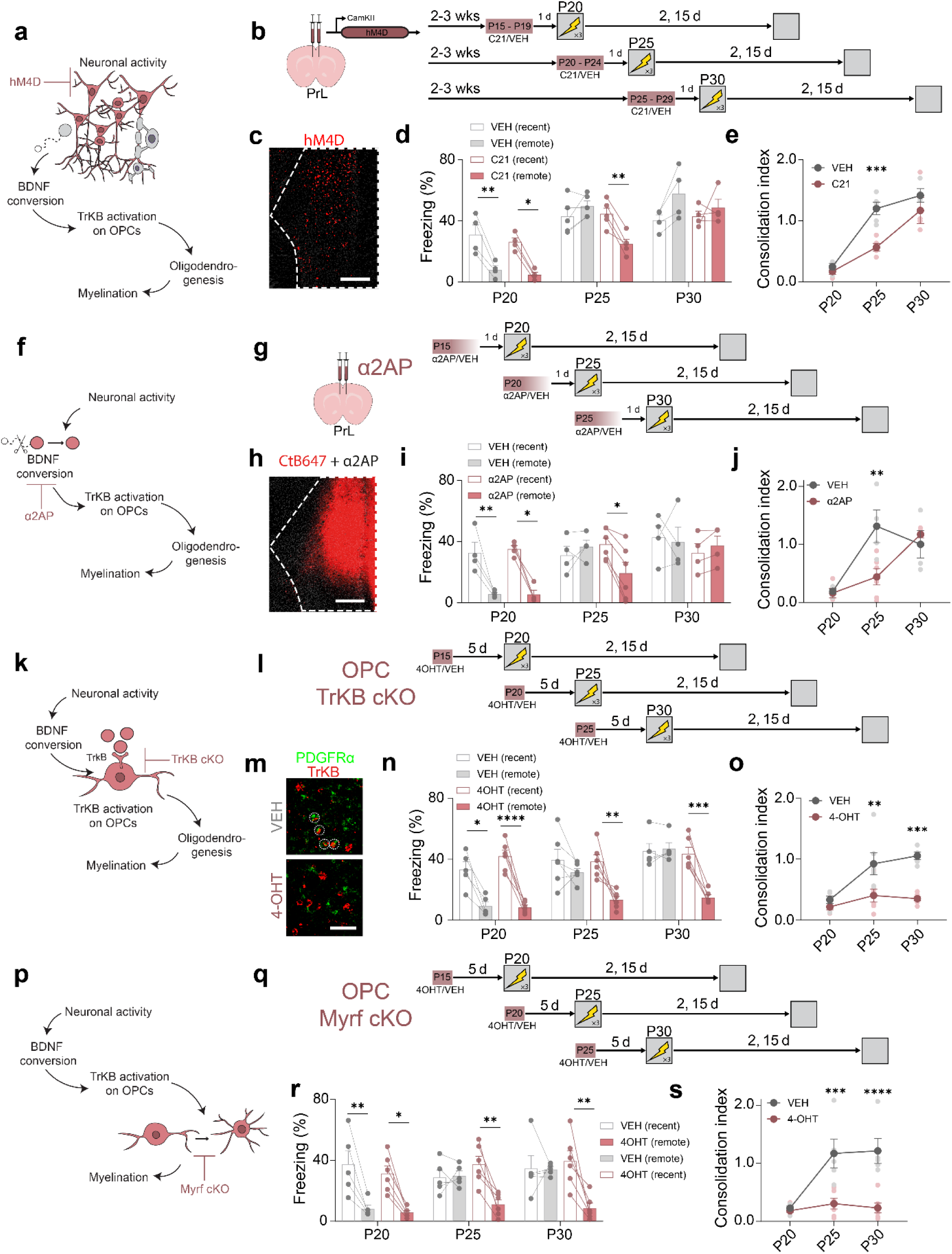
Inhibiting neural activity, BDNF-TrkB signaling, or oligodendrocyte maturation in PrL delays infantile amnesia offset. **a**,**b**, To suppress PrL neural activity at different developmental stages, mice were microinjected with an AAV expressing inhibitory DREADD, hM4D_i_ and administered C21 or VEH for 5 days. One day following C21/VEH treatment, mice were conditioned at P20, P25, or P30 and tested 2 and 15 days later. **c**, hM4D_i_ expression in PrL. Scale bar: 200 µm. **d**, Freezing at recent and remote delays for VEH- and C21-treated mice trained at P20, P25, or P30 (ANOVA, Age × Inhibition × Delay interaction: *F*_2,20_ = 4.68, *P* < 0.05). **e**, The offset of infantile amnesia was delayed in C21-treated mice (ANOVA, Age × Inhibition interaction: *F*_2,20_ = 3.64, *P* < 0.05). **f**,**g**, To block proBDNF→BDNF conversion at different developmental stages, α2AP (or VEH) was microinjected into the PrL. Five days later, mice were conditioned at P20, P25, or P30 and tested 2 and 15 days later. **h**, Representative injection (α2AP was mixed with CTb-647 for visualization). Scale bar: 200 µm. **i**, Freezing at recent and remote delays for VEH- and α2AP-treated mice trained at P20, P25, or P30 (ANOVA, Age × α2AP × Delay interaction: *F*_2,20_ = 3.94, *P* < 0.05). **j**, The offset of infantile amnesia was delayed in α2AP-treated mice (ANOVA, Age × α2AP interaction: *F*_2,20_ = 5.80, *P* < 0.05). **k**,**l**, In TrKB_CKO_ mice, 4-OHT administration permitted OPC-specific TrkB deletion at different developmental stages. Five days later, mice were conditioned at P20, P25, or P30 and tested 2 and 15 days later. **m**, TrkB loss was confirmed in OPCs (scale bar: 50 µm) **n**, Freezing at recent and remote delays for VEH- and 4-OHT-treated mice trained at P20, P25 or P30 (ANOVA, TrKB-KO × Delay interaction: *F*_1,28_ = 20.80, *P* < 0.0001). **o**, OPC-specific TrkB deletion impaired consolidation (ANOVA, Age × TrKB-KO interaction: *F*_2,28_ = 4.35, *P* < 0.05). **p**,**q**, In Myrf_CKO_ mice, 4-OHT administration permitted OPC-specific Myrf deletion at different developmental stages. Five days later, mice were conditioned at P20, P25, or P30 and tested 2 and 15 days later. **r**, Freezing at recent and remote delays for VEH- and 4-OHT-treated mice trained at P20, P25 or P30 (ANOVA, Age × Myrf-KO × Time-point interaction: *F*_2,27_ = 4.21, *P* < 0.05). **s**, OPC-specific Myrf deletion impaired consolidation (ANOVA, Age × Myrf-KO interaction: *F*_2,27_ = 7.15, *P* < 0.01).

Blocking the conversion of proBDNF to mature BDNF in PrL using α2-antiplasmin (α2AP), an inhibitor of the tPA–plasmin pathway that normally cleaves proBDNF^53^, produced a similar pattern of results. Oligodendrocyte maturation was delayed and the emergence of remote memory postponed until P30 in α2AP-treated mice (Fig. 4f-j and Extended Data Fig. 5h-n). Therefore, disrupting either neural activity or activity-dependent BDNF conversion during this window delays the developmental trajectory of PrL oligodendrogenesis and offset of infantile amnesia. These same manipulations did not affect learning or retrieval since recent memory was intact.

Given that PrL activity and BDNF availability regulate the timing of oligodendrogenesis, we next asked whether downstream effectors—TrkB signaling in OPCs and their differentiation into myelin-forming oligodendrocytes—are required for remote memory consolidation. OPC-specific conditional deletion of TrkB or of Myrf, which blocks OPC differentiation into myelinating oligodendrocytes^40^, each disrupted oligodendrocyte maturation in the PrL and impaired remote memory in P25–P30 mice (Fig. 4k-s and Extended Data Fig. 5o-ab). These findings support a model in which TrkB-dependent generation of myelinating oligodendrocytes is necessary for the transition out of infantile amnesia.

### Promoting BDNF–TrkB signaling in a key developmental window leads to the precocious emergence of adult-like memory persistence

Given that inhibiting BDNF-TrkB signaling in a key developmental window delays oligodendrocyte maturation and the offset of infantile amnesia, we reasoned that promoting BDNF–TrkB signaling in PrL might have the opposite effect and lead to the precocial emergence of adult-like memory persistence. To test this, we microinjected recombinant BDNF or a TrkB agonist antibody (38B8, which promotes TrkB activation^54^) into PrL in developing mice. Mice were trained 5 days after this treatment at P15, P20, P25, or P30. Contextual fear memory was assessed at recent and remote delays (Fig. 5a-c).

**Fig. 5.**
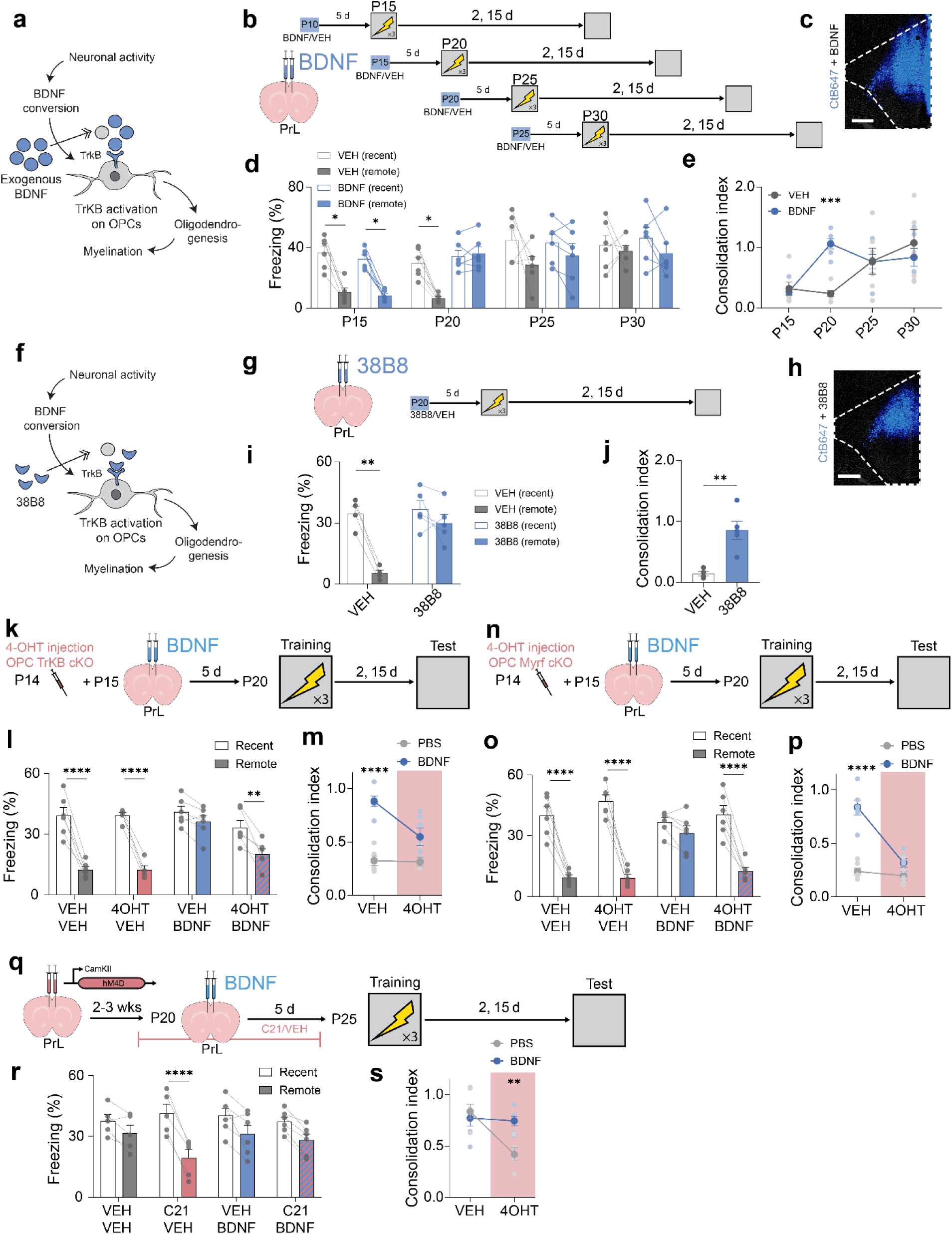
Promoting BDNF–TrkB signaling accelerates the emergence of adult-like memory persistence via TrkB-dependent oligodendrogenesis. a,b, To test whether BDNF accelerates the onset of adult-like memory persistence, recombinant BDNF (or VEH) was microinjected into the PrL 5 days prior to conditioning at P15, P20, P25 or P30. Mice were tested at recent (2 days) and remote (15 days) delays. **c**, Representative BDNF injection site (BDNF was mixed with CTb-647). Scale bar: 200 µm. **d**, Freezing at recent and remote delays for VEH- and BDNF-injected mice trained at P15–P30. BDNF advanced the age at which remote memory emerged (ANOVA, Age × BDNF interaction: *F*_3,40_ = 2.60, *P* = 0.06). **e**, The offset of infantile amnesia was advanced in BDNF-treated mice (ANOVA, Age × BDNF interaction: *F*_3,40_ = 5.62, *P* < 0.01). **f**,**g**, To assess whether TrkB activation is sufficient to induce precocious adult-like memory persistence, the TrkB agonist 38B8 (or VEH) was microinjected into the PrL at P15. Mice were trained at P20 and tested 2 and 15 days later. **h**, Representative 38B8 injection site. Scale bar: 200 µm. **i**, 38B8-treated mice exhibited both recent and remote recall, unlike controls (ANOVA, Time-point × 38B8 interaction: *F*_1,7_ = 11.32, *P* < 0.05). **j**, Successful remote memory consolidation in 38B8-treated P20 mice (unpaired t test: *t* = 4.60, *P* < 0.01). **k**, To test whether BDNF acts through TrkB on OPCs, TrKB_cKO_ mice received 4-OHT (or VEH) at P14 and intra-PrL microinjection of BDNF (or VEH) at P15. Mice were conditioned at P20 and tested 2 and 15 days later. **l**, Freezing at recent and remote delays. BDNF-induced advancement of infantile amnesia offset was blocked by OPC-specific TrkB deletion (ANOVA, TrkB-KO × BDNF interaction: *F*_1,19_ = 5.70, *P* < 0.05). **m**, Consolidation enhancement by BDNF required OPC TrkB (ANOVA, TrkB-KO × BDNF interaction: *F*_1,19_ = 7.88, *P* < 0.05). **n**, To test whether BDNF acts through oligodendrogenesis, Myrf_cKO_ mice received 4-OHT (or VEH) at P14 and intra-PrL microinjection of BDNF (or VEH) at P15. Mice were conditioned at P20 and tested 2 and 15 days later. **o**, Freezing at recent and remote delays. BDNF-induced advancement of infantile amnesia offset was blocked by OPC-specific Myrf deletion (ANOVA, Myrf-KO × BDNF × Delay: *F*_1,20_ = 5.06). **p**, Consolidation enhancement by BDNF required Myrf (ANOVA, Myrf-KO × BDNF interaction: *F*_1,20_ = 25.19, *P* < 0.0001). **q**, To determine whether BDNF can overcome reduced PrL activity, an AAV-hM4D_i_ was injected into PrL, followed by C21 from P20–P24 and BDNF (or VEH) at P21. Mice were trained at P25 and tested 2 and 15 days later. **r**, PrL silencing impaired remote recall, but co-administration of BDNF restored adult-like memory (ANOVA, Inhibition × BDNF × Delay: *F*_1,18_ = 9.87, *P* < 0.01). **s**, BDNF rescued the PrL silencing-induced consolidation deficit (ANOVA, inhibition × BDNF interaction: *F*_1,18_ = 8.55, *P* < 0.01).

Enhancing BDNF–TrkB signaling accelerated oligodendrocyte maturation, shifting the peak in number of mature oligodendrocytes to earlier ages (Extended Data Fig. 6a-e). Consistent with this, the offset of infantile amnesia also occurred earlier. While successful conversion of recent into remote memory emerged at P25 in VEH-treated mice, BDNF-treated mice exhibited robust remote memory already at P20 (Fig. 5d,e and Extended Data Fig. 6f,g). A similar pattern of results was observed following 38B8 treatment (Fig. 5f-j), indicating that promoting BDNF– TrkB signaling during this developmental window advances the maturation of PrL myelination and leads to the earlier emergence of adult-like memory persistence.

Although enhancing BDNF–TrkB signaling accelerated the emergence of remote memory, BDNF has many cellular targets^45^. We therefore asked whether the BDNF-dependent advance in infantile amnesia offset specifically requires TrkB signaling in OPCs and their subsequent differentiation into myelinating oligodendrocytes. To test this, we used a 2×2 design in P20 mice, which normally fail to express remote memory. As before, intra-PrL BDNF administration resulted in the precocial emergence of remote memory, with BDNF-treated mice successfully converting recent into remote memory. However, when either TrkB or Myrf was conditionally deleted in OPCs, BDNF was no longer effective: P20 mice continued to show impaired remote memory, similar to untreated developmental controls (Fig. 5k-p). These findings indicate that the ability of BDNF to accelerate the emergence of adult-like memory persistence requires TrkB signaling in OPCs and their differentiation into myelinating oligodendrocytes, causally linking developmental myelination in PrL to infantile amnesia offset.

To further test whether BDNF acts downstream of neural activity in this developmental pathway, we performed a converse experiment in P25 mice. Although chemogenetic silencing of PrL excitatory neurons for 5 days preceding training impaired conversion of recent into remote memory, neural activity could influence multiple processes beyond BDNF release. To determine whether BDNF is the principal mediator, we combined PrL neuronal silencing with intra-PrL BDNF injections (Fig. 5q). Consistent with our previous observations, VEH/VEH controls exhibited normal remote memory, whereas chemogenetic silencing alone impaired remote memory. Remarkably, this deficit was fully rescued by BDNF treatment (Fig. 5r and Extended Data Fig. 5s), indicating that activity-dependent BDNF release is a key mediator linking PrL neuronal activity to OPC maturation and the emergence of remote memory in developing mice.

## DISCUSSION

Infant humans and non-human infant animals can form event-based memories, but these memories fade with time. A leading hypothesis is that the neural circuits required for the conversion of recent into remote memory (or systems consolidation) are not yet fully mature^6,14,15,19,23^. Our results identify myelination of PrL circuits as a critical developmental milestone for the emergence of adult-like systems consolidation. Specifically, we provide causal evidence for a developmental pathway in which PrL neuronal activity drives BDNF–TrkB signaling in OPCs, promoting their differentiation into myelinating oligodendrocytes and the maturation of circuits necessary for adult-like memory persistence.

In developing sensory cortices, the maturation of oligodendrocytes and the onset of myelination marks the closure of critical periods and the emergence of adult-like perceptual function^55–58^. During these windows, activity-dependent BDNF–TrkB signaling promotes selective myelination of active circuits, refining conduction speed and coordinating spike timing across distributed networks^59^. Once established, myelin-associated inhibitory cues (e.g., Nogo-A, myelin-associated glycoprotein [MAG] and oligodendrocyte-myelin glycoprotein [OMgp]) limit structural remodeling and stabilize the mature circuit architecture via their interactions with Nogo receptors and PirB^58,60–62^. Our findings suggest the PrL undergoes a comparable developmental transition for memory: activity-driven BDNF–TrkB signaling in OPCs promotes oligodendrocyte maturation and myelination, enabling remote memory consolidation. Similar to studies of critical period plasticity in the developing cortex, we found that interventions that disrupt this pathway delay infantile amnesia offset, whereas enhancing this pathway leads to precocial emergence of adult-like memory persistence.

Systems consolidation depends on coordinated interactions between hippocampal and cortical networks, through which initially hippocampus-dependent representations are gradually reorganized and stabilized in distributed cortical circuits^24,25^. A defining feature of infantile amnesia is that the neural systems supporting this transformation are themselves immature early in life. In earlier work, we showed that elevated levels of hippocampal neurogenesis during infancy contribute to the weakening of hippocampal memory traces, and that reducing postnatal neurogenesis prolongs the persistence of infant-acquired memories^12^. However, even under conditions of reduced neurogenesis, these memories were eventually lost, indicating that preserving hippocampal traces alone is not sufficient for successful remote memory consolidation. Instead, the cortical circuits destined to support long-term storage must themselves reach a mature functional state in order for memories to be transformed from a recent, hippocampus-dependent form into a stable remote representation. Our findings identify developmental myelination of PrL circuits as a critical and rate-limiting step in this process.

Importantly, weak or incomplete systems consolidation does not necessarily imply that infant memories are completely lost. Optogenetic studies indicate that memories formed early in life can persist in a latent form and be recovered by direct reactivation of hippocampal engram neurons, even after they have become behaviorally inaccessible^13,63,64^. These findings indicate that early forgetting reflects, at least initially, a failure of natural retrieval rather than a true loss of the memory trace^65^. Within this framework, insufficient myelination of PrL circuits may impair the stabilization and cortical embedding of memory representations, resulting in impoverished or unstable access routes despite continued engram persistence. Accordingly, the emergence of adult-like memory persistence may reflect not only the stabilization of memory content in cortex, but also the maturation of the circuit infrastructure required for its long-term accessibility.

## METHODS

### Mice

Three mouse lines were used. Male and female C57BL/6NTa wild-type (WT) mice were used for all experiments unless otherwise specified. Mice were bred at the Hospital for Sick Children and group-housed on a 12-h light/dark cycle with food and water available ad libitum. All experiments were conducted during the light phase. Both primiparous and multiparous dams were used for breeding, and the date of birth was assigned as postnatal day (P) 0. Litter sizes ranged from 5 to 12 pups. Pre-weaning mice (≤ P20) remained in their breeding cages (identical to standard cages) with the dam for the duration of the experiments. Older mice (≥ P25) were weaned from the dam on P21 and subsequently group-housed with same-sex littermates in standard housing cages (2–5 mice per cage). Each experimental condition included mice derived from 2–8 independent litters, with no more than two same-sex littermates assigned to the same experimental condition.

TrkB_CKO_ mice (NG2-cre^ERTM^ × TrkB^fl/fl^) were generated using a tamoxifen-dependent recombinase (cre^ERTM^) expressed in NG2^+^ cells (NG2-cre^ERTM^, strain: 008538) crossed with a line carrying loxP sites flanking the Ntrk2 (TrkB) gene (B6N.129X1-Ntrk2^tm1Lfr^/Mmucd, strain: 000187-UCD) ^66^. Crossing NG2-cre^ERTM^ mice with homozygous floxed-TrkB mice produced hemizygous NG2 and heterozygous floxed-TrkB offspring, which were subsequently backcrossed to obtain hemizygous NG2 and homozygous floxed-TrkB mice (TrkB_CKO_).

Similarly, Myrf_CKO_ mice (NG2-cre^ERTM^ × Myrf^fl/fl^) were generated using the NG2-cre^ERTM^ line crossed with a strain containing loxP sites flanking the Myrf gene (B6-Myrf^tm1Barr^/J, strain: 010607) ^66,67^. Offspring were first produced as hemizygous NG2 and heterozygous floxed-Myrf, then backcrossed to generate hemizygous NG2 and homozygous floxed-Myrf mice (Myrf_CKO_).

### Drugs

#### DREADD agonist 21 (C21)

C21 dihydrochloride (Tocris, cat# 6422) was dissolved as a 10 mg/ml stock solution in dH₂O and stored at –20 °C. C21 was administered in two ways. First, the stock was thawed and diluted 1:10 in PBS for injections. The diluted solution was administered via i.p. injection (1.0 mg/kg) as described for specific experiments. Second, the stock was diluted 1:1250 in drinking water and provided as the sole water source for mice.

#### α2-antiplasmin (α2AP)

This inhibitor of the tPA–plasmin system (Sigma Aldrich, cat# SRP6313) was dissolved in PBS at 0.5 mg/ml and stored at –80 °C until use.

*Agonist anti-TrkB antibody (38B8).* This monoclonal antibody activates the TrkB receptor (Absolute Antibodies, cat# Ab03000-3.0). It was dissolved in PBS at 1 mg/ml and stored at 4 °C until use.

#### Recombinant Brain-Derived Neurotrophic Factor (BDNF)

Recombinant BDNF (Peprotech, cat# 450-02) was dissolved in PBS at 0.36 mg/ml and stored at –80 °C until use.

#### 4-hydroxy tamoxifen (4-OHT)

4-OHT (Toronto Research Chemicals, cat# T006000) was mixed with absolute ethanol (10 mg in 250 μl) and vortexed. The suspension was maintained at 50 °C with intermittent vortexing until the compound fully dissolved. Cremophor (250 μl) was then added in equal volume to generate a stock solution, which was stored at −20 °C until use. On the experimental day, this stock was diluted 1:4 with PBS. The final 4-OHT solution was delivered by intraperitoneal injection at 25 mg/kg.

#### 5-ethynyl-2′-deoxyuridine (EdU)

EdU is a thymidine analog that is incorporated into the DNA of cells undergoing division, enabling the identification of newly generated cells. EdU (Biosynth, NE08701, CAS: 61135-33-9) was prepared by dissolving 10 mg of the compound in 1 ml of 0.1 M PBS. Mice were treated for 4 days with single intraperitoneal injections per day (100 mg/kg). Later the collected tissues were processed by Click chemistry. Briefly, 50-µm sections were incubated in Tris-buffered saline (TBS) for 30 min. The Click reaction mix was then prepared by combining dH₂O, 1 M Tris buffer (pH 8.5), CuSO₄·5H₂O, and sulfo-Cyanine5 azide (Lumiprobe, cat# B3330, 10 mM) at a ratio of 599:100:200:1. Activation of the reaction was achieved by adding 1 M ascorbic acid to the mixture at a 1:9 ratio. Tissue sections were placed in the activated Click solution for 30 min and subsequently rinsed three times with PBS.

### Viruses

#### HSV-hM4Di-mCherry

Replication-defective HSV vectors were used to express the inhibitory DREADD hM4D_i_ in PrL neurons in both adult and developing mice. The virus was produced in-house by subcloning an hM4D_i_ construct (hSyn-hM4D(Gi)-mCherry; Addgene #50475) into an HSV-p1006 vector backbone. This construct drives expression of hM4Di and the fluorescent marker mCherry under the IE4/5 promoter.

#### AAV8-CaMKII-hM4Di-IRES-mCitrine

This virus was used to drives hM4Di expression in excitatory PrL neurons in developing mice. It was purchased premade (Addgene #50467-AAV8) and expresses hM4D_i_ and mCitrine under the CaMKII promoter.

### Surgery

Mice were pre-treated with atropine sulfate (0.1 mg/kg, i.p.), anesthetized with either chloral hydrate (400 mg/kg, i.p.), given meloxicam (4 mg/kg, s.c.) for analgesia, and positioned in stereotaxic frames. The scalp was incised and retracted, and holes were drilled above the PrL. Unless otherwise specified, viruses or drugs were injected bilaterally through a glass micropipette connected to a microsyringe (Hamilton) at 0.1 μl/min, with injectors left in place for an additional 5 min to ensure diffusion.

Coordinates and injection volumes for PrL were as follows. For P15 surgeries: AP +1.65 mm, ML ±0.35 mm, DV –1.7 mm from bregma. For older mice: AP +1.7 mm, ML ±0.35 mm, DV – 1.8 mm. Injection volumes were: 1.0 μl HSV; 0.7 μl AAV; 0.7 μl BDNF; 0.5 μl α2AP; 0.5 μl 38B8. Following bilateral injections, the scalp was sutured and Polysporin applied. Mice received 0.9% saline (0.5–1.0 ml, s.c.) and were placed on a heating pad to recover. Once recovered, P15 or P20 mice were reunited with the dam in a clean cage, and older mice (≥ P21) were returned to their home cages.

Surgical procedures for neonatal (P1) surgeries followed established protocols^68^. P1 pups were anesthetized via hypothermia and placed on a chilled metal plate. After scalp incision and retraction, a glass micropipette connected to a nanoliter injector (Nanoject III, Drummond Scientific) was used to penetrate the skull over the PrL (approximate coordinates AP 0.6 mm, ML ±0.2 mm from bregma). The pipette was lowered to DV –0.9 mm from the skull surface, and 100 nl of AAV was delivered over 2 min, with the pipette left in place for an additional 2 min. Wounds were sealed with Vetbond Tissue Adhesive (3M) and covered with Polysporin. Some pups received non-toxic ink tattoos for identification. The full procedure was completed within 10–12 min. Pups remained on a heating pad until mobile, and the full litter was returned to the dam once all surgeries were completed.

For all virus injection experiments, only mice exhibiting strong bilateral expression confined to the target region and visible in at least three brain sections were included in analyses.

### Contextual fear conditioning

#### Context configuration

The conditioning chamber (32 × 25 × 25 cm) had transparent acrylic panels on the front and top and aluminum walls along the sides and rear. Its floor was made of metal bars that delivered foot shocks. A camera positioned in front of the acrylic wall was used to record behavior.

#### Conditioning

On the conditioning day, mice were moved from their housing area to a quiet holding room and left there for 60 min. They were then brought individually to the conditioning room and placed into the chamber. Three shocks (0.5 mA, 2 s) occurred at 120, 150, and 180 s. All mice were removed from the chamber exactly 240 s after being placed inside. Following conditioning, they remained in the holding room for an additional 60 min before being returned to their home cages.

#### Retrieval test

Mice were brought to the holding room 60 min before testing. Mice were tested in the original conditioning environment (4 min test duration). In these tests, mouse behavior was monitored via a front view camera (30 frames/s), and freezing was manually tracked with the experimenter blind to the conditions. The percentage of time spent freezing time across the entire testing period was used for statistical analyses^27^.

### Histology

#### On slide tissue preparation

For the TrkB/pTrkB staining experiments, mice were decapitated, the brain was dissected and flash frozen on dry ice. Next, the frozen brains were coronally sectioned on a cryostat (Leica CM1850) at 20 μm thickness. The sections were transferred to glass slide and fixed for 10 min in ice cold 100% ethanol. The slides were stored in -80 °C or immediately used for immunohistochemistry.

#### Perfusion and free-floating tissue preparation

For all other experiments, mice were transcardially perfused with chilled PBS followed by 4% PFA. Brains were fixed in PFA overnight at 4 °C and transferred to 30% sucrose for at least 48 h. PBS and PFA volumes were adjusted based on mouse age. Brains were coronally sectioned on a cryostat (Leica CM1850) into 50 μm sections. Sections for immunohistochemistry were stored in 0.1% NaN₃ until staining.

#### Immunohistochemistry

Immunofluorescence was performed using standard procedures. For experiments requiring quantification of cell numbers or fluorescence intensity, all sections were stained simultaneously using identical antibody solutions. For free-floating sections, a 24-well plate was used and for on slide staining the solution was directly applied on the slide. Samples were blocked for 2 h at room temperature in PBS containing 4% normal goat serum (NGS) or normal donkey serum (NDS) and 0.5% Triton-X. Sections were then incubated for 72 h at 4 °C with primary antibodies diluted in fresh blocking solution. Used primary antibodies include rabbit anti-c-Fos (1:1000, Synaptic Systems, cat# 226003), chicken anti-GFP (1:1000, Aves, cat# GFP-1010), rabbit anti-RFP (1:1000, Rockland Immunochemicals, cat# 600-401-379), rabbit anti-CaMK2ɑ (1:200, GeneTex, cat#GTX127939), chicken anti-Gad67 (1:200, Aves cat#GAD0508), mouse anti-MBP (1:200, Cell Signaling #83683), mouse anti-CC1 (1:250, Sigma, cat#OP80), rabbit anti-Olig2 (1:250, Sigma, cat#AB9610), Goat anti-pdgfrα (1:250, R&D systems cat#AF1062), mouse anti-TrkB (1:250, Absolute Antibodies,cat#Ab03000-3.0), mouse anti-pTrkB (1:250, Sigma, cat#ABN1381). After three 5-min PBS washes, sections were incubated for 24 h at 4 °C with secondary antibodies and 0.1% Tween20 in PBS. Used secondary antibodies include goat anti-chicken Alexa Fluor 488 (1:500, Invitrogen, cat# A-11039), goat anti-mouse Alexa Fluor 568 (1:500, Invitrogen, cat# A-21124), goat anti-rabbit Alexa Fluor 647 (1:500, Invitrogen, cat#A32733), donkey anti-mouse Alexa Fluor 568 (1:500, Invitrogen, cat#A-10037), donkey anti-rabbit Alexa Fluor 647 (1:500, Invitrogen, cat#A-31573), donkey anti-mouse Alexa Fluor 647 (1:500, Invitrogen, cat#A-31571), donkey anti-goat Alexa Fluor 488 (1:500, Invitrogen, cat#A-11055). Sections were washed, counterstained with DAPI (1:5000) when required, the free-floating sections were mounted on gel-coated slides, and coverslipped with Permafluor mounting medium (ThermoFisher Scientific, cat# TA-030-FM).

#### Imaging

Images were acquired using a confocal laser scanning microscope (Zeiss LSM 710 or Zeiss LSM 880) using a 20× objective. Z-stacks were collected for quantification. Identical imaging settings (laser power, detector gain, pinhole size, filter settings) were used for all samples within a given experiment, and parameters were set using a control mouse section. In experiments, with different age groups, P60 samples served as controls.

#### Quantification

Cells expressing the marker of interest were counted across the full cross-sectional area of each region of interest. All images were processed identically, including z-stack projection, background filtering, and thresholding to generate countable masks. With the exception of DAPI^+^ cell counts, all cells were counted manually in Fiji (NIH). DAPI^+^ nuclei were counted using Fiji’s particle counter (size: 6–20 μm), and automatic counts were validated by manual scoring of randomly selected images. Counts from replicate images for each mouse were pooled to generate the final value used for analysis.

### Western Blot

The mice were rapidly decapitated, and PrL was dissected and flash frozen on dry ice. The tissues were sonicated in homogenization buffer (50 mM Tris-HCl pH 7.5, 0.25 M sucrose, 25 mM KCl, 5 mM MgCl2) supplemented with protease inhibitor cocktail (BioShop, cat# PIC002). Homogenized samples were centrifuged at 14,000 rpm for 15 min at 4 °C and the protein concentration of the supernatant was determined by Pierce BCA Protein Assay (ThermoFisher Scientific, cat# 23227). Samples were diluted and supplemented with SDS sample buffer (50 mM Tris-HCl pH 6.8, 2% SDS, 1% β-mercaptoethanol, 5% glycerol, bromophenol blue). Protein samples (20 µg) were separated by electrophoresis on 4-15% mini-PROTEAN TGX precast gels (Bio-Rad, cat# 456-1083) and transferred to PVDF membranes (Bio-Rad, cat# 162-0177). Membranes were blocked with 5% skimmed milk in Tris-buffered saline with 0.1% Tween-20 (TBS-T) for 1 h at room temperature. Membranes were incubated in primary antibodies diluted in 5% skimmed milk in TBST: rabbit anti-BDNF (1:1000, Abcam cat# ab108319), mouse anti-b-actin (1:2000, Cell Signaling Technology, cat#3700S), chicken anti-GAPDH (1:5000, Millipore, AB2302) overnight at 4°C. Membranes were incubated with horseradish peroxidase (HRP)-conjugated secondary antibodies diluted in 5% skimmed milk in TBST: goat anti-rabbit IgG (1:50,000, Sigma Aldrich cat# A0545), horse anti-mouse IgG (1:50,000, Cell Signaling Technology, cat# 7076) for 2 h at room temperature. Protein bands were visualized with Amersham ECL Prime Western Blotting Detection Reagent (Cytiva, cat# RPN2236) and chemiluminescence was imaged with a ChemiDoc XRS+ System (Bio-Rad). Band intensities were quantified using Image Lab 6.1 software (Bio-Rad). The intensities of the proteins of interest were normalized to the intensity of the loading control, β-actin and the data within each set were normalized to the maximum BDNF:β-actin (or BDNF:GAPDH) ratio which was at P20 for every blot.

### 10x Genomics Xenium

#### Tissue preparation and data processing

Single-cell spatial transcriptomics was performed using the 10x Genomics Xenium *In Situ* platform on brain tissue sections^69^. Brains were dissected from C57BL/6 mice at P15, P20, P25 and P60. Unfixed samples were immediately embedded in OCT compound (Tissue-Tek) on dry ice, stored at -70 C, and cryosectioned coronally onto Xenium slides (chemistry v1) to produce 10 μm sections. The slide was then stored at -70 °C until subsequent steps were performed. The tissue was processed using the Xenium workflow for fresh frozen tissue, first undergoing fixation and permeabilization. In brief, sections were incubated at 37 °C for 1 min, then fixed in 4% PFA (Electron Microscopy Sciences) for 30 min at room temperature. Afterwards, sections were washed in RNase-free 1X PBS (Thermo Fisher) for 1 min and underwent a series of permeabilization steps and PBS washes. These steps included a 2 min incubation in 1% SDS (Millipore Sigma) and a 60 min incubation in chilled 70% methanol (Sigma). Following this, sections were washed twice for 5 min in PBS and placed in PBS-Tween (PBS-T) containing 0.05% Tween-20 (Thermo Fisher). Sections were then incubated in a probe solution for 16-24 h at 50 °C for hybridization. The probe solution contained a set of 347 gene-targeting probes (see Xenium panel design) in TE buffer (Fisher). Afterwards, sections underwent a series of washes in PBS and were incubated at 37 °C for 30 min in Xenium post-hybridization wash buffer. Sections were next washed with a series of PBS-T washes and incubated in Xenium ligation enzymes for 2 h at 37 °C for probe ligation. After several PBS-T washes, probe amplification was carried out by incubating sections in Xenium amplification enzyme for 2 h at 30 °C. Sections were then washed twice in TE buffer and stored overnight at 4 °C. Autofluorescence quenching and nuclei staining steps were carried out next. Sections were washed in PBS, incubated for 10 min in a reducing agent, washed in 70% and 100% ethanol (Sigma), and incubated in Xenium autofluorescence solution for 10 min. Following this, sections were washed three times in 100% ethanol, dried at 37 °C for 5 min, and rehydrated via subsequent washes in PBS and PBS-T. Sections were then incubated in a nuclear staining buffer for 1 min and washed four times in PBS-T. Following this, the sample was loaded into the Xenium analyzer instrument (software v1.7.6.0) and subjected to multiple cycles of reagent application, probe hybridization, imaging, and probe removal. Initial pre-processing of captured Z-stack images was performed using Xenium onboard analysis pipeline (v1.7.1.0) ^69,70^. In brief, our custom Xenium codebook was used to decode puncta into transcripts, with each codeword assigned to genes in the gene panel. Quality scores (Q-Scores) were assigned to transcripts based on maximum likelihood codewords, and negative controls were used to ensure accurate calibration. On-instrument nuclei segmentation was performed based on DAPI morphology, with each segmented nuclei assigned a cell ID. All data were subsequently processed through Xenium Ranger (v1.7.1.1) (via resegment function) with settings as recommended by the manufacturer to assign transcripts to the closest nucleus within a maximum distance of 15 μm. Following pre-processing, standardized Xenium output files were exported for downstream analyses.

#### Xenium panel design

Single-cell spatial transcriptomics was performed using a panel targeting 347 mRNA transcripts. 247 of these probes were from the mouse brain panel designed by 10x Genomics (10xgenomics.com/products/xenium-panels). A 100-probe custom add-on panel was generated to target the following genes: *Dll3*, *Nfix*, *Slc38a1*, *Tek*, *Bmx*, *Agt*, *Hbegf*, *Eps8*, *Gjb6*, *Lcat, Icam2*, *Vcam1*, *Cxcl12*, *Cxcr4*, *Jag1*, *Notch1*, *Clic6*, *Kcnj13*, *Aldoc*, *Hes5*, *Mt3*, *Slc1a3*, *Rgcc*, *Ttyh1*, *Gsta4*, *Grasp*, *Cdh5*, *Dll4*, *Tie1*, *Foxj1*, *Crybb1*, *Ecscr*, *Epas1*, *Npas1*, *Nrarp*, *Pcp4*, *Ptprn*, *Hes1*, *Clu*, *Il1r1*, *Enpp6*, *Fyn*, *Angpt2, Vegfa*, *Il6*, *Il1b*, *Osm*, *Il6st*, *Lifr, Eomes*, *Neurod1*, *AB124611*, *Cybb*, *Lgals3*, *Wfdc17*, *Cd14*, *Ly96*, *Myd88*, *Nfkb1*, *Tlr4*, *Cebpd*, *Cfn*, *Icam1*, *Lcn2*, *Bmp7*, *Clec11a*, *Lama1*, *Slc22a6*, *Aif1*, *Tmem119*, *Dlx1*, *Dlx2*, *Dlx5*, *Sp8*, *Sp9*, *P22ry12*, *Mdfi, Nes*, *Shroom3*, *Tfap2c*, *Thbs4*, *Tspan18*, *Veph1*, *Vnn1*, *Fam107a*, *Prex2*, *Mag*, *Mbp*, *Olig1*, *Olig2*, *Gpnmb*, *Igf1r*, *Ascl1*, *Egfr, Gsx2*, *Mki67*, *Flt1*, *Nr4a1*, *Emx1*, and *Nkx2-1*.

#### 10x Xenium Data Analysis

Xenium data files, generated by the onboard analysis pipeline, were first visualized in Xenium Explorer (v1.2.0). Regions of interest (ROIs) were defined for each section, a total of 12 sections were analyzed across 12 mice which encompassed the cells in the prelimbic cortex (PrL) area. The cell IDs corresponding to each ROI were then imported into R alongside transcript count, feature, and coordinate data via the Seurat package (v5.0.1) to generate Seurat objects containing expression data corresponding to the regions of interest per tissue section. Low-quality cells were identified based on the mean number of detected genes and total transcript counts, and cells greater than +/-2.5 standard deviations from the mean were filtered out. Filtered data were transformed using SCTransform and dimensionality reduction was performed using PCA based on highly variable genes. This was subsequently used to construct a shared nearest neighbor (SNN) graph using the FindNeighbors function in Seurat. The data was then clustered using the FindClusters function across a range of resolutions (0.4 to 2.4) and visualized in two-dimensional space using UMAP embeddings. Spatial plots and UMAPs, generated using the ImageDimPlot, FeaturePlots, and ImageFeaturePlots functions, were used to investigate gene expression and interpret cell clustering patterns. Each section was initially analyzed separately. Afterwards, data corresponding to different sections were merged using the Seurat SelectIntegrationFeatures, FindIntegrationAnchors, and IntegrateData functions. Where relevant for improved annotation, the Seurat subset function was used to subset out clusters of interest. Normalization, transformation, PCA, and clustering was performed on each produced merged dataset and/or subset.

We used the following well-defined markers genes to identify and annotate cell types: For cortical excitatory neurons, *Satb2*, *Neurod6*, *Slc17a7*, *Slc17a6*, *Fezf2*, *Bcl11b*, *Cux2*, *Neurod1*; for PV INs *Pvalb*; for somatostatin interneurons *Sst*; for other INs *Dlx5*, *Gad2*, *Sp9*, *Gad1*, *Dlx1*, *Dlx2*, *Sp8*; for VIP interneurons *Vip*; for OPCs *Pdgfra*, *Cspg4*; for the oligodendrocyte lineage, *Olig1*, *Sox10*; for immature oligodendrocytes, *Gpr17*, *Enpp6*, *Fyn*; for mature oligodendrocytes, *Mbp*, *Mag*, *Opalin*; for microglia, *Cd68*, *Aif1*, *Tmem119* ; for endothelial, *Adgrl4*, *Pecam1*, *Car4*; for pericytes *Inpp4b*, *Sntb1*, *Carmn*, *Ano1*, *Acta2*, *Cspg4*; for meninges *Bmp7*, *Clec11a*, *Gjb2*, *Slc22a6*, *Lama1*, and for cortical astrocytes *Lcat*, *Aqp4*, *Agt*, *Gjb6*, *Hbegf*, *Eps8*, *Gfap*. Annotation of single cell spatial transcriptomics data was also informed by expected cell location. Specifically, for L2/3, L5, L6a and L6b cortical excitatory neurons, the spatial location of cell clusters on each section was used for annotation.

Each dataset was analyzed by choosing the most conservative resolution; resolution was only increased to interrogate cell heterogeneity or if two cell populations of known identity, based on the expression of canonical markers, were not separating into distinct clusters as expected. For the full dataset containing all cell types from all sections, clusters were assigned at a resolution of 0.8 (28 clusters identified). For the subset containing a selection of oligodendrogenic cell types, clusters were assigned at a resolution of 0.4 (4 clusters identified).

#### Differentially expressed genes

To investigate transcriptional differences across the developmental period of interest, differential gene expression analysis was performed. This analysis was performed using the Seurat FindMarkers function on the juvenile ages (i.e., P15, P20, P25) versus the adult age (i.e., P60) within the oligodendrocyte-lineage subset. Differentially enriched gene lists were refined by including only genes with a Bonferroni adjusted p-value <0.05 (Adj. p-value) and ≥0.25 log2 fold-change in average expression. The upregulated differentially expressed genes (DEGs), (corresponding to juvenile enriched genes) and downregulated DEGs (corresponding to adult enriched genes) were evaluated separately in Gene Ontology (GO) enrichment analysis. The GO enrichment was preformed using g:Profiler^71^ (version e113_eg59_p19_f6a03c19). The g:Profiler results were filtered for molecular functions and biological process and the select top terms were presented in Fig. 2m,n. Full lists are available in Tables S4 and S5.

### In vivo bioluminescence imaging

The generation of *Bdnf-Luc* mice has been described previously^72^. Bioluminescence imaging was performed using an IVIS in vivo imaging system (PerkinElmer, Boston, MA, USA) as reported previously^73^ Briefly, the hair on the head of Bdnf-Luc mice was shaved on postnatal day 14 (P14), one day before the start of imaging. AkaLumine hydrochloride (Kurogane Kasei Co., Ltd., Aichi, Japan), a synthetic luciferase substrate, was administered intraperitoneally at a dose of 50 mg/kg body weight under 3% isoflurane anesthesia. Imaging was initiated 3 min after substrate injection and repeated every 2 min to capture the peak intensity of the bioluminescence signals.

### Transmission electron microscopy

Brain regions of interest were dissected using a brain matrix and razor blades, then immersed in a fixative containing 2% PFA and 2.5% glutaraldehyde. Tissue remained in fixative for up to one week at 4 °C. After thorough rinsing, samples were post-fixed with 1% osmium tetroxide, dehydrated through an ethanol series, and embedded in Quetol–Spurr resin, polymerized overnight at 65 °C. Ultrathin sections (90 nm) were generated on a Leica EM UC7 ultramicrotome and subsequently stained with uranyl acetate followed by lead citrate. Imaging was carried out using a Tecnai 20 TEM, and quantification was conducted in FIJI (National Institutes of Health). All image collection and analyses were performed with experimenters blinded to group identity. For g-ratio measurements, axon diameter was taken at its narrowest point, and the myelinated fiber diameter was measured inclusive of the myelin sheath; the g-ratio was calculated as the ratio of axonal diameter to total fiber diameter.

### Statistical Analyses

The sample size for the experimental groups was determined based on previous publications^12,30,68,74^ Mice were pseudo-randomly assigned to different groups with roughly equal numbers of male and female mice. Data were analyzed using unpaired *t*-test, one-way or factorial analysis of variance (ANOVA) with repeated measures when appropriate. When appropriate, ANOVAs were followed by either Tukey or Sidack’s post-hoc comparisons. Distinct cohorts of mice were used for all behavior experiments. For histological analysis, whenever the conditions were identical, the tissue from the same animal is used for multiple different staining combinations (e.g., two sets of tissue from the same animal are used for staining Fos marker and for oligodendrogenic markers). Except for the transcriptome analysis, GraphPad Prism (version 8.0.1) was used for all statistical analyses and graphs.

## Acknowledgements

We thank A. DeCristofaro, D. Lin and M. Yamamato for technical assistance. This work was funded by a Natural Sciences and Engineering Research Council of Canada (RGPIN-2022-03520) grant to PWF.

## Author contributions

Conceptualization: AG, SAJ, PWF. Methodology: AG, MF, SK, AR, AIR. Data collection: AG, AIR, AR, MLdS, BS, YL, MF, SK. Data collection (transcriptomics): BSW, AW. Data analysis: AG, BS, YL, MF, SK. Data analysis (transcriptomics): AG, DK. Visualization: AG, JW, DK. Funding acquisition: PWF, SAJ, FDM, DRK. Writing – original draft: AG, PWF. Writing – review & editing: AG, AIR, FDM, DRK, SAJ, PWF.

## Competing interests

The authors declare no competing financial interests.

## Correspondence and requests for materials

Further information and requests for resources and reagents should be directed to and will be fulfilled by Paul Frankland and Sheena Josselyn. The transcriptomic data used in the analysis are available at https://www.ncbi.nlm.nih.gov/geo/query/acc.cgi?acc=GSE312677. (Reviewer token: ajqhyiistruvvqb)

---

**Extended Data Fig. 1.**
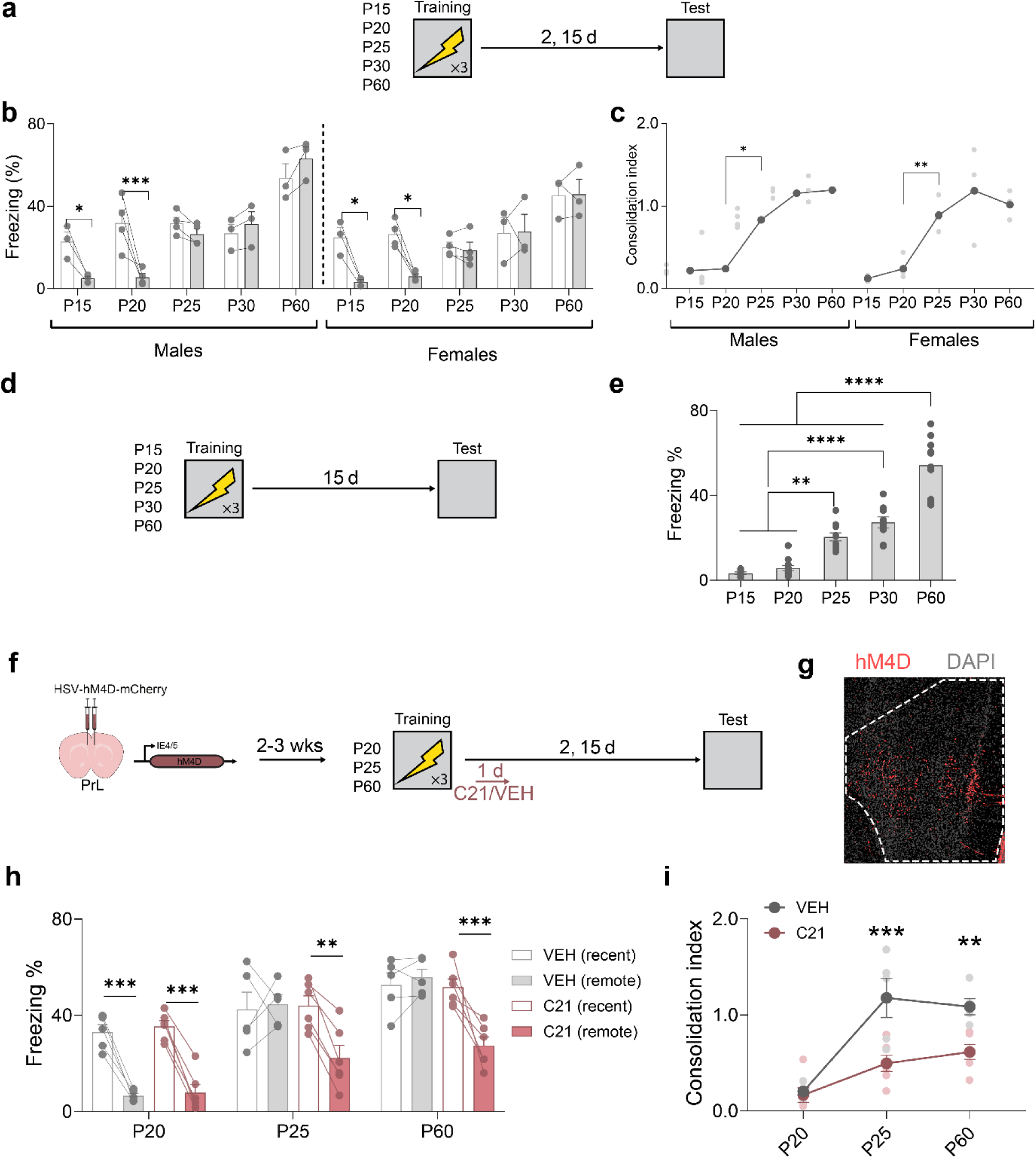
Characterizing infantile amnesia in mice. **a**, Mice were conditioned at different ages and tested 2 and 15 days later. **b**, Freezing at recent and remote delays in male and female mice. Successful remote memory consolidation emerged at P25 in both sexes (ANOVA, Age × Sex × Time-point interaction: *F*_4,24_ = 0.80, *P* = 0.54; main effect of Age: *F*_4,24_ = 26.35, *P* < 0.0001; main effect of Sex: *F*_1,24_ = 4.48, *P* < 0.05; main effect of Delay: *F*_1,24_ = 25.10, *P* < 0.0001; Age × Sex interaction: *F*_4,24_ = 0.99, *P* = 0.43; Age × Time-point interaction: *F*_4,24_ = 14.18, *P* < 0.0001; Sex × Time-point interaction: *F*_1,24_ = 0.18, *P* = 0.68). **c**, Consolidation indices showed the same age-dependent pattern in males and females (ANOVA, Age × Sex interaction: *F*_4,24_ = 0.29, *P* = 0.88). **d**, To rule out extinction as a confound, an independent cohort was trained at different ages and tested only at the remote delay (15 days). **e**, Only mice trained ≥P25 froze at the remote test (ANOVA, effect of Age: *F*_4,41_ = 56.04, *P* < 0.0001). **f**, Viral expression of hM4D_i_ in PrL neurons permitted C21-induced silencing for 24 h following training. **g**, Representative PrL hM4D_i_ expression (Scale bar, 200 µm). **h**, Freezing at recent and remote delays following post-training PrL silencing (ANOVA, Age × Inhibition × Time-point [Retention] interaction: *F*_2,27_ = 4.31, *P* < 0.05). **i**, Compared to VEH-treated controls, C21 impaired consolidation in ≥P25 mice (ANOVA, Age × main effect of Inhibition: *F*_1,27_ = 5.07, *P* < 0.05). * *P* < 0.05; ** *P* < 0.01; *** *P* < 0.001, **** *P* < 0.0001.

**Extended Data Fig. 2.**
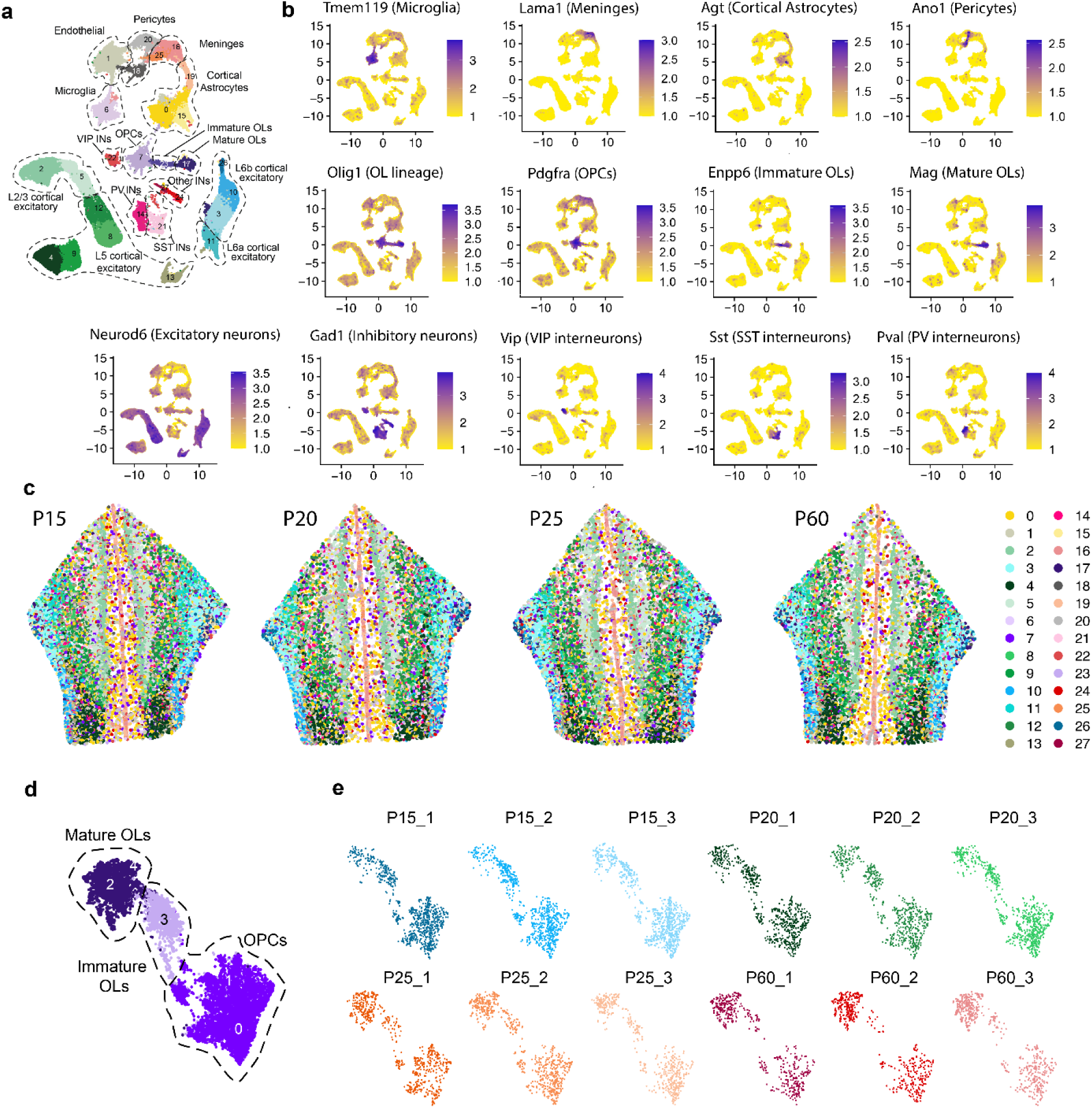
Spatial transcriptomic analysis of developing PrL. **a**, UMAP visualization of merged transcriptomes. **b**, Expression of cell-type specific marker genes. **c**, transcriptomes color coded for each cluster identified in (**a**). **d**, UMAP visualization of subsetted oligodendrogenic lineage cells. **e**, Original identity of cells in the oligodendrocyte UMAP. In UMAPs, numbers indicate clusters.

**Extended Data Fig. 3.**
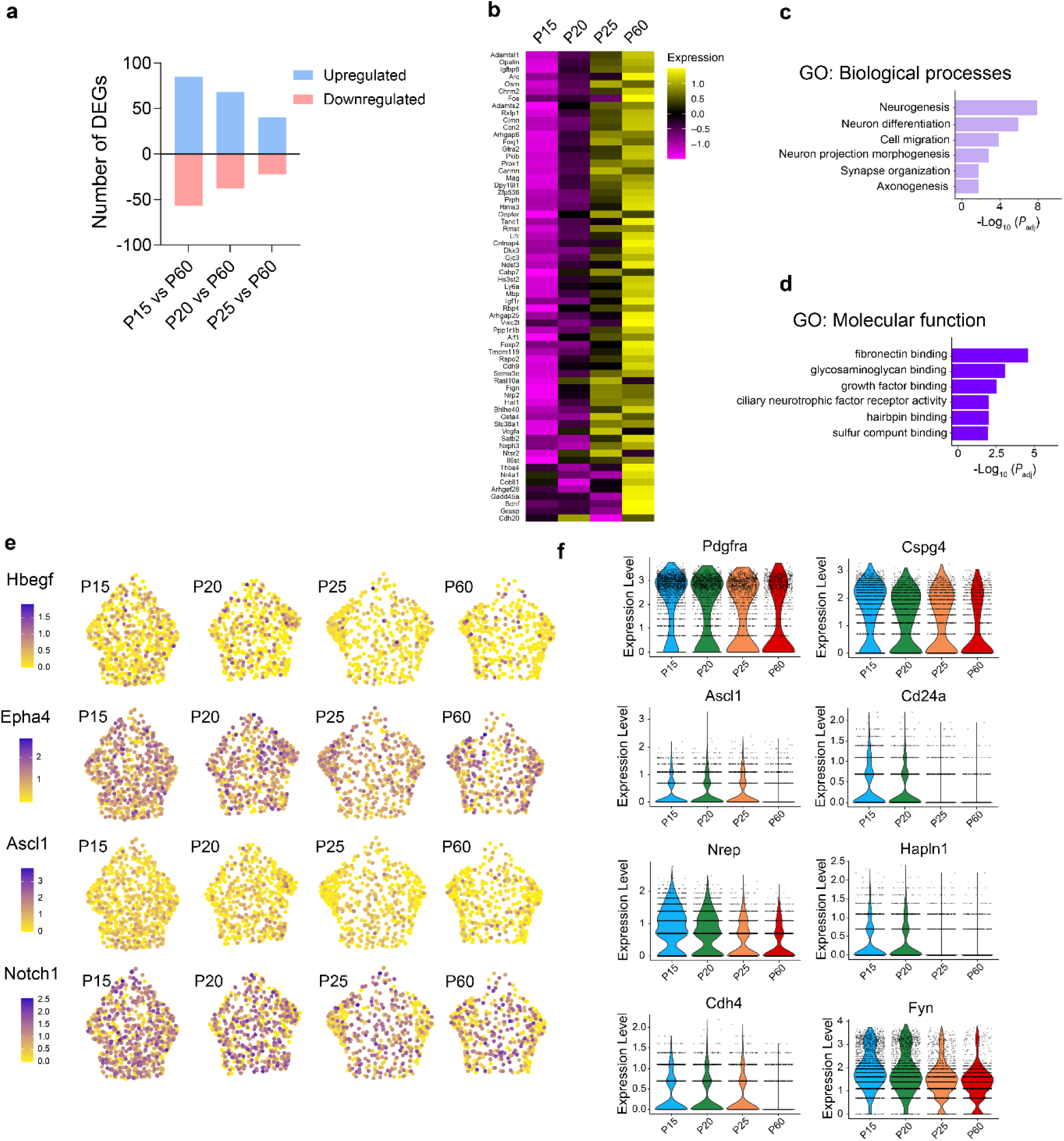
Differential gene expression analysis for PrL oligodendrogenic cells across the development. **a**, Number of down-regulated and upregulated DEGs across developmental period. **b**, Average expression of cell-type specific marker genes. **c**,**d**, GO terms enriched among upregulated genes, enriched at adult age, (molecular functions; biological processes). **e**, Spatial plots of cell clusters in sample sections of PrL across development. **f**, Violin plots of selected developmentally regulated genes (Pdgfra, Cspg4, Ascl1, Cd24a, Nrep, Hapln1, Cdh4, Fyn).

**Extended Data Fig. 4.**
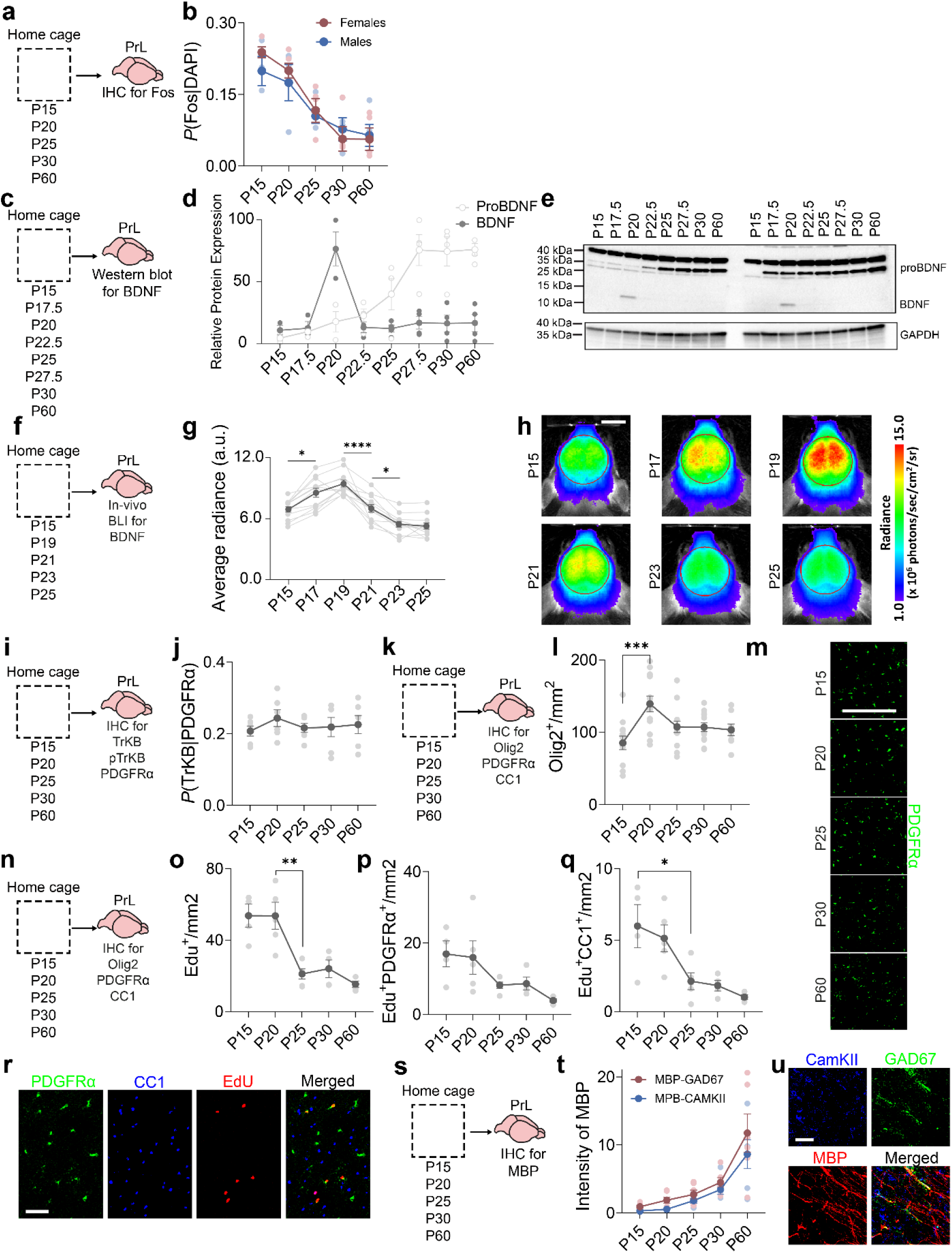
Home cage measures of activity, BDNF-TrkB signalling and oligodendrogenesis/myelination across development. **a**, Fos staining in PrL of home-cage mice was assessed across developmental ages. **b**, Fos expression decreased with age and did not differ by sex (ANOVA, Age × Sex interaction: *F*_4,28_ = 0.46, *P* = 0.77; main effect of Age: *F*_4,24_ = 15.80, *P* < 0.0001; main effect of Sex: *F*_1,28_ = 0.36, *P* = 0.55). **c**, Western blot for BDNF and proBDNF for PrL samples from home cage mice at P15–P30 and P60. **d**, proBDNF increased with age (effect of Age: *F*_7,21_ = 11.01, *P* < 0.0001); mature BDNF peaked at P20 (effect of Age: *F*_7,21_ = 10.06, *P* < 0.0001). **e**, Representative blot for BDNF, proBDNF, and GAPDH. **f**, *In vivo* BDNF mRNA expression measured in *BDNF-Luc* mice using bioluminescence imaging. **g**, BDNF mRNA peaked at P19 (ANOVA, main effect of Age: *F*_5,66_ = 23.33, *P* < 0.0001). **h**, Representative bioluminescence images. Scale bar: 5 mm. **i**, PrL tissue was stained for TrkB, pTrkB, and PDGFRα in home cage mice from P15-P60. **j**, ∼20% of PDGFRα^+^ OPCs expressed TrkB, and this was stable across ages (ANOVA, main effect of Age: *F*_4,25_ = 0.41, *P* = 0.80). **k**, PrL was stained for oligodendrocyte lineage markers in home cage mice from P15-P60. **l**, Olig2^+^ cells peaked at P20 (ANOVA, main effect of Age: *F*_4,53_ = 5.24, *P* < 0.01). **m**, Representative PDGFRα staining. Scale bar: 200 µm. **n**, EdU was administered for 4 days; one day following EdU treatment brains were collected at P15, P20, P25, P30, or P60. **o**, Total EdU^+^ cells peaked at ≤P20 (ANOVA, main effect of Age: *F*_4,17_ = 12.10, *P* < 0.0001). **p**, EdU^+^ PDGFRα+ OPCs decreased with age (ANOVA, main effect of Age: *F*_4,17_ = 3.45, *P* < 0.05). **q**, EdU^+^CC1^+^ oligodendrocytes also decreased (ANOVA, main effect of Age: *F*_4,17_ = 6.50, *P* < 0.01). **r**, Representative EdU/PDGFRα/CC1 image. Scale bar: 50 µm. **s**, PrL stained for MBP, CaMKII, and GAD67 in home cage mice from P15-P60. **t**, MBP–CaMKII and MBP–GAD67 overlap increased with age, indicating myelination of both excitatory and inhibitory neurons (ANOVA, Age × Neuron type interaction: *F*_4,25_ = 0.46, *P* = 0.76; main effect of Age: *F*_4,25_ = 16.67, *P* < 0.0001; main effect of Neuron type: *F*_1,25_ = 1.74, *P* = 0.09). **u**, Representative MBP/CaMKII/GAD67 image. Scale bar: 50 µm.

**Extended Data Fig. 5.**
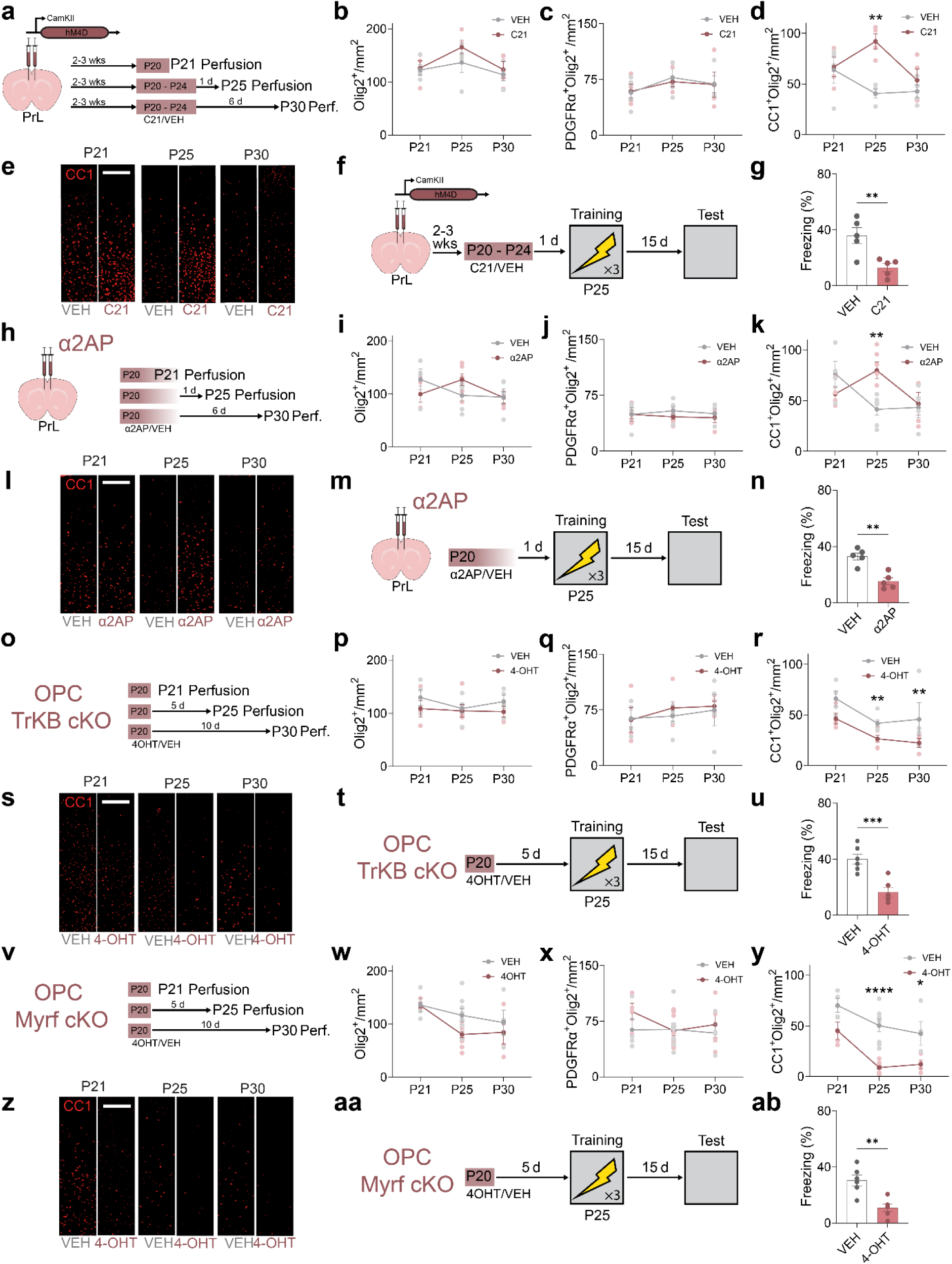
Inhibiting PrL activity, BDNF signaling, or TrkB activation delays oligodendrogenesis. **a**, hM4D_i_ was expressed in PrL, and mice received C21 or VEH for 5 days starting from P20; PrL tissue was collected 1, 5, or 10 days after start of the treatment. **b**,**c**, Olig2^+^ (Age × Inhibition interaction: *F*_2,18_ = 0.43, *P* = 0.66; main effect of Age: *F*_2,18_ = 3.03, *P* = 0.07; main effect of Inhibition: *F*_1,18_ = 1.50, *P* = 0.23) and PDGFRα^+^ (Age × Inhibition interaction: *F*_2,18_ = 0.05, *P* = 0.95; main effect of Age: *F*_2,18_ = 1.04, *P* = 0.37; main effect of Inhibition: *F*_1,18_ = 0.03, *P* = 0.87) cell numbers were unaffected. **d**, CC1^+^ mature oligodendrocytes was elevated 5 days of C21-induced inhibition and 1 day of recovery (i.e., at P25). However, with extended recovery time (6 days) the density of mature oligodendrocytes was decreased (i.e., at P30), indicating a delayed peak in oligodendrogenesis (ANOVA, Age × Inhibition interaction: *F*_2,18_ = 4.02, *P* < 0.05; main effect of Age: *F*_2,18_ = 2.47, *P* = 0.11; main effect of Inhibition: *F*_1,18_ = 1.74, *P* < 0.01). **e**, Representative CC1 staining. Scale bar: 200 µm (F, G) PrL-silenced mice conditioned at P25 showed reduced freezing in a remote test (unpaired t test, *t* = 3.59, *P* < 0.05). **h**, α2AP was injected into PrL at P20; tissue collected 1, 5, or 10 days later. **i**,**j**, Olig2^+^ (Age × α2AP interaction: *F*_2,22_ = 2.37, *P* = 0.12; main effect of Age: *F*_2,22_ = 1.21, *P* = 0.32; main effect of α2AP: *F*_1,22_ = 0.00, *P* = 0.97) and PDGFRα^+^ (Age × α2AP interaction: *F*_2,22_ = 0.22, *P* = 0.80; main effect of Age: *F*_2,22_ = 0.08, *P* = 0.92; main effect of α2AP: *F*_1,22_ = 0.84, *P* = 0.37) cells were unchanged. **k**, CC1^+^ mature oligodendrocytes elevated in C21 group after 5 days of inhibition and 1 day of recovery (i.e., P25). However, with extended recovery time (6 days) the density of mature oligodendrocytes was decreased by P30, indicating a delayed peak in oligodendrogenesis (ANOVA, Age × α2AP interaction: *F*_2,22_ = 5.75, *P* < 0.01; main effect of Age: *F*_2,22_ = 2.67, *P* = 0.09; main effect of α2AP: *F*_1,22_ = 0.97, *P* = 0.34). **l**, Representative CC1 staining. Scale bar: 200 µm. **m**,**n**, α2AP-treated mice conditioned at P25 showed reduced freezing in a remote test (unpaired t test, *t* = 5.04, *P* < 0.01). **o**, TrkB_CKO_ mice received 4-OHT at P20; PrL collected 1, 5, or 10 days later. **p**,**q**, Olig2^+^ (Age × TrkB-KO interaction: *F*_2,22_ = 0.32, *P* = 0.73; main effect of Age: *F*_2,22_ = 0.54, *P* = 0.59; main effect of TrkB-KO: *F*_1,22_ = 2.18, *P* = 0.15) and PDGFRα^+^ (Age × TrkB-KO interaction: *F*_2,22_ = 0.15, *P* = 0.86; main effect of Age: *F*_2,22_ = 0.69, *P* = 0.51; main effect of TrkB-KO: *F*_1,22_ = 0.23, *P* = 0.64) cells were unchanged. **r**, CC1^+^ cells decreased at P25 and P30, indicating sustained impairment of oligodendrogenesis (ANOVA, Age × TrkB-KO interaction: *F*_2,22_ = 0.16, *P* = 0.85; main effect of Age: *F*_2,22_ = 6.00, *P* < 0.01; main effect of TrkB-KO: *F*_1,22_ = 10.92, *P* < 0.01). **s**, Representative CC1 staining. Scale bar: 200 µm. **t**,**u**, TrkB-deleted mice conditioned at P25 showed reduced freezing in a remote test (unpaired t test, *t* = 4.83, *P* < 0.001). **v**, Myrf_CKO_ mice received 4-OHT at P20; PrL collected 1, 5, or 10 days later. **w**,**x**, Olig2^+^ (Age × Myrf-KO interaction: *F*_2,29_ = 0.16, *P* = 0.85; main effect of Age: *F*_2,29_ = 2.69, *P* = 0.08; main effect of Myrf-KO: *F*_1,29_ = 0.83, *P* = 0.37) and PDGFRα^+^ (Age × Myrf-KO interaction: *F*_2,29_ = 0.78, *P* = 0.47; main effect of Age: *F*_2,29_ = 0.72, *P* = 0.50; main effect of Myrf-KO: *F*_1,29_ = 1.52, *P* = 0.23) cells were unchanged. **y**, CC1^+^ cells decreased at P25 and P30, indicating sustained impairment of oligodendrogenesis (ANOVA, Age × Myrf-KO interaction: *F*_2,29_ = 1.12, *P* = 0.34; main effect of Age: *F*_2,29_ = 13.12, *P* < 0.0001; main effect of Myrf-KO: *F*_1,29_ = 39.21, *P* < 0.0001). **z**, Representative CC1 staining. Scale bar: 200 µm. **aa**,**bb**, Myrf-deleted mice conditioned at P25 showed reduced freezing in a remote test (unpaired t test, *t* = 4.21, *P* < 0.01).

**Extended Data Fig. 6.**
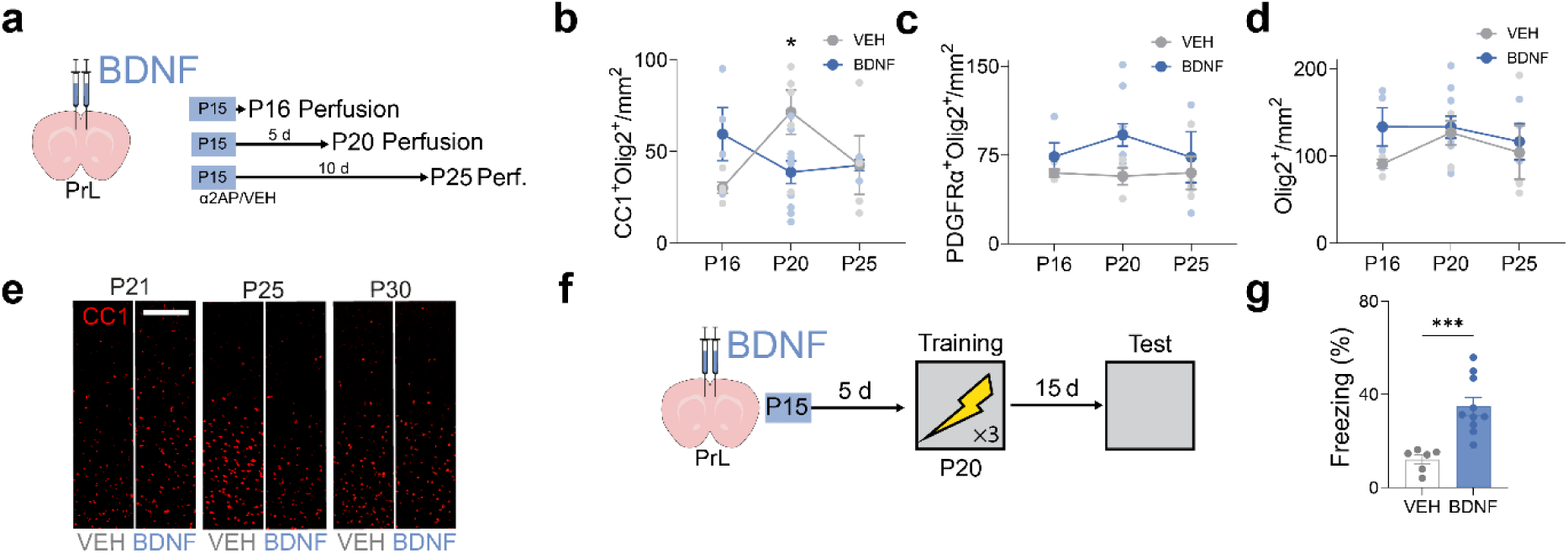
BDNF advances the timing of oligodendrogenesis in PrL. **a**, Mice received PrL BDNF microinjection at P15 and were perfused 1, 5, or 10 days later for oligodendrocyte marker analysis. **b**, Olig2⁺ cell numbers were unchanged (ANOVA, Age × BDNF interaction: *F*_2,24_ = 0.50, *P* = 0.61; main effect of Age: *F*_2,24_ = 0.72, *P* = 0.50; main effect of BDNF: *F*_1,24_ = 1.73, *P* = 0.20.). **c**, Olig2⁺PDGFRα⁺ cell numbers were unchanged (ANOVA, Age × BDNF interaction: *F*_2,24_ = 0.50, *P* = 0.62; main effect of Age: *F*_2,24_ = 0.28, *P* = 0.76; main effect of BDNF: *F*_1,24_ = 3.74, *P* = 0.07). **d**, CC1^+^Olig2^+^ cell numbers were reduced 5 days after injection (P20), indicating an earlier peak of oligodendrogenesis (ANOVA, Age × BDNF interaction: *F*_2,26_ = 5.52, *P* < 0.05). **e**, Representative CC1 staining confirms an age-advanced shift in oligodendrogenesis after BDNF treatment. **f**, After PrL BDNF treatment, mice were conditioned at P20 and tested 15 days post-training. **g**, The BDNF-treated group exhibited increased freezing compared to the control group (unpaired t test, *t* = 5.04, *P* < 0.01).

**Extended Data Table 1.**
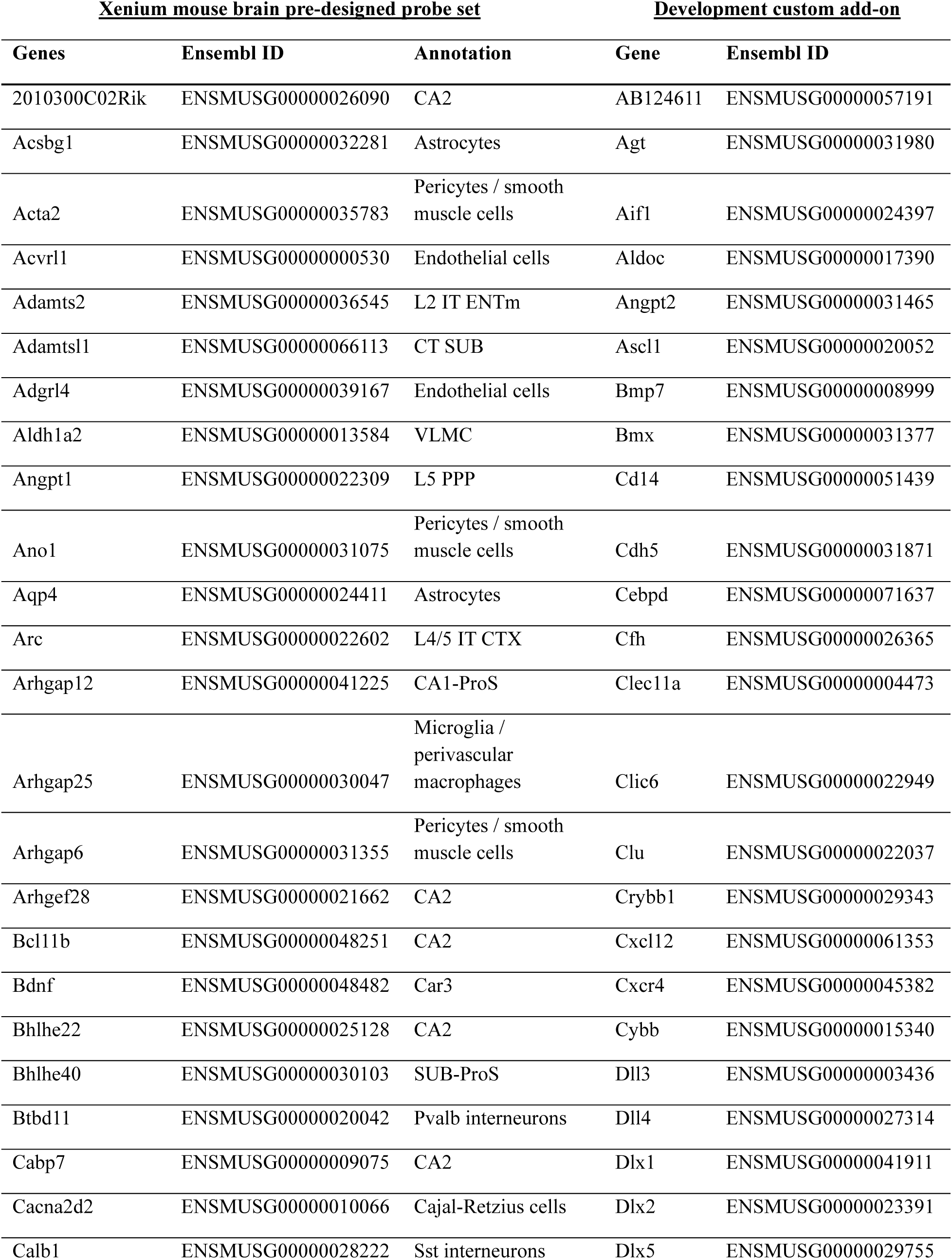
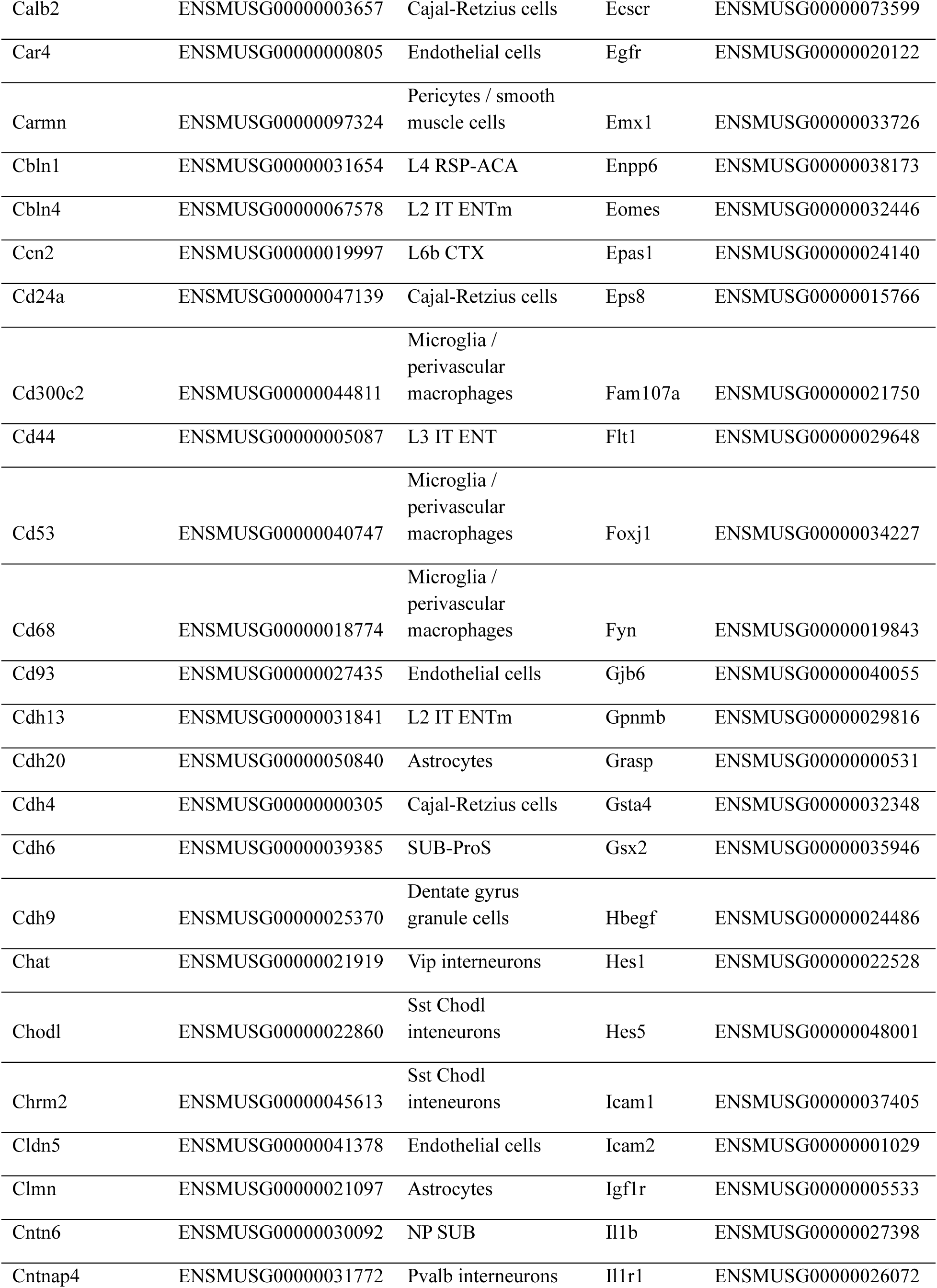
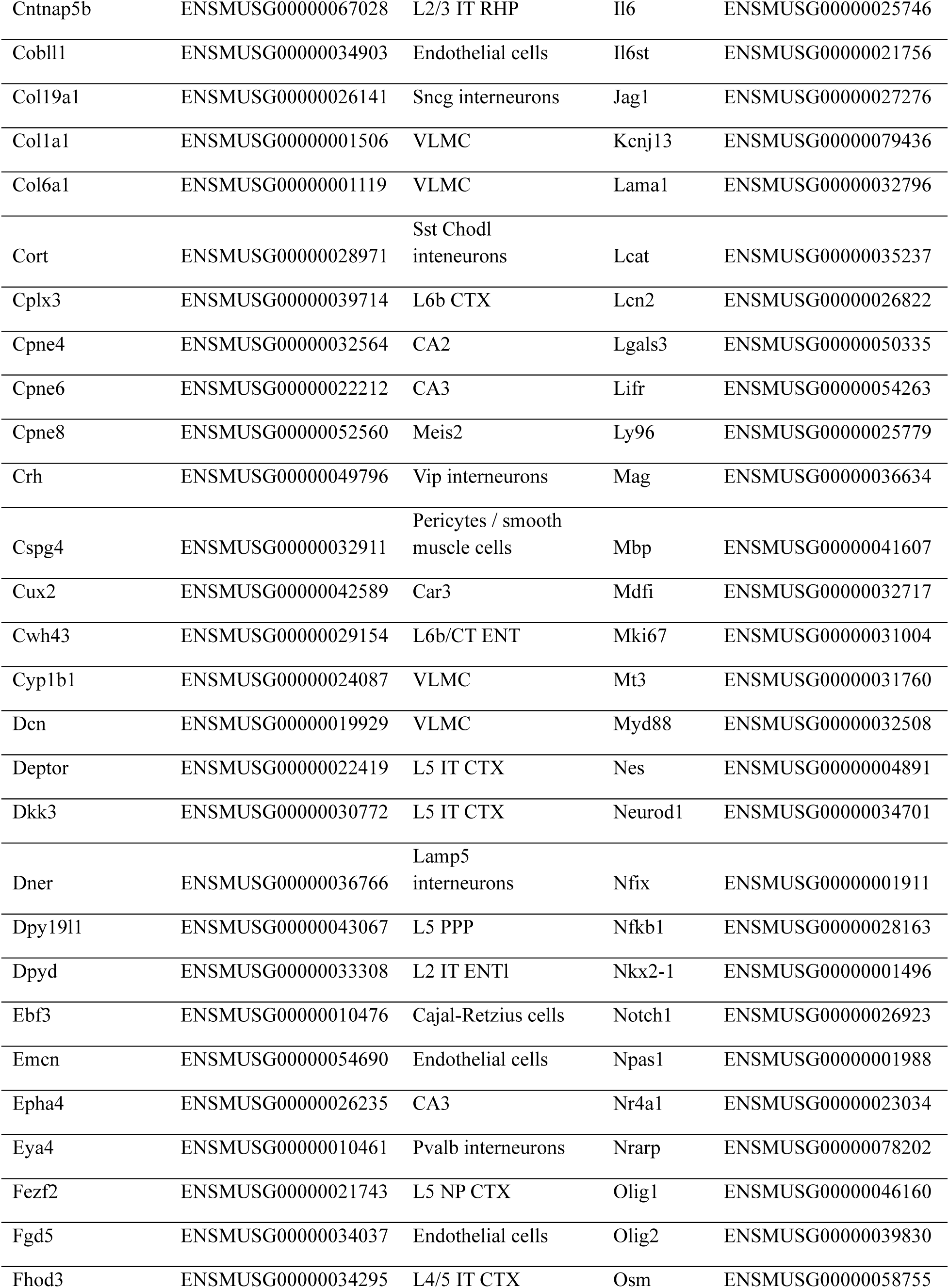
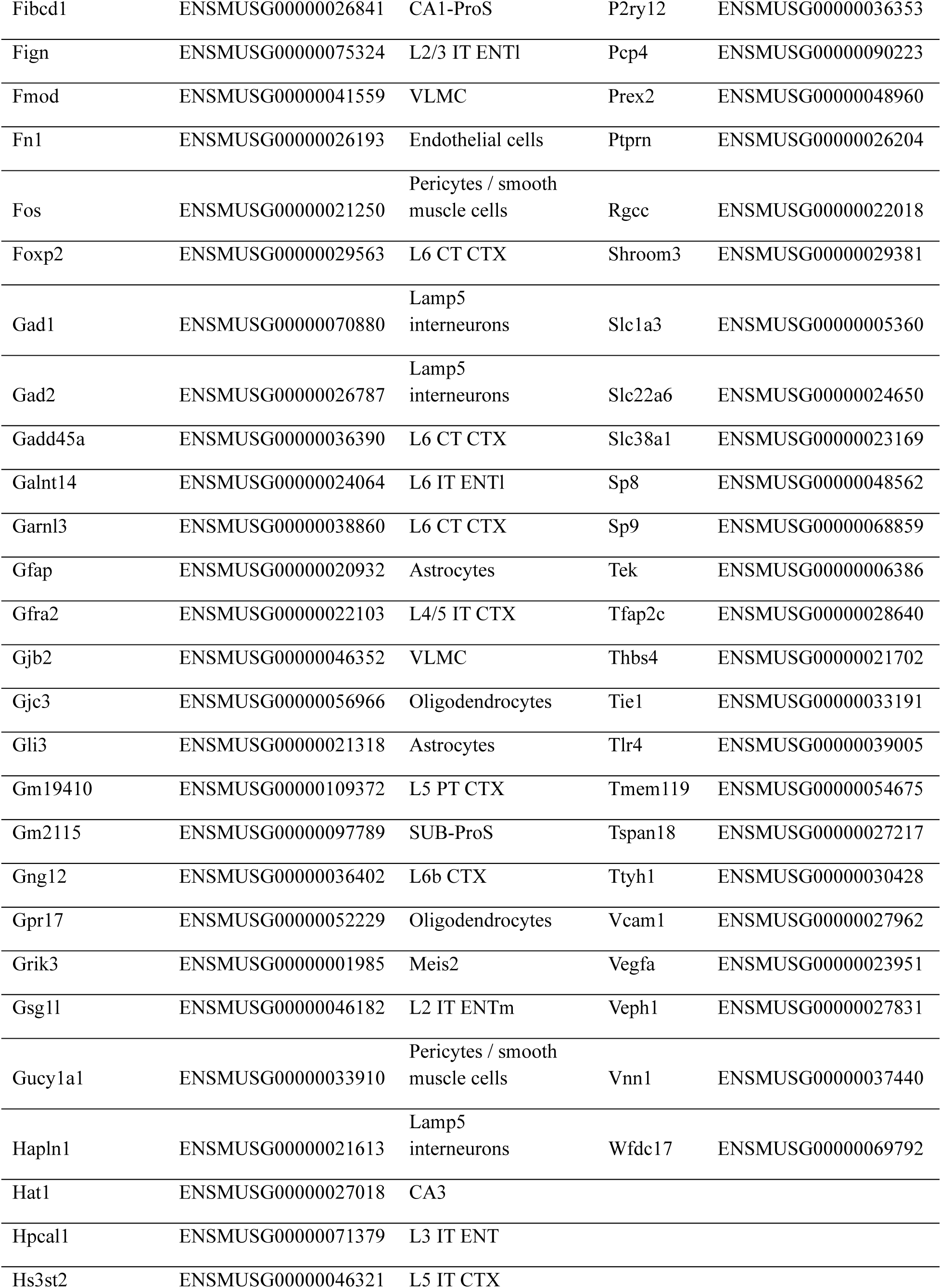
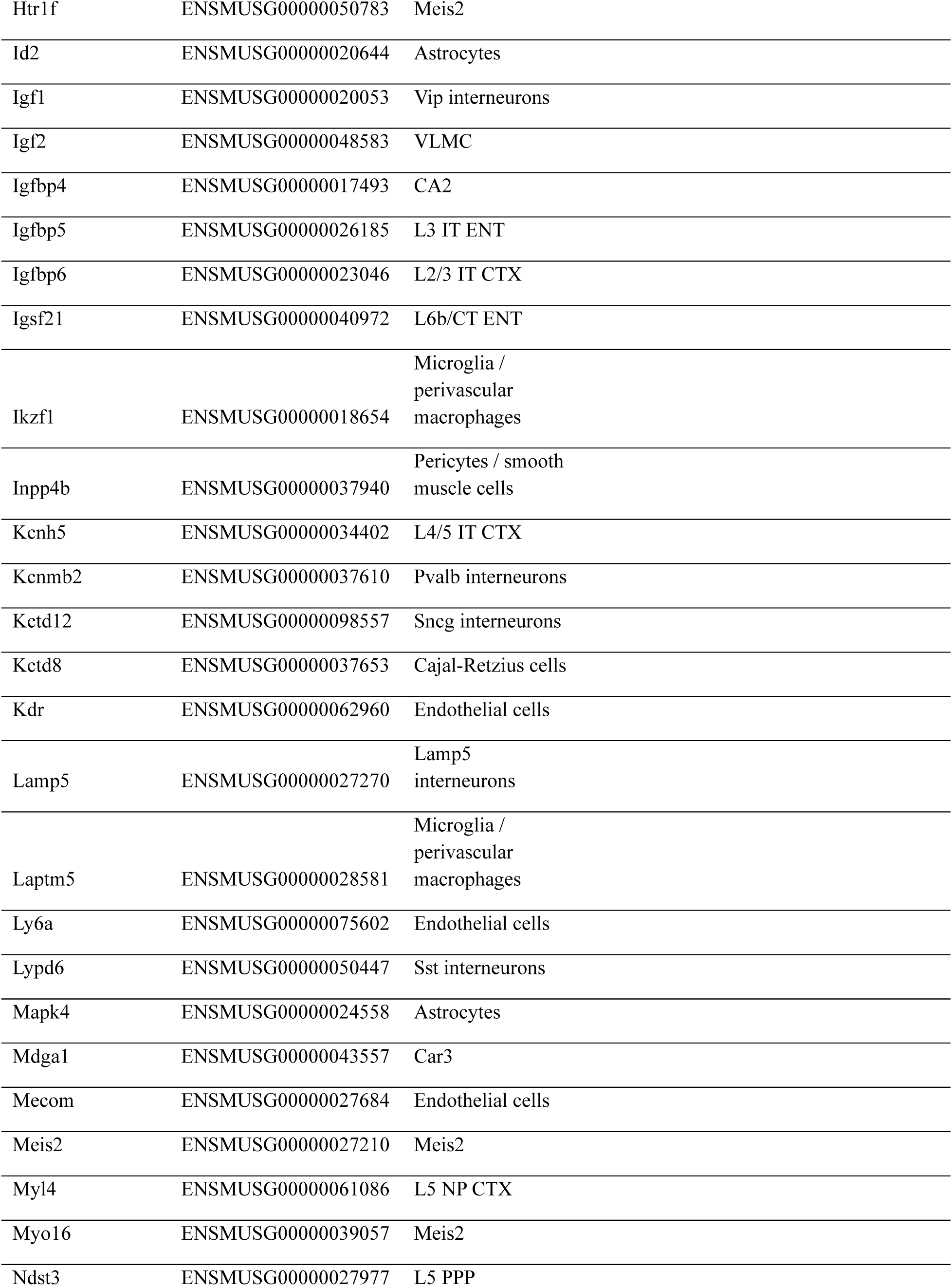
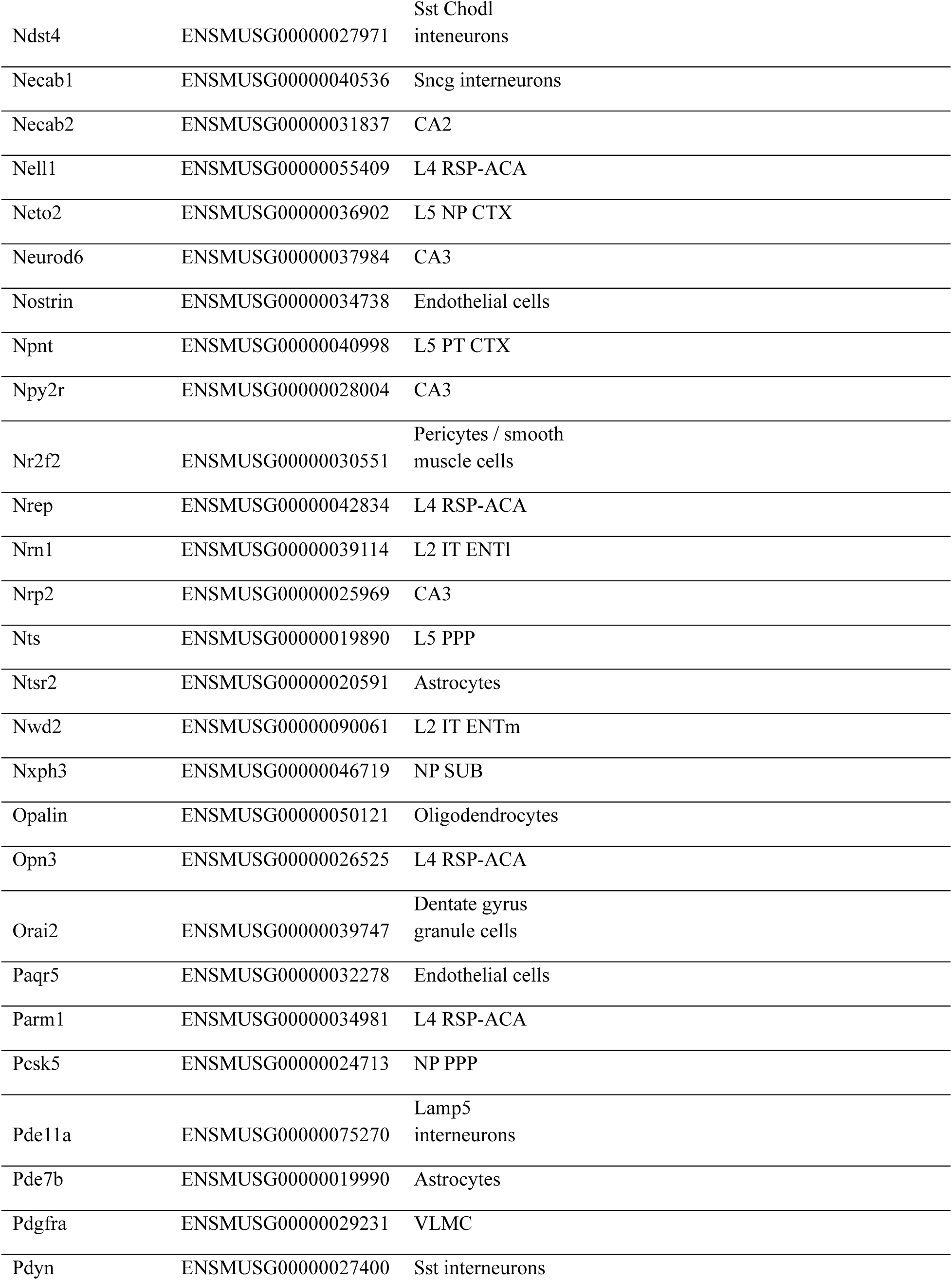
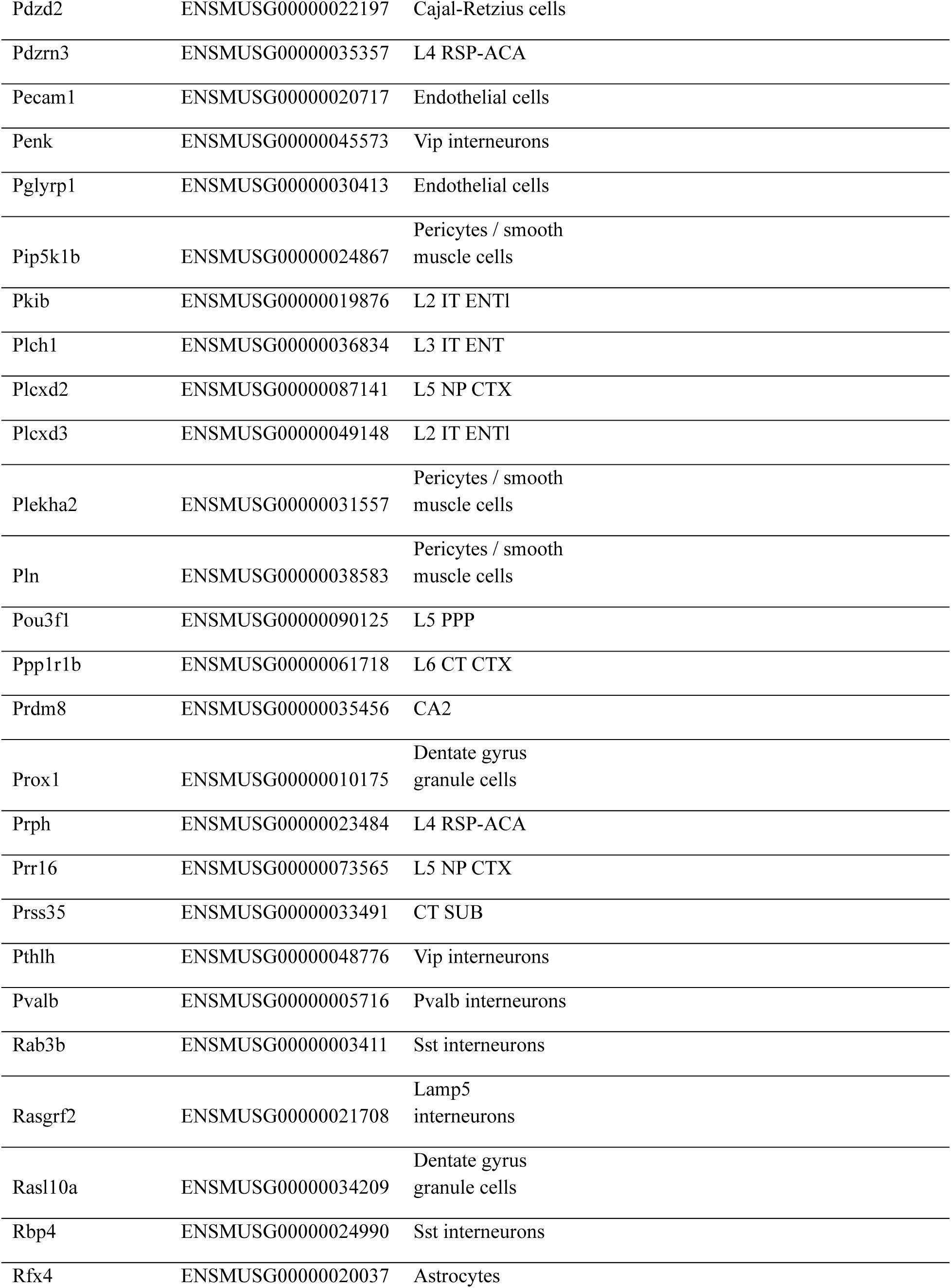
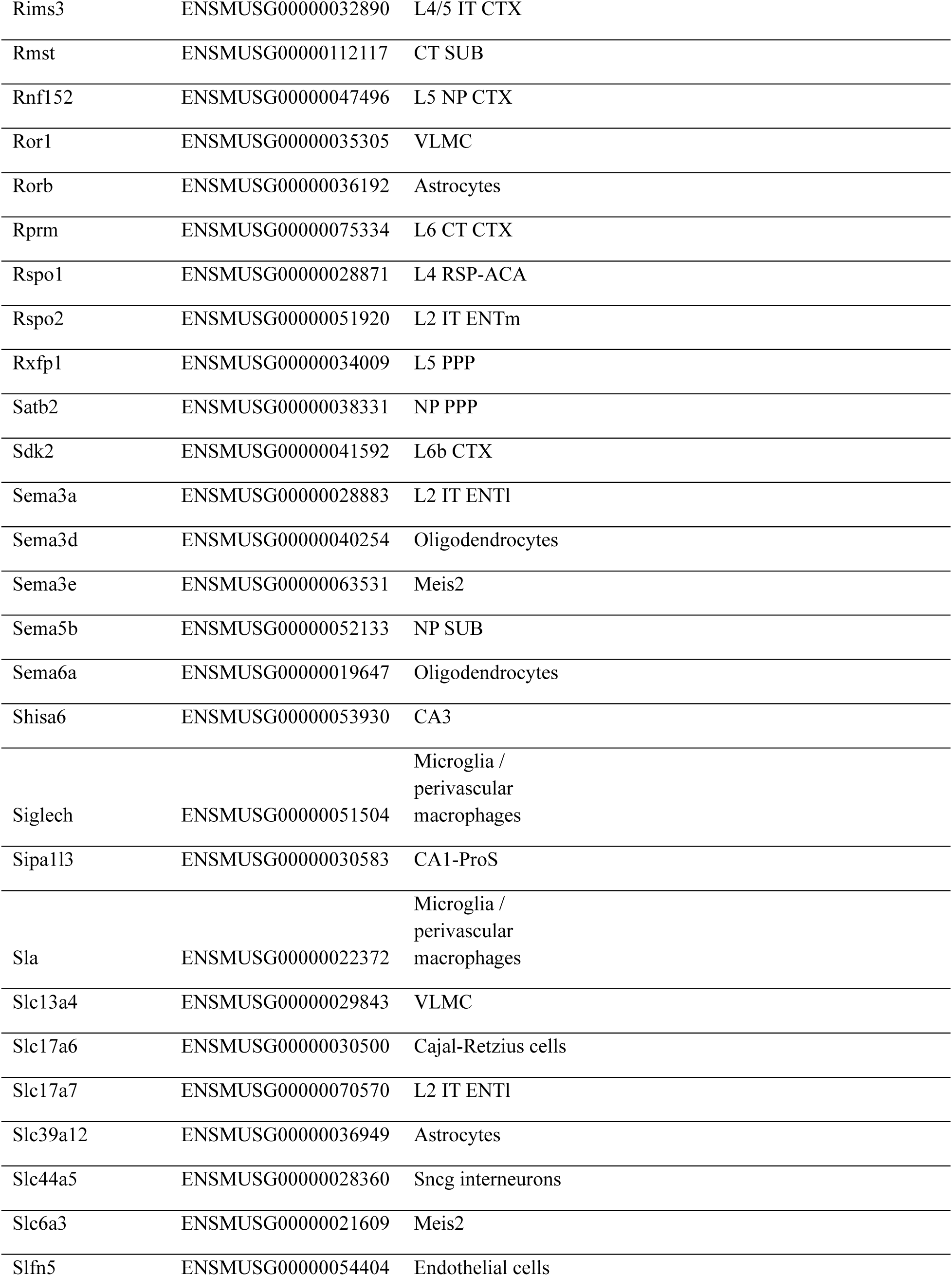
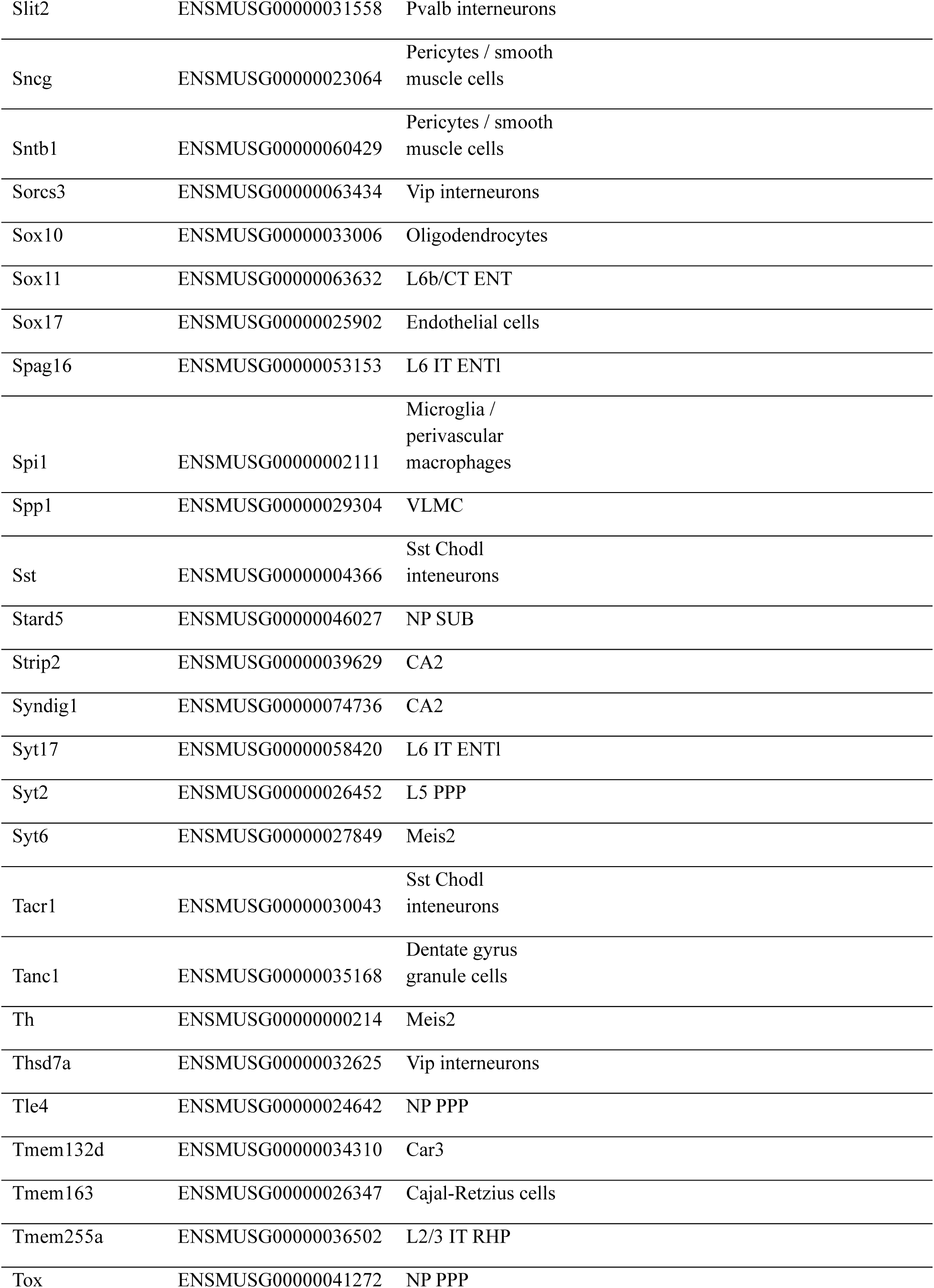
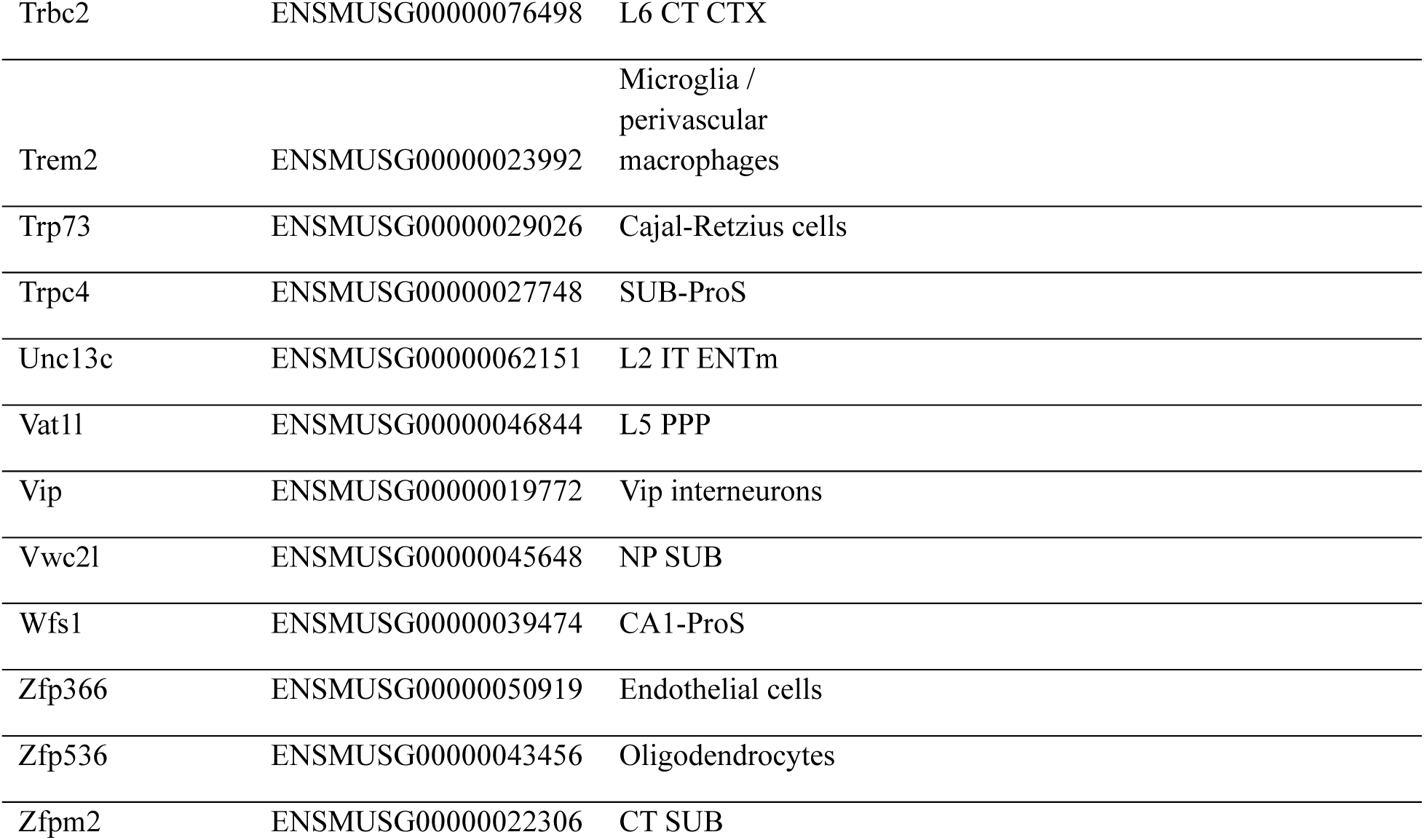
Xenium probe sets.

**Extended Data Table 2.**
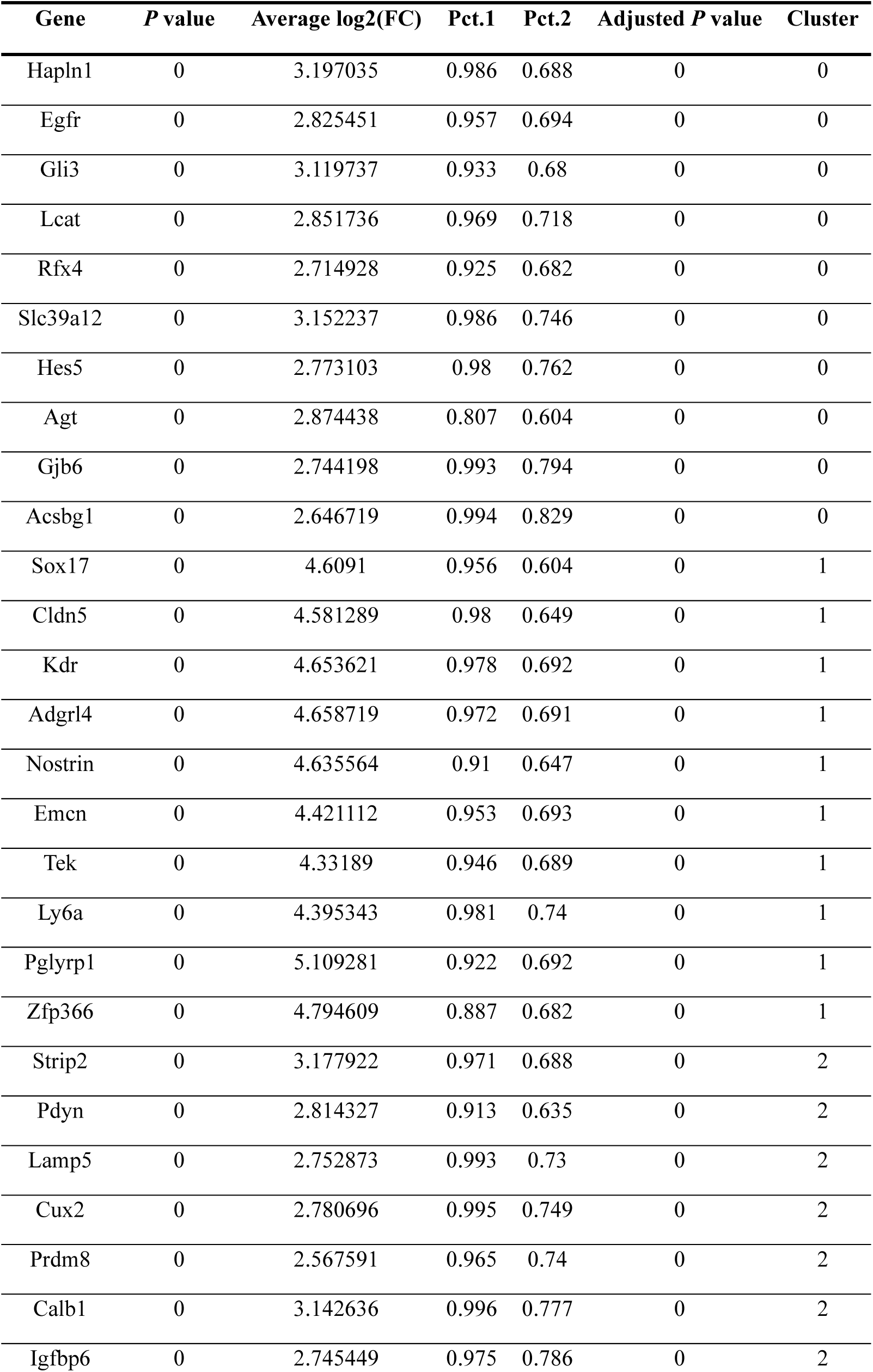
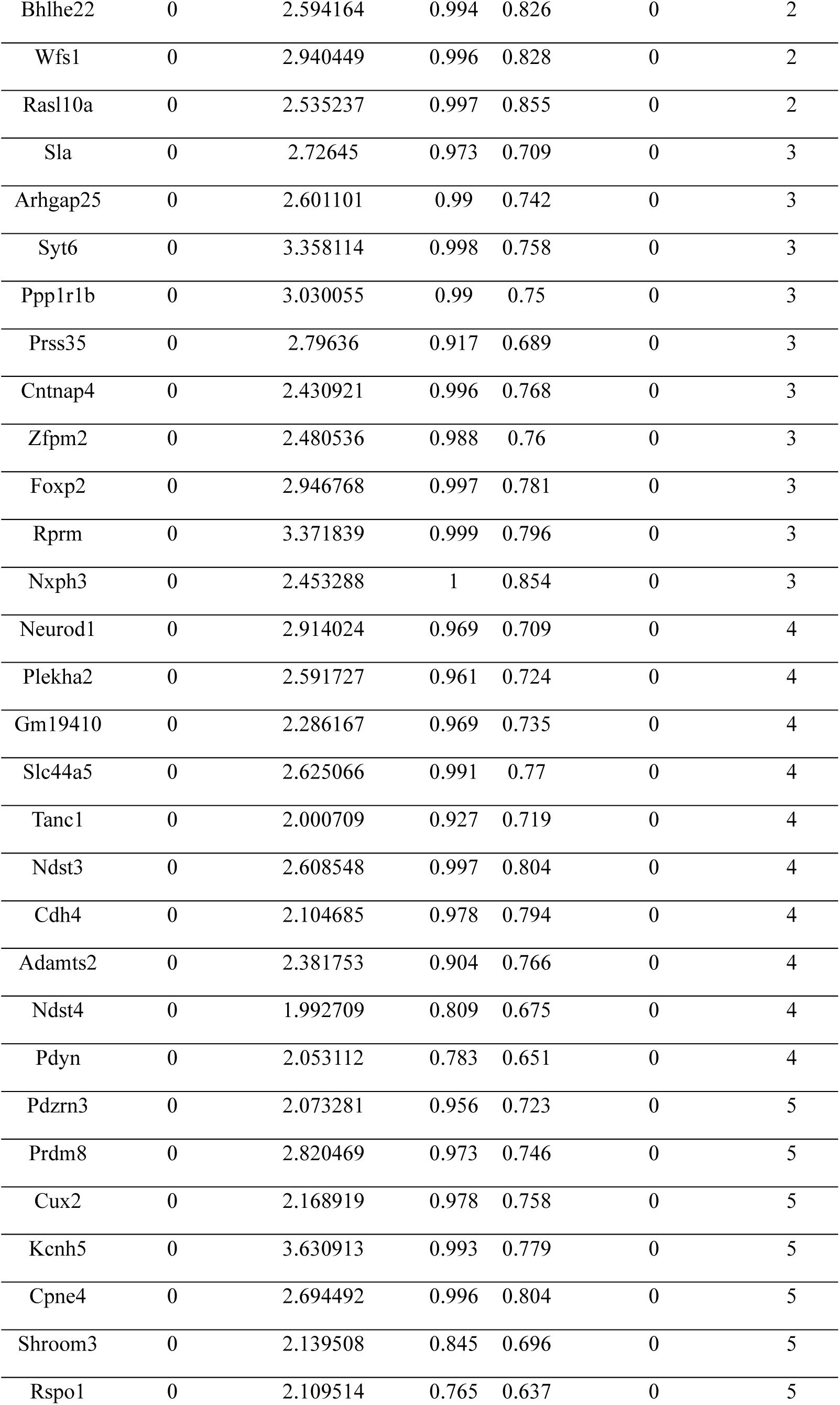
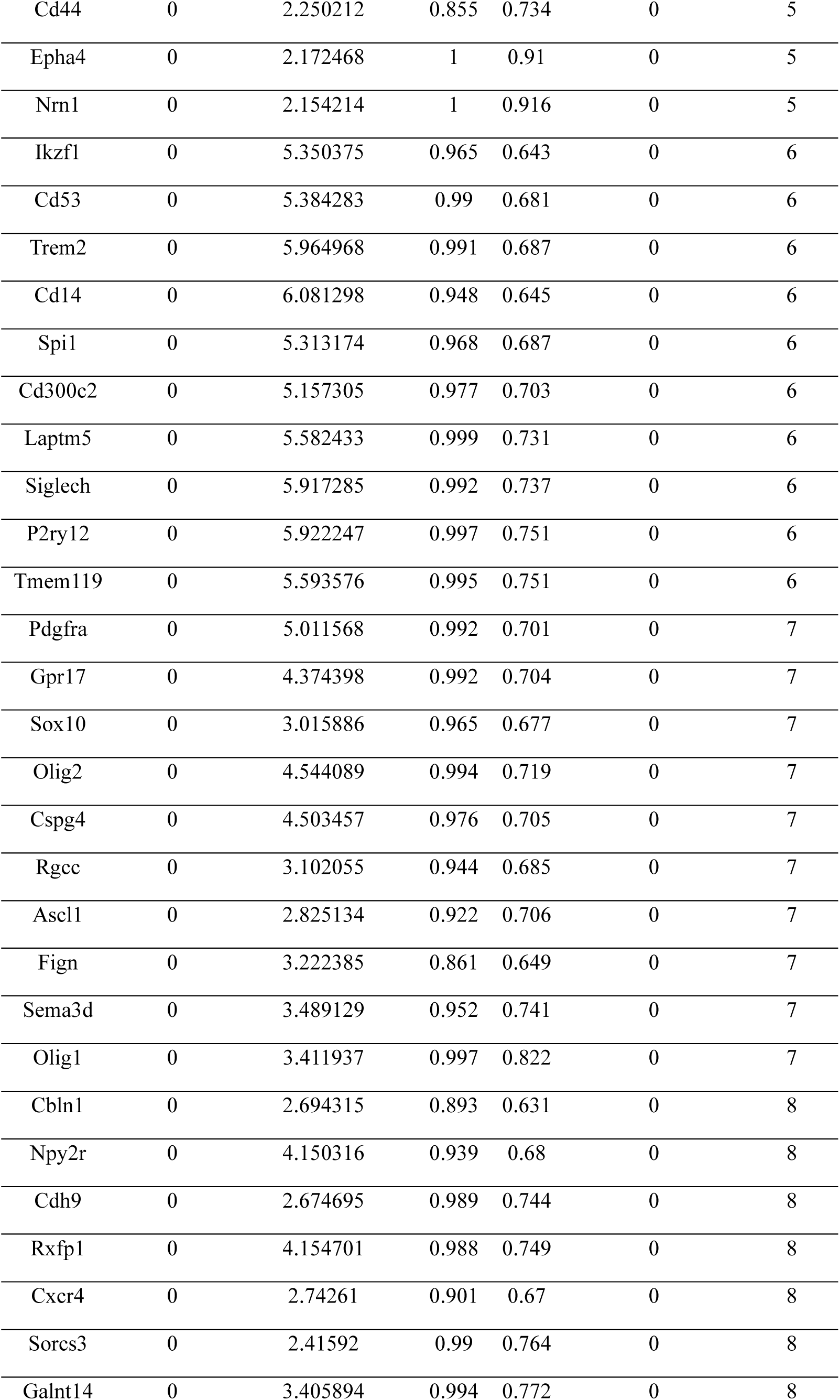
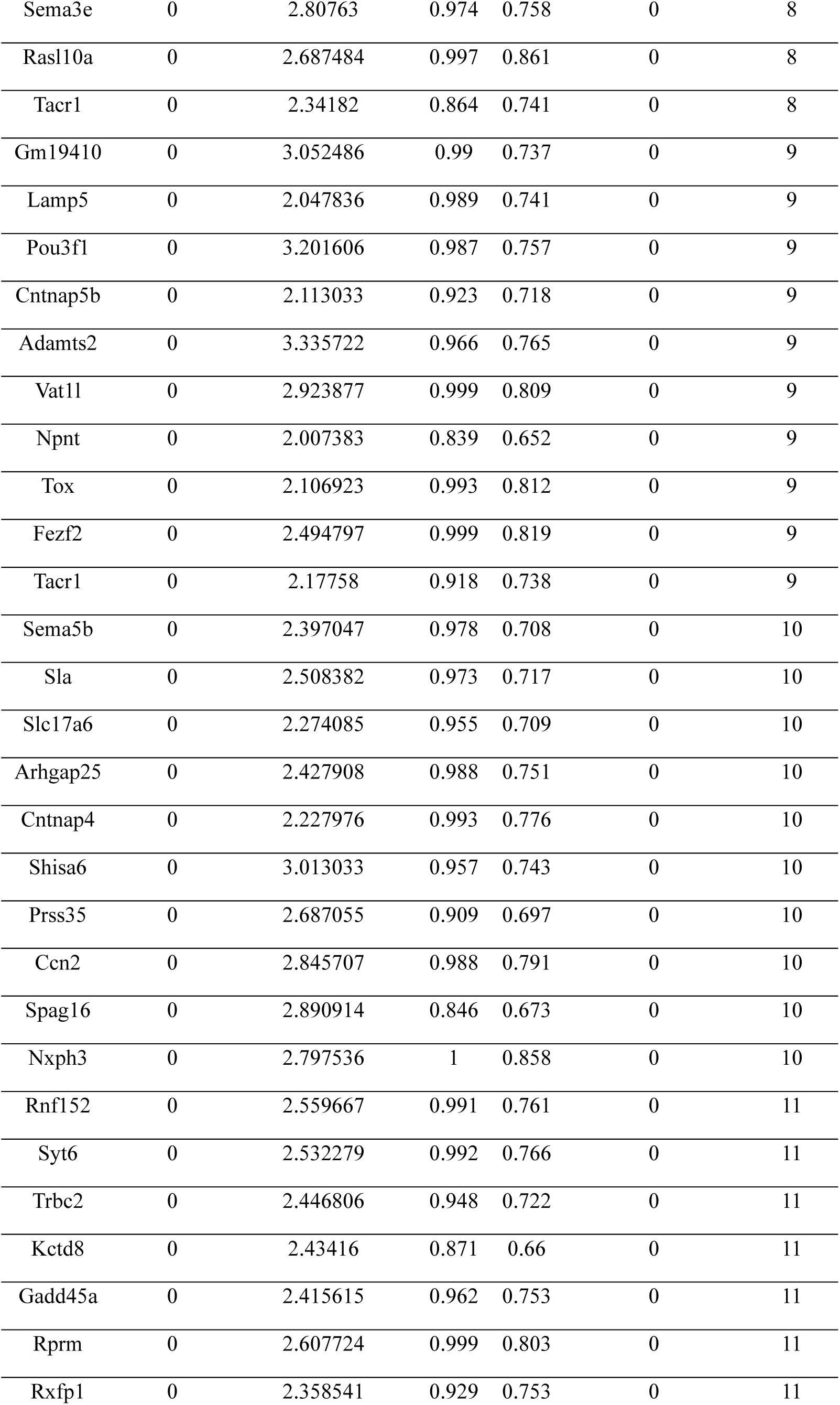
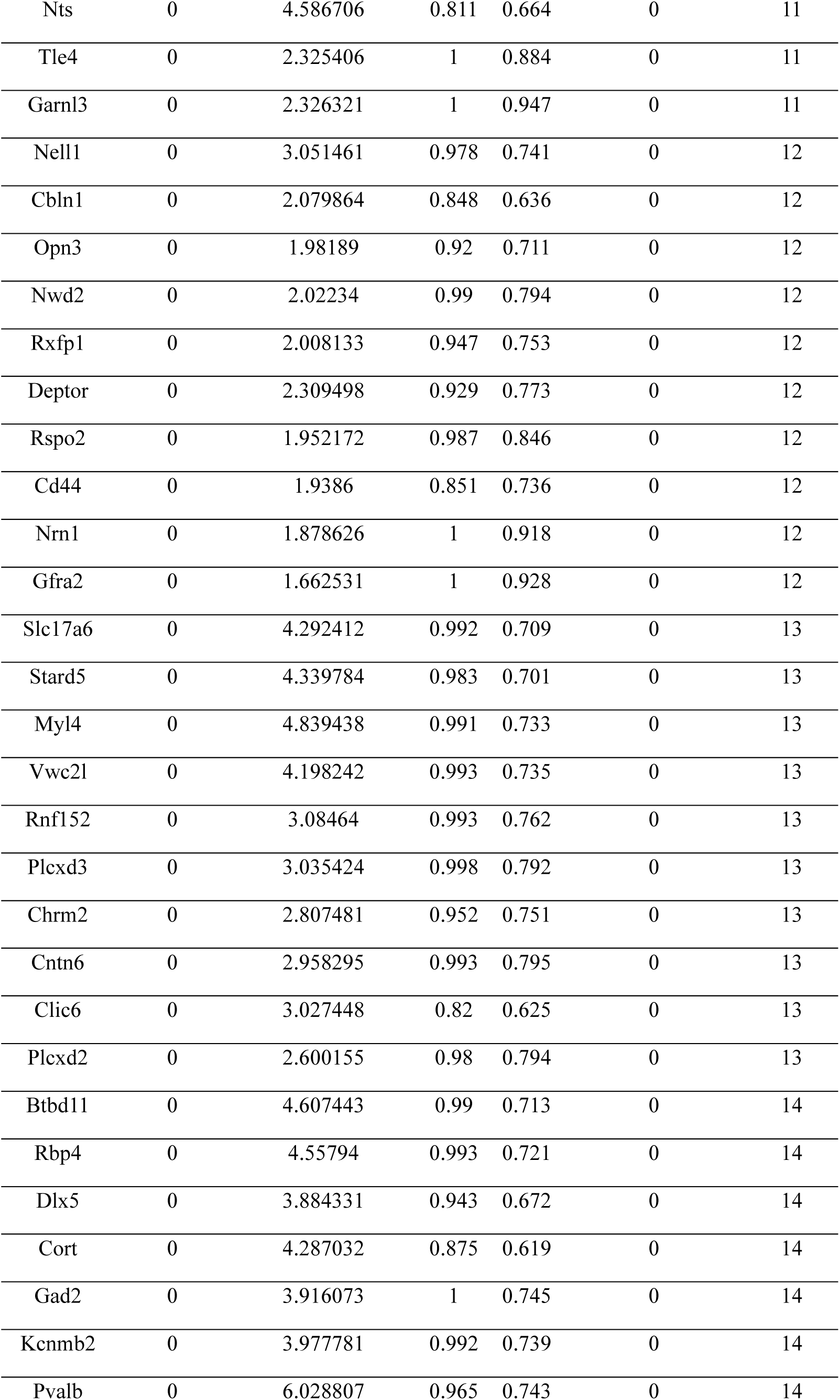
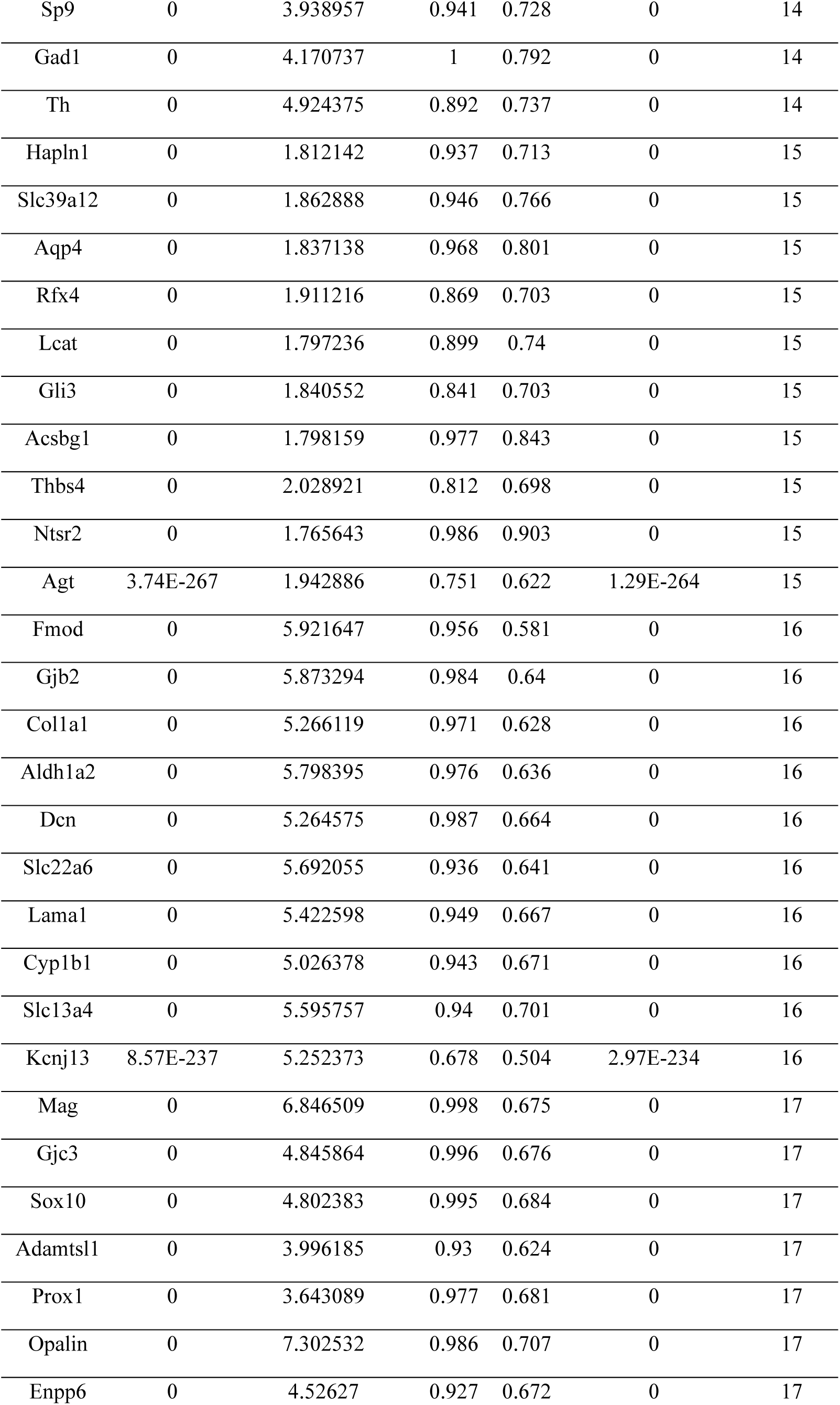
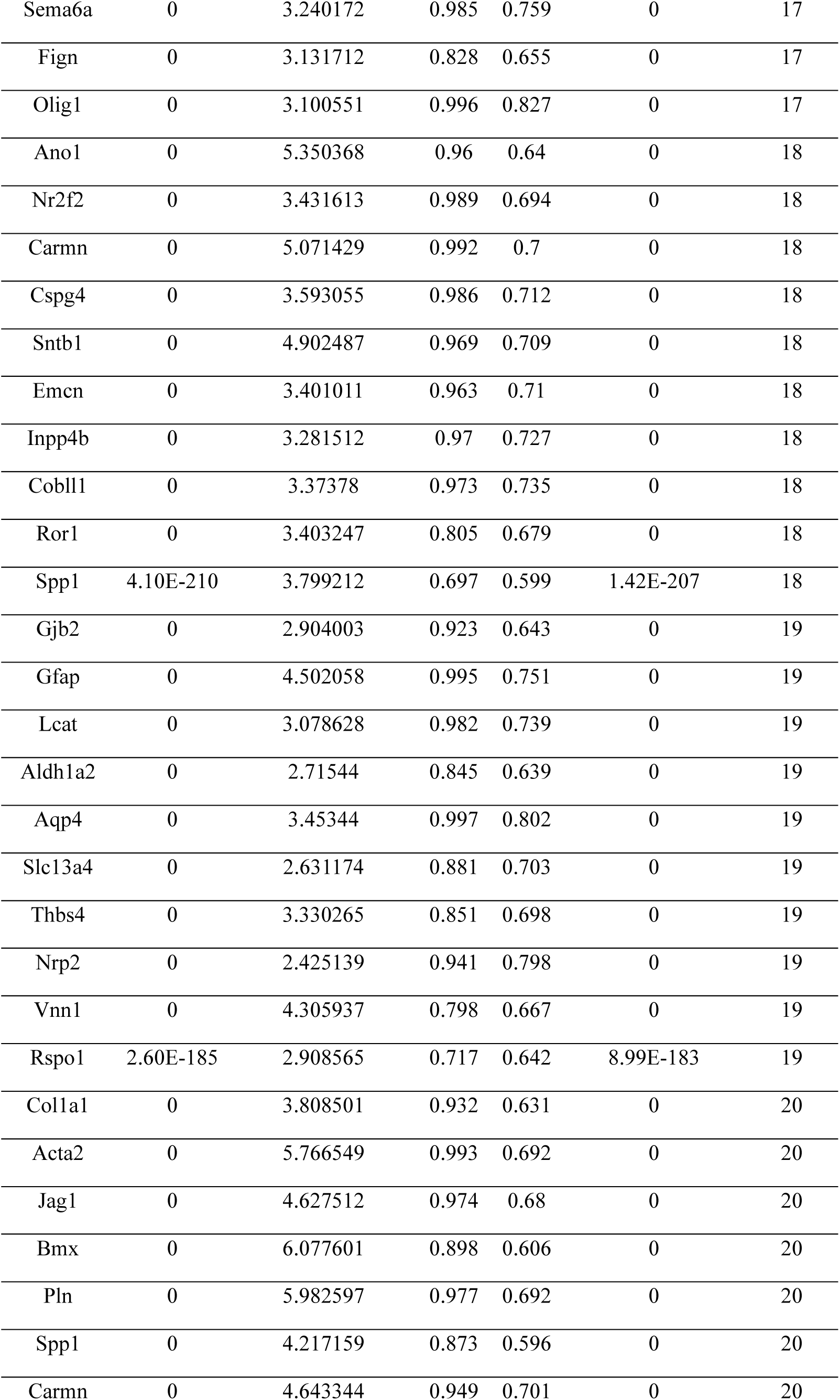
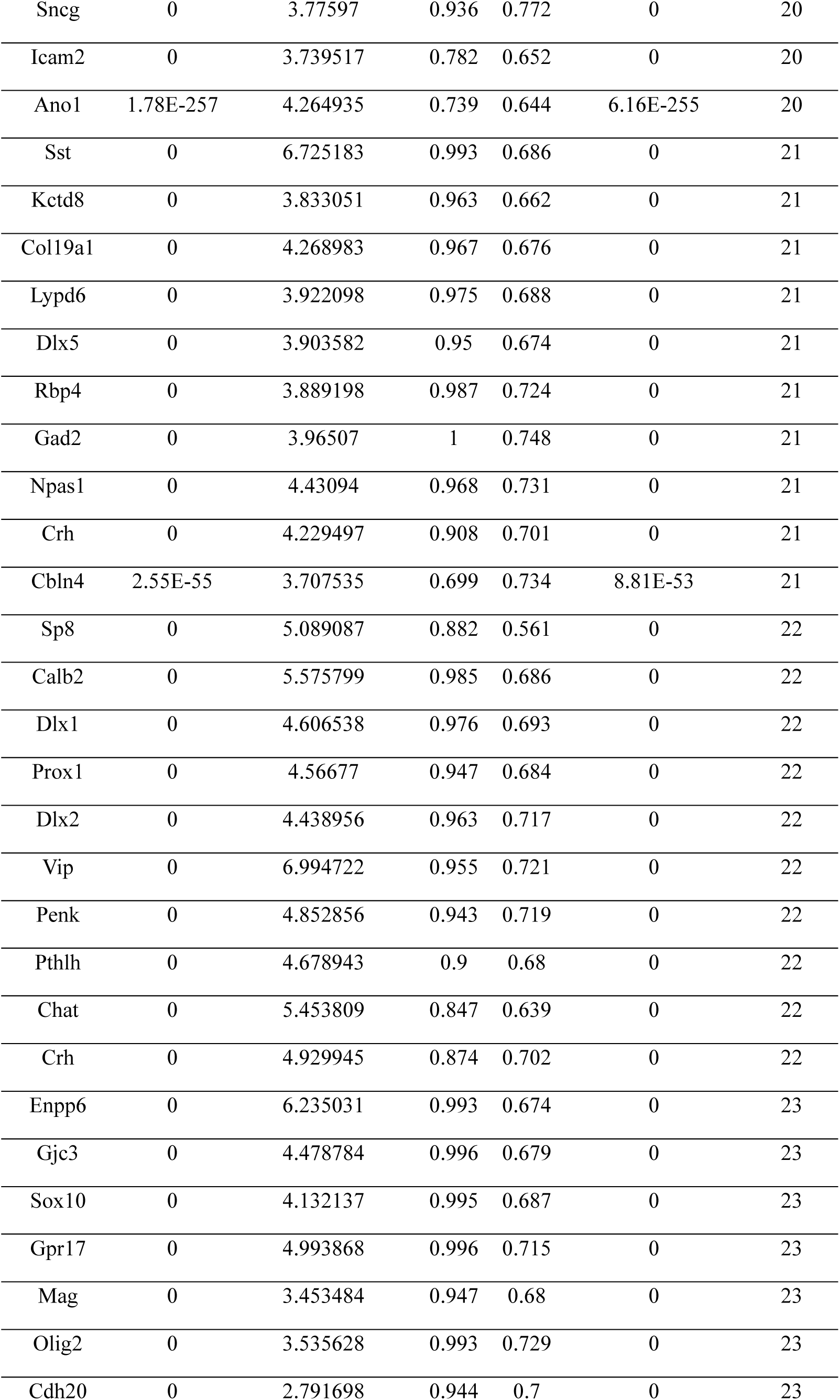
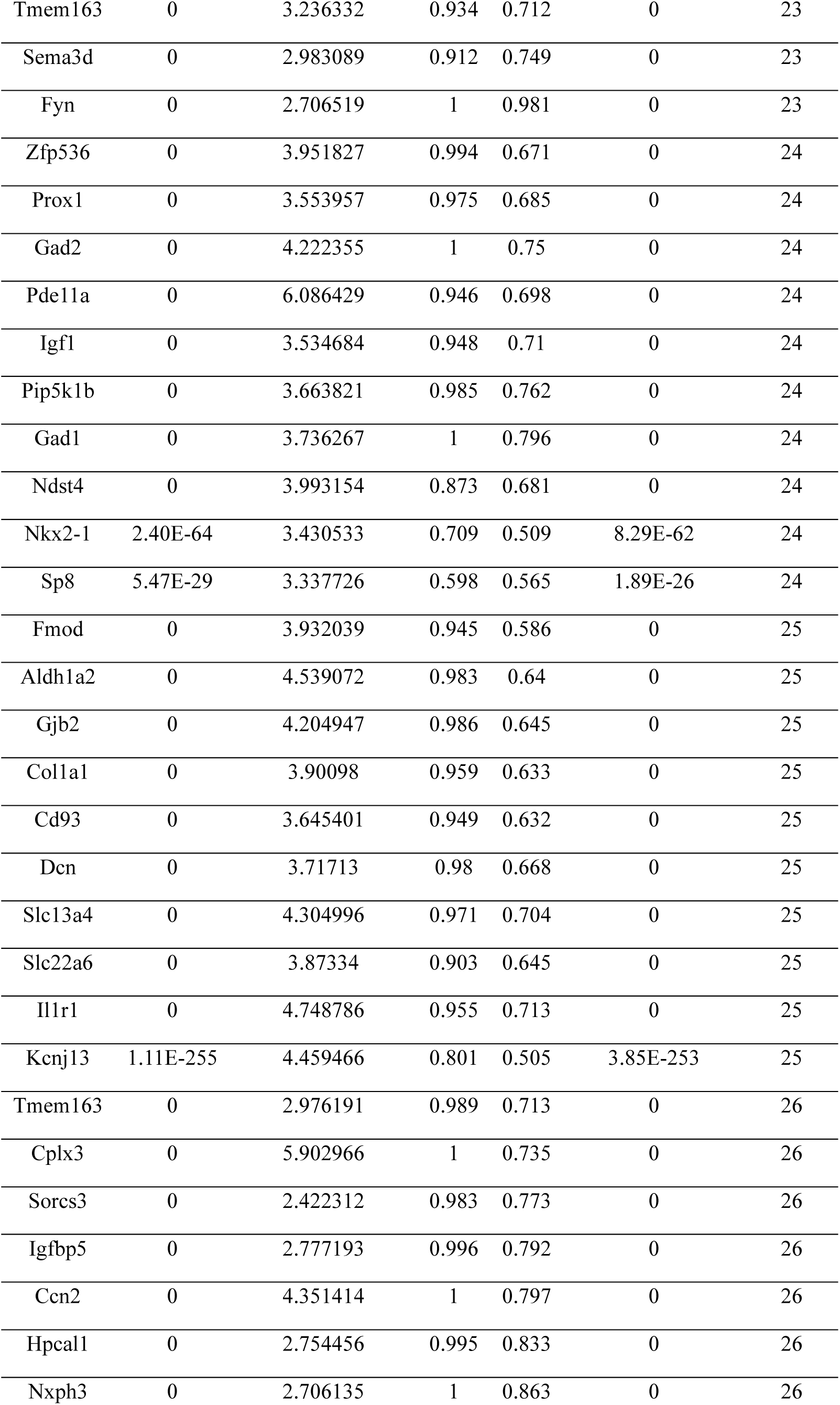
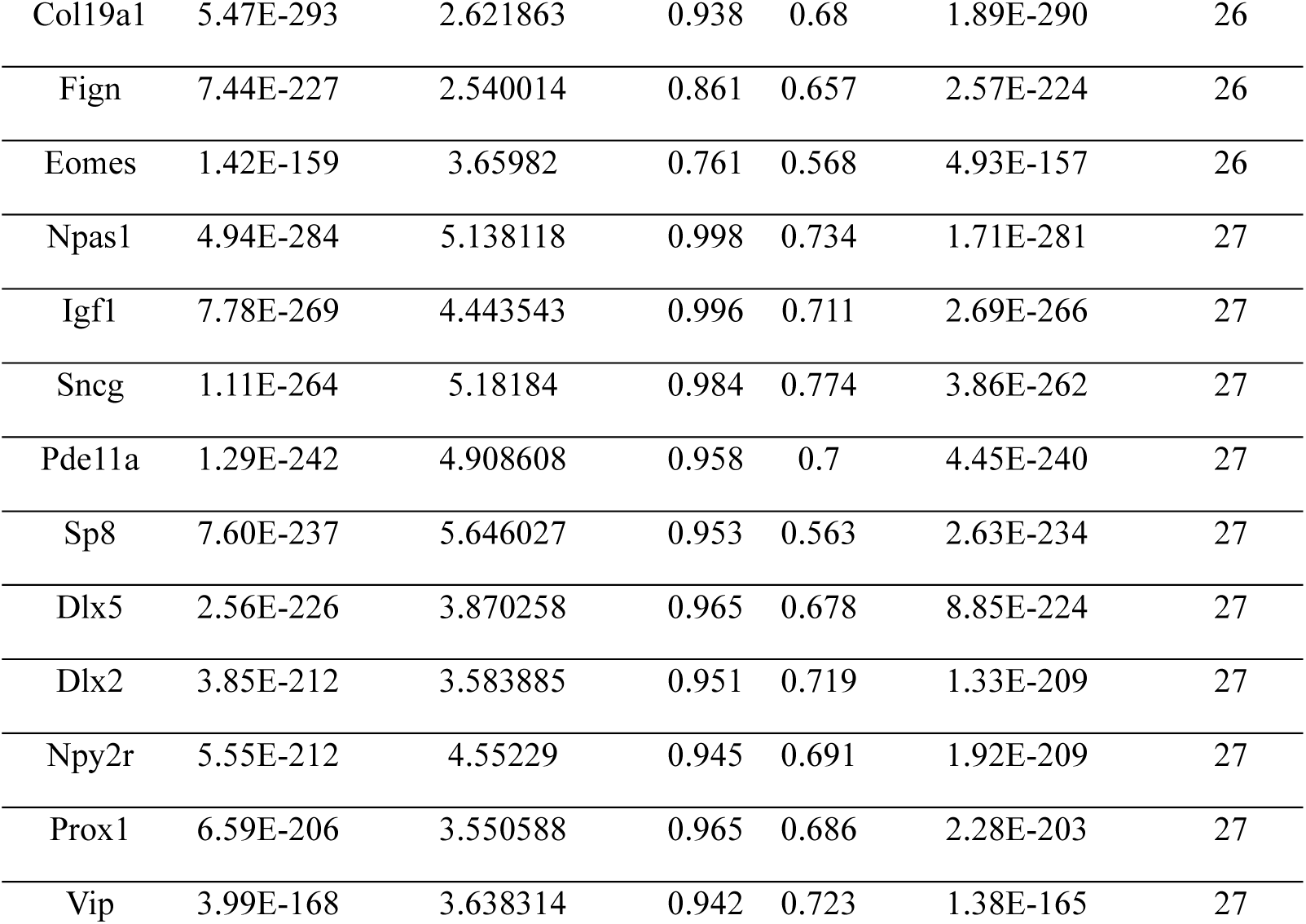
Top 10 marker genes for each cluster identified in all of the PFC population. (Pct.1: percentage of cells where the gene is detected in the target cluster, Pct.2: percentage of cells where the gene is detected in all other clusters, FC: Fold change)

**Extended Data Table 3.** Cell counts for each cell-type per analyzed sample.

| Cluster | Sample ID | Count | Age group | Percentage | Cell Type |
| --- | --- | --- | --- | --- | --- |
| 0 | P15_1M | 978 | P15 | 10.10018 | Cortical astrocytes |
| 15 | P15_1M | 227 | P15 | 2.344315 | Cortical astrocytes |
| 19 | P15_1M | 153 | P15 | 1.580089 | Cortical astrocytes |
| 0 | P15_2F | 848 | P15 | 9.744886 | Cortical astrocytes |
| 15 | P15_2F | 185 | P15 | 2.125948 | Cortical astrocytes |
| 19 | P15_2F | 168 | P15 | 1.930591 | Cortical astrocytes |
| 0 | P15_3F | 1130 | P15 | 11.17263 | Cortical astrocytes |
| 15 | P15_3F | 248 | P15 | 2.452047 | Cortical astrocytes |
| 19 | P15_3F | 177 | P15 | 1.750049 | Cortical astrocytes |
| 0 | P20_1M | 901 | P20 | 10.17045 | Cortical astrocytes |
| 15 | P20_1M | 203 | P20 | 2.291455 | Cortical astrocytes |
| 19 | P20_1M | 162 | P20 | 1.828649 | Cortical astrocytes |
| 0 | P20_2M | 865 | P20 | 9.645406 | Cortical astrocytes |
| 15 | P20_2M | 224 | P20 | 2.49777 | Cortical astrocytes |
| 19 | P20_2M | 174 | P20 | 1.940232 | Cortical astrocytes |
| 0 | P20_3M | 1076 | P20 | 10.84131 | Cortical astrocytes |
| 15 | P20_3M | 227 | P20 | 2.287154 | Cortical astrocytes |
| 19 | P20_3M | 222 | P20 | 2.236776 | Cortical astrocytes |
| 0 | P25_1M | 774 | P25 | 9.242895 | Cortical astrocytes |
| 15 | P25_1M | 262 | P25 | 3.128732 | Cortical astrocytes |
| 19 | P25_1M | 146 | P25 | 1.743492 | Cortical astrocytes |
| 0 | P25_2M | 897 | P25 | 9.800066 | Cortical astrocytes |
| 15 | P25_2M | 207 | P25 | 2.261554 | Cortical astrocytes |
| 19 | P25_2M | 162 | P25 | 1.769912 | Cortical astrocytes |
| 0 | P25_3F | 798 | P25 | 10.43683 | Cortical astrocytes |
| 15 | P25_3F | 176 | P25 | 2.301857 | Cortical astrocytes |
| 19 | P25_3F | 149 | P25 | 1.948731 | Cortical astrocytes |
| 0 | P60_1M | 744 | P60 | 10.20576 | Cortical astrocytes |
| 15 | P60_1M | 154 | P60 | 2.112483 | Cortical astrocytes |
| 19 | P60_1M | 134 | P60 | 1.838134 | Cortical astrocytes |
| 0 | P60_2F | 756 | P60 | 10.24251 | Cortical astrocytes |
| 15 | P60_2F | 174 | P60 | 2.357404 | Cortical astrocytes |
| 19 | P60_2F | 120 | P60 | 1.625796 | Cortical astrocytes |
| 0 | P60_3M | 820 | P60 | 10.53578 | Cortical astrocytes |
| 15 | P60_3M | 154 | P60 | 1.978671 | Cortical astrocytes |
| 19 | P60_3M | 127 | P60 | 1.631762 | Cortical astrocytes |
| 1 | P15_1M | 819 | P15 | 8.458122 | Endothelial |
| 1 | P15_2F | 716 | P15 | 8.227994 | Endothelial |
| 1 | P15_3F | 837 | P15 | 8.275658 | Endothelial |
| 1 | P20_1M | 862 | P20 | 9.730218 | Endothelial |
| 1 | P20_2M | 768 | P20 | 8.563782 | Endothelial |
| 1 | P20_3M | 810 | P20 | 8.161209 | Endothelial |
| 1 | P25_1M | 728 | P25 | 8.693575 | Endothelial |
| 1 | P25_2M | 852 | P25 | 9.308423 | Endothelial |
| 1 | P25_3F | 706 | P25 | 9.233586 | Endothelial |
| 1 | P60_1M | 580 | P60 | 7.956104 | Endothelial |
| 1 | P60_2F | 667 | P60 | 9.036716 | Endothelial |
| 1 | P60_3M | 732 | P60 | 9.405114 | Endothelial |
| 23 | P15_1M | 138 | P15 | 1.425178 | Immature OLs |
| 23 | P15_2F | 155 | P15 | 1.7812 | Immature OLs |
| 23 | P15_3F | 119 | P15 | 1.176587 | Immature OLs |
| 23 | P20_1M | 129 | P20 | 1.456146 | Immature OLs |
| 23 | P20_2M | 116 | P20 | 1.293488 | Immature OLs |
| 23 | P20_3M | 138 | P20 | 1.390428 | Immature OLs |
| 23 | P25_1M | 60 | P25 | 0.716503 | Immature OLs |
| 23 | P25_2M | 71 | P25 | 0.775702 | Immature OLs |
| 23 | P25_3F | 49 | P25 | 0.640858 | Immature OLs |
| 23 | P60_1M | 36 | P60 | 0.493827 | Immature OLs |
| 23 | P60_2F | 21 | P60 | 0.284514 | Immature OLs |
| 23 | P60_3M | 40 | P60 | 0.513941 | Immature OLs |
| 2 | P15_1M | 829 | P15 | 8.561396 | L2/3 Cortical excitatory |
| 5 | P15_1M | 556 | P15 | 5.742022 | L2/3 Cortical excitatory |
| 2 | P15_2F | 722 | P15 | 8.296943 | L2/3 Cortical excitatory |
| 5 | P15_2F | 533 | P15 | 6.125029 | L2/3 Cortical excitatory |
| 2 | P15_3F | 896 | P15 | 8.859007 | L2/3 Cortical excitatory |
| 5 | P15_3F | 575 | P15 | 5.685189 | L2/3 Cortical excitatory |
| 2 | P20_1M | 720 | P20 | 8.127328 | L2/3 Cortical excitatory |
| 5 | P20_1M | 479 | P20 | 5.406931 | L2/3 Cortical excitatory |
| 2 | P20_2M | 748 | P20 | 8.340767 | L2/3 Cortical excitatory |
| 5 | P20_2M | 496 | P20 | 5.530776 | L2/3 Cortical excitatory |
| 2 | P20_3M | 775 | P20 | 7.808564 | L2/3 Cortical excitatory |
| 5 | P20_3M | 514 | P20 | 5.178841 | L2/3 Cortical excitatory |
| 2 | P25_1M | 703 | P25 | 8.395032 | L2/3 Cortical excitatory |
| 5 | P25_1M | 456 | P25 | 5.445426 | L2/3 Cortical excitatory |
| 2 | P25_2M | 779 | P25 | 8.510871 | L2/3 Cortical excitatory |
| 5 | P25_2M | 499 | P25 | 5.451764 | L2/3 Cortical excitatory |
| 2 | P25_3F | 665 | P25 | 8.697358 | L2/3 Cortical excitatory |
| 5 | P25_3F | 382 | P25 | 4.996076 | L2/3 Cortical excitatory |
| 2 | P60_1M | 661 | P60 | 9.067215 | L2/3 Cortical excitatory |
| 5 | P60_1M | 368 | P60 | 5.048011 | L2/3 Cortical excitatory |
| 2 | P60_2F | 626 | P60 | 8.481236 | L2/3 Cortical excitatory |
| 5 | P60_2F | 387 | P60 | 5.243192 | L2/3 Cortical excitatory |
| 2 | P60_3M | 650 | P60 | 8.351535 | L2/3 Cortical excitatory |
| 5 | P60_3M | 403 | P60 | 5.177952 | L2/3 Cortical excitatory |
| 4 | P15_1M | 433 | P15 | 4.471755 | L5 Cortical excitatory |
| 8 | P15_1M | 442 | P15 | 4.564701 | L5 Cortical excitatory |
| 9 | P15_1M | 460 | P15 | 4.750594 | L5 Cortical excitatory |
| 12 | P15_1M | 330 | P15 | 3.408035 | L5 Cortical excitatory |
| 13 | P15_1M | 286 | P15 | 2.95363 | L5 Cortical excitatory |
| 4 | P15_2F | 420 | P15 | 4.826477 | L5 Cortical excitatory |
| 8 | P15_2F | 374 | P15 | 4.297863 | L5 Cortical excitatory |
| 9 | P15_2F | 365 | P15 | 4.194438 | L5 Cortical excitatory |
| 12 | P15_2F | 371 | P15 | 4.263388 | L5 Cortical excitatory |
| 13 | P15_2F | 270 | P15 | 3.102735 | L5 Cortical excitatory |
| 4 | P15_3F | 584 | P15 | 5.774174 | L5 Cortical excitatory |
| 8 | P15_3F | 478 | P15 | 4.726122 | L5 Cortical excitatory |
| 9 | P15_3F | 344 | P15 | 3.401226 | L5 Cortical excitatory |
| 12 | P15_3F | 313 | P15 | 3.09472 | L5 Cortical excitatory |
| 13 | P15_3F | 318 | P15 | 3.144157 | L5 Cortical excitatory |
| 4 | P20_1M | 343 | P20 | 3.871769 | L5 Cortical excitatory |
| 8 | P20_1M | 446 | P20 | 5.034428 | L5 Cortical excitatory |
| 9 | P20_1M | 443 | P20 | 5.000564 | L5 Cortical excitatory |
| 12 | P20_1M | 361 | P20 | 4.074952 | L5 Cortical excitatory |
| 13 | P20_1M | 244 | P20 | 2.754261 | L5 Cortical excitatory |
| 4 | P20_2M | 484 | P20 | 5.396967 | L5 Cortical excitatory |
| 8 | P20_2M | 382 | P20 | 4.25959 | L5 Cortical excitatory |
| 9 | P20_2M | 421 | P20 | 4.694469 | L5 Cortical excitatory |
| 12 | P20_2M | 322 | P20 | 3.590544 | L5 Cortical excitatory |
| 13 | P20_2M | 258 | P20 | 2.876896 | L5 Cortical excitatory |
| 4 | P20_3M | 979 | P20 | 9.86398 | L5 Cortical excitatory |
| 8 | P20_3M | 354 | P20 | 3.566751 | L5 Cortical excitatory |
| 9 | P20_3M | 363 | P20 | 3.657431 | L5 Cortical excitatory |
| 12 | P20_3M | 288 | P20 | 2.901763 | L5 Cortical excitatory |
| 13 | P20_3M | 322 | P20 | 3.244332 | L5 Cortical excitatory |
| 4 | P25_1M | 407 | P25 | 4.860282 | L5 Cortical excitatory |
| 8 | P25_1M | 395 | P25 | 4.716981 | L5 Cortical excitatory |
| 9 | P25_1M | 347 | P25 | 4.143778 | L5 Cortical excitatory |
| 12 | P25_1M | 292 | P25 | 3.486984 | L5 Cortical excitatory |
| 13 | P25_1M | 304 | P25 | 3.630284 | L5 Cortical excitatory |
| 4 | P25_2M | 457 | P25 | 4.992899 | L5 Cortical excitatory |
| 8 | P25_2M | 441 | P25 | 4.818092 | L5 Cortical excitatory |
| 9 | P25_2M | 429 | P25 | 4.686988 | L5 Cortical excitatory |
| 12 | P25_2M | 316 | P25 | 3.45242 | L5 Cortical excitatory |
| 13 | P25_2M | 300 | P25 | 3.277614 | L5 Cortical excitatory |
| 4 | P25_3F | 355 | P25 | 4.642951 | L5 Cortical excitatory |
| 8 | P25_3F | 408 | P25 | 5.336123 | L5 Cortical excitatory |
| 9 | P25_3F | 364 | P25 | 4.760659 | L5 Cortical excitatory |
| 12 | P25_3F | 297 | P25 | 3.884384 | L5 Cortical excitatory |
| 13 | P25_3F | 245 | P25 | 3.20429 | L5 Cortical excitatory |
| 4 | P60_1M | 432 | P60 | 5.925926 | L5 Cortical excitatory |
| 8 | P60_1M | 382 | P60 | 5.240055 | L5 Cortical excitatory |
| 9 | P60_1M | 329 | P60 | 4.513032 | L5 Cortical excitatory |
| 12 | P60_1M | 277 | P60 | 3.799726 | L5 Cortical excitatory |
| 13 | P60_1M | 253 | P60 | 3.470508 | L5 Cortical excitatory |
| 4 | P60_2F | 378 | P60 | 5.121257 | L5 Cortical excitatory |
| 8 | P60_2F | 375 | P60 | 5.080612 | L5 Cortical excitatory |
| 9 | P60_2F | 362 | P60 | 4.904484 | L5 Cortical excitatory |
| 12 | P60_2F | 236 | P60 | 3.197399 | L5 Cortical excitatory |
| 13 | P60_2F | 230 | P60 | 3.116109 | L5 Cortical excitatory |
| 4 | P60_3M | 456 | P60 | 5.858923 | L5 Cortical excitatory |
| 8 | P60_3M | 324 | P60 | 4.162919 | L5 Cortical excitatory |
| 9 | P60_3M | 384 | P60 | 4.93383 | L5 Cortical excitatory |
| 12 | P60_3M | 244 | P60 | 3.135038 | L5 Cortical excitatory |
| 13 | P60_3M | 222 | P60 | 2.852371 | L5 Cortical excitatory |
| 11 | P15_1M | 378 | P15 | 3.903749 | L6a Cortical excitatory |
| 11 | P15_2F | 284 | P15 | 3.263618 | L6a Cortical excitatory |
| 11 | P15_3F | 465 | P15 | 4.597588 | L6a Cortical excitatory |
| 11 | P20_1M | 364 | P20 | 4.108816 | L6a Cortical excitatory |
| 11 | P20_2M | 298 | P20 | 3.322926 | L6a Cortical excitatory |
| 11 | P20_3M | 288 | P20 | 2.901763 | L6a Cortical excitatory |
| 11 | P25_1M | 280 | P25 | 3.343683 | L6a Cortical excitatory |
| 11 | P25_2M | 302 | P25 | 3.299465 | L6a Cortical excitatory |
| 11 | P25_3F | 275 | P25 | 3.596652 | L6a Cortical excitatory |
| 11 | P60_1M | 238 | P60 | 3.264746 | L6a Cortical excitatory |
| 11 | P60_2F | 254 | P60 | 3.441268 | L6a Cortical excitatory |
| 11 | P60_3M | 264 | P60 | 3.392008 | L6a Cortical excitatory |
| 3 | P15_1M | 686 | P15 | 7.084581 | L6b Cortical excitatory |
| 10 | P15_1M | 328 | P15 | 3.38738 | L6b Cortical excitatory |
| 26 | P15_1M | 43 | P15 | 0.444077 | L6b Cortical excitatory |
| 3 | P15_2F | 538 | P15 | 6.182487 | L6b Cortical excitatory |
| 10 | P15_2F | 330 | P15 | 3.792232 | L6b Cortical excitatory |
| 26 | P15_2F | 61 | P15 | 0.700988 | L6b Cortical excitatory |
| 3 | P15_3F | 659 | P15 | 6.515721 | L6b Cortical excitatory |
| 10 | P15_3F | 344 | P15 | 3.401226 | L6b Cortical excitatory |
| 26 | P15_3F | 52 | P15 | 0.514139 | L6b Cortical excitatory |
| 3 | P20_1M | 662 | P20 | 7.472627 | L6b Cortical excitatory |
| 10 | P20_1M | 283 | P20 | 3.194491 | L6b Cortical excitatory |
| 26 | P20_1M | 42 | P20 | 0.474094 | L6b Cortical excitatory |
| 3 | P20_2M | 581 | P20 | 6.478591 | L6b Cortical excitatory |
| 10 | P20_2M | 350 | P20 | 3.902765 | L6b Cortical excitatory |
| 26 | P20_2M | 87 | P20 | 0.970116 | L6b Cortical excitatory |
| 3 | P20_3M | 531 | P20 | 5.350126 | L6b Cortical excitatory |
| 10 | P20_3M | 353 | P20 | 3.556675 | L6b Cortical excitatory |
| 26 | P20_3M | 54 | P20 | 0.544081 | L6b Cortical excitatory |
| 3 | P25_1M | 628 | P25 | 7.499403 | L6b Cortical excitatory |
| 10 | P25_1M | 330 | P25 | 3.940769 | L6b Cortical excitatory |
| 26 | P25_1M | 95 | P25 | 1.134464 | L6b Cortical excitatory |
| 3 | P25_2M | 622 | P25 | 6.795586 | L6b Cortical excitatory |
| 10 | P25_2M | 295 | P25 | 3.222987 | L6b Cortical excitatory |
| 26 | P25_2M | 84 | P25 | 0.917732 | L6b Cortical excitatory |
| 3 | P25_3F | 537 | P25 | 7.02328 | L6b Cortical excitatory |
| 10 | P25_3F | 294 | P25 | 3.845148 | L6b Cortical excitatory |
| 26 | P25_3F | 66 | P25 | 0.863196 | L6b Cortical excitatory |
| 3 | P60_1M | 552 | P60 | 7.572016 | L6b Cortical excitatory |
| 10 | P60_1M | 301 | P60 | 4.128944 | L6b Cortical excitatory |
| 26 | P60_1M | 63 | P60 | 0.864198 | L6b Cortical excitatory |
| 3 | P60_2F | 625 | P60 | 8.467687 | L6b Cortical excitatory |
| 10 | P60_2F | 315 | P60 | 4.267714 | L6b Cortical excitatory |
| 26 | P60_2F | 66 | P60 | 0.894188 | L6b Cortical excitatory |
| 3 | P60_3M | 557 | P60 | 7.156623 | L6b Cortical excitatory |
| 10 | P60_3M | 313 | P60 | 4.021586 | L6b Cortical excitatory |
| 26 | P60_3M | 71 | P60 | 0.912245 | L6b Cortical excitatory |
| 17 | P15_1M | 121 | P15 | 1.249613 | Mature OLs |
| 17 | P15_2F | 102 | P15 | 1.172144 | Mature OLs |
| 17 | P15_3F | 146 | P15 | 1.443544 | Mature OLs |
| 17 | P20_1M | 151 | P20 | 1.704481 | Mature OLs |
| 17 | P20_2M | 122 | P20 | 1.360393 | Mature OLs |
| 17 | P20_3M | 143 | P20 | 1.440806 | Mature OLs |
| 17 | P25_1M | 250 | P25 | 2.985431 | Mature OLs |
| 17 | P25_2M | 226 | P25 | 2.469136 | Mature OLs |
| 17 | P25_3F | 160 | P25 | 2.092597 | Mature OLs |
| 17 | P60_1M | 224 | P60 | 3.072702 | Mature OLs |
| 17 | P60_2F | 260 | P60 | 3.522558 | Mature OLs |
| 17 | P60_3M | 313 | P60 | 4.021586 | Mature OLs |
| 16 | P15_1M | 175 | P15 | 1.807291 | Meninges |
| 25 | P15_1M | 132 | P15 | 1.363214 | Meninges |
| 16 | P15_2F | 209 | P15 | 2.401747 | Meninges |
| 25 | P15_2F | 69 | P15 | 0.792921 | Meninges |
| 16 | P15_3F | 183 | P15 | 1.809373 | Meninges |
| 25 | P15_3F | 89 | P15 | 0.879968 | Meninges |
| 16 | P20_1M | 174 | P20 | 1.964104 | Meninges |
| 25 | P20_1M | 69 | P20 | 0.778869 | Meninges |
| 16 | P20_2M | 235 | P20 | 2.620428 | Meninges |
| 25 | P20_2M | 84 | P20 | 0.936664 | Meninges |
| 16 | P20_3M | 255 | P20 | 2.56927 | Meninges |
| 25 | P20_3M | 92 | P20 | 0.926952 | Meninges |
| 16 | P25_1M | 188 | P25 | 2.245044 | Meninges |
| 25 | P25_1M | 68 | P25 | 0.812037 | Meninges |
| 16 | P25_2M | 239 | P25 | 2.611166 | Meninges |
| 25 | P25_2M | 110 | P25 | 1.201792 | Meninges |
| 16 | P25_3F | 169 | P25 | 2.210306 | Meninges |
| 25 | P25_3F | 69 | P25 | 0.902433 | Meninges |
| 16 | P60_1M | 149 | P60 | 2.043896 | Meninges |
| 25 | P60_1M | 59 | P60 | 0.809328 | Meninges |
| 16 | P60_2F | 140 | P60 | 1.896762 | Meninges |
| 25 | P60_2F | 50 | P60 | 0.677415 | Meninges |
| 16 | P60_3M | 182 | P60 | 2.33843 | Meninges |
| 25 | P60_3M | 81 | P60 | 1.04073 | Meninges |
| 6 | P15_1M | 506 | P15 | 5.225653 | Microglia |
| 6 | P15_2F | 457 | P15 | 5.251666 | Microglia |
| 6 | P15_3F | 483 | P15 | 4.775559 | Microglia |
| 6 | P20_1M | 487 | P20 | 5.497234 | Microglia |
| 6 | P20_2M | 506 | P20 | 5.642284 | Microglia |
| 6 | P20_3M | 549 | P20 | 5.531486 | Microglia |
| 6 | P25_1M | 383 | P25 | 4.57368 | Microglia |
| 6 | P25_2M | 461 | P25 | 5.0366 | Microglia |
| 6 | P25_3F | 373 | P25 | 4.878368 | Microglia |
| 6 | P60_1M | 333 | P60 | 4.567901 | Microglia |
| 6 | P60_2F | 299 | P60 | 4.050942 | Microglia |
| 6 | P60_3M | 335 | P60 | 4.304253 | Microglia |
| 7 | P15_1M | 625 | P15 | 6.454611 | OPCs |
| 7 | P15_2F | 547 | P15 | 6.285911 | OPCs |
| 7 | P15_3F | 635 | P15 | 6.278426 | OPCs |
| 7 | P20_1M | 420 | P20 | 4.740941 | OPCs |
| 7 | P20_2M | 403 | P20 | 4.493756 | OPCs |
| 7 | P20_3M | 496 | P20 | 4.997481 | OPCs |
| 7 | P25_1M | 358 | P25 | 4.275137 | OPCs |
| 7 | P25_2M | 409 | P25 | 4.46848 | OPCs |
| 7 | P25_3F | 308 | P25 | 4.02825 | OPCs |
| 7 | P60_1M | 218 | P60 | 2.990398 | OPCs |
| 7 | P60_2F | 231 | P60 | 3.129657 | OPCs |
| 7 | P60_3M | 258 | P60 | 3.314917 | OPCs |
| 24 | P15_1M | 88 | P15 | 0.908809 | Other INs |
| 27 | P15_1M | 45 | P15 | 0.464732 | Other INs |
| 24 | P15_2F | 90 | P15 | 1.034245 | Other INs |
| 27 | P15_2F | 35 | P15 | 0.402206 | Other INs |
| 24 | P15_3F | 110 | P15 | 1.087601 | Other INs |
| 27 | P15_3F | 53 | P15 | 0.524026 | Other INs |
| 24 | P20_1M | 85 | P20 | 0.959476 | Other INs |
| 27 | P20_1M | 43 | P20 | 0.485382 | Other INs |
| 24 | P20_2M | 93 | P20 | 1.037021 | Other INs |
| 27 | P20_2M | 36 | P20 | 0.401427 | Other INs |
| 24 | P20_3M | 93 | P20 | 0.937028 | Other INs |
| 27 | P20_3M | 47 | P20 | 0.473552 | Other INs |
| 24 | P25_1M | 78 | P25 | 0.931455 | Other INs |
| 27 | P25_1M | 29 | P25 | 0.34631 | Other INs |
| 24 | P25_2M | 79 | P25 | 0.863105 | Other INs |
| 27 | P25_2M | 33 | P25 | 0.360538 | Other INs |
| 24 | P25_3F | 70 | P25 | 0.915511 | Other INs |
| 27 | P25_3F | 31 | P25 | 0.405441 | Other INs |
| 24 | P60_1M | 61 | P60 | 0.836763 | Other INs |
| 27 | P60_1M | 36 | P60 | 0.493827 | Other INs |
| 24 | P60_2F | 84 | P60 | 1.138057 | Other INs |
| 27 | P60_2F | 31 | P60 | 0.419997 | Other INs |
| 24 | P60_3M | 81 | P60 | 1.04073 | Other INs |
| 27 | P60_3M | 32 | P60 | 0.411153 | Other INs |
| 18 | P15_1M | 199 | P15 | 2.055148 | Pericytes |
| 20 | P15_1M | 150 | P15 | 1.549107 | Pericytes |
| 18 | P15_2F | 186 | P15 | 2.13744 | Pericytes |
| 20 | P15_2F | 175 | P15 | 2.011032 | Pericytes |
| 18 | P15_3F | 202 | P15 | 1.997232 | Pericytes |
| 20 | P15_3F | 103 | P15 | 1.01839 | Pericytes |
| 18 | P20_1M | 175 | P20 | 1.975392 | Pericytes |
| 20 | P20_1M | 114 | P20 | 1.286827 | Pericytes |
| 18 | P20_2M | 178 | P20 | 1.984835 | Pericytes |
| 20 | P20_2M | 243 | P20 | 2.709634 | Pericytes |
| 18 | P20_3M | 191 | P20 | 1.924433 | Pericytes |
| 20 | P20_3M | 220 | P20 | 2.216625 | Pericytes |
| 18 | P25_1M | 172 | P25 | 2.053977 | Pericytes |
| 20 | P25_1M | 160 | P25 | 1.910676 | Pericytes |
| 18 | P25_2M | 211 | P25 | 2.305255 | Pericytes |
| 20 | P25_2M | 146 | P25 | 1.595105 | Pericytes |
| 18 | P25_3F | 141 | P25 | 1.844101 | Pericytes |
| 20 | P25_3F | 144 | P25 | 1.883338 | Pericytes |
| 18 | P60_1M | 126 | P60 | 1.728395 | Pericytes |
| 20 | P60_1M | 170 | P60 | 2.331962 | Pericytes |
| 18 | P60_2F | 147 | P60 | 1.9916 | Pericytes |
| 20 | P60_2F | 108 | P60 | 1.463216 | Pericytes |
| 18 | P60_3M | 165 | P60 | 2.120005 | Pericytes |
| 20 | P60_3M | 152 | P60 | 1.952974 | Pericytes |
| 14 | P15_1M | 262 | P15 | 2.705773 | Pvalb Ins |
| 14 | P15_2F | 232 | P15 | 2.666054 | Pvalb Ins |
| 14 | P15_3F | 251 | P15 | 2.481709 | Pvalb Ins |
| 14 | P20_1M | 213 | P20 | 2.404335 | Pvalb Ins |
| 14 | P20_2M | 237 | P20 | 2.64273 | Pvalb Ins |
| 14 | P20_3M | 267 | P20 | 2.690176 | Pvalb Ins |
| 14 | P25_1M | 223 | P25 | 2.663005 | Pvalb Ins |
| 14 | P25_2M | 274 | P25 | 2.993554 | Pvalb Ins |
| 14 | P25_3F | 200 | P25 | 2.615747 | Pvalb Ins |
| 14 | P60_1M | 208 | P60 | 2.853224 | Pvalb Ins |
| 14 | P60_2F | 226 | P60 | 3.061916 | Pvalb Ins |
| 14 | P60_3M | 204 | P60 | 2.621097 | Pvalb Ins |
| 21 | P15_1M | 183 | P15 | 1.88991 | Sst INs |
| 21 | P15_2F | 161 | P15 | 1.850149 | Sst INs |
| 21 | P15_3F | 187 | P15 | 1.848922 | Sst INs |
| 21 | P20_1M | 174 | P20 | 1.964104 | Sst INs |
| 21 | P20_2M | 163 | P20 | 1.817574 | Sst INs |
| 21 | P20_3M | 141 | P20 | 1.420655 | Sst INs |
| 21 | P25_1M | 150 | P25 | 1.791259 | Sst INs |
| 21 | P25_2M | 145 | P25 | 1.58418 | Sst INs |
| 21 | P25_3F | 133 | P25 | 1.739472 | Sst INs |
| 21 | P60_1M | 127 | P60 | 1.742112 | Sst INs |
| 21 | P60_2F | 122 | P60 | 1.652893 | Sst INs |
| 21 | P60_3M | 132 | P60 | 1.696004 | Sst INs |
| 22 | P15_1M | 111 | P15 | 1.146339 | Vip INs |
| 22 | P15_2F | 99 | P15 | 1.13767 | Vip INs |
| 22 | P15_3F | 133 | P15 | 1.315009 | Vip INs |
| 22 | P20_1M | 110 | P20 | 1.241675 | Vip INs |
| 22 | P20_2M | 94 | P20 | 1.048171 | Vip INs |
| 22 | P20_3M | 137 | P20 | 1.380353 | Vip INs |
| 22 | P25_1M | 108 | P25 | 1.289706 | Vip INs |
| 22 | P25_2M | 107 | P25 | 1.169016 | Vip INs |
| 22 | P25_3F | 82 | P25 | 1.072456 | Vip INs |
| 22 | P60_1M | 75 | P60 | 1.028807 | Vip INs |
| 22 | P60_2F | 91 | P60 | 1.232895 | Vip INs |
| 22 | P60_3M | 87 | P60 | 1.117821 | Vip INs |

**Extended Data Table 4.**
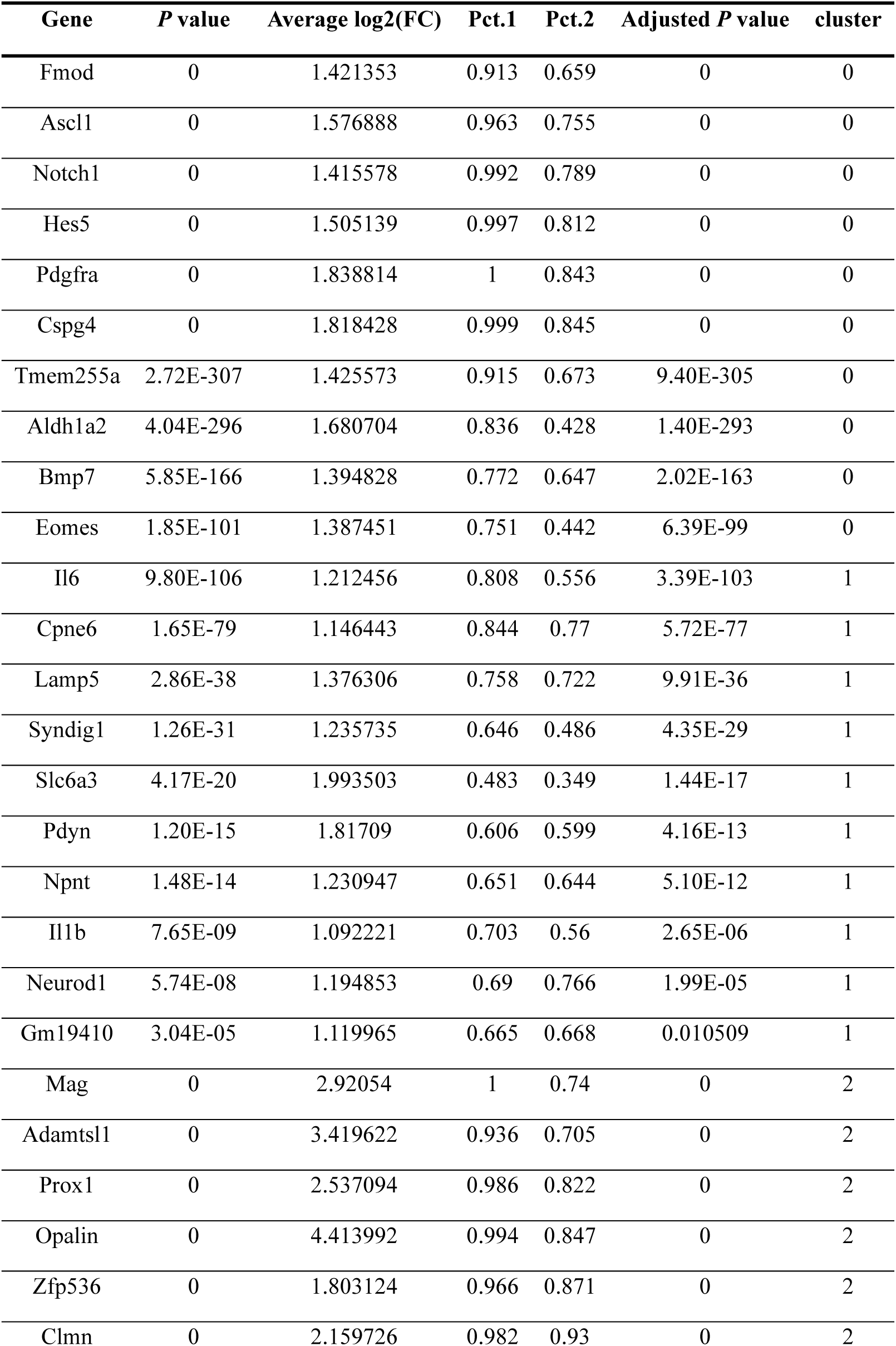
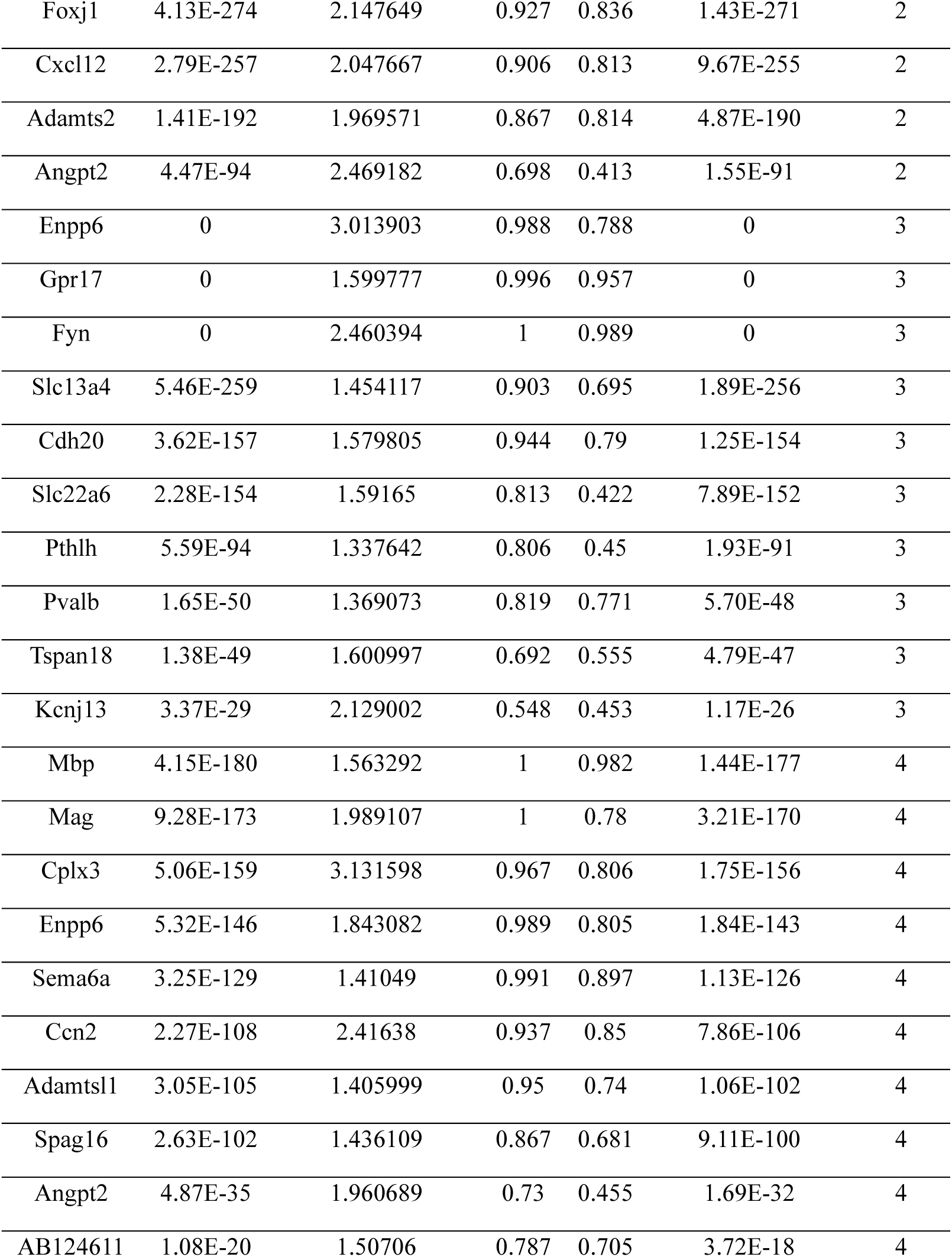
Top 10 marker genes for each cluster identified in Oligodendrocyte subset. (Pct.1: percentage of cells where the gene is detected in the target cluster, Pct.2: percentage of cells where the gene is detected in all other clusters, FC: Fold change)

**Extended Data Table 5.**
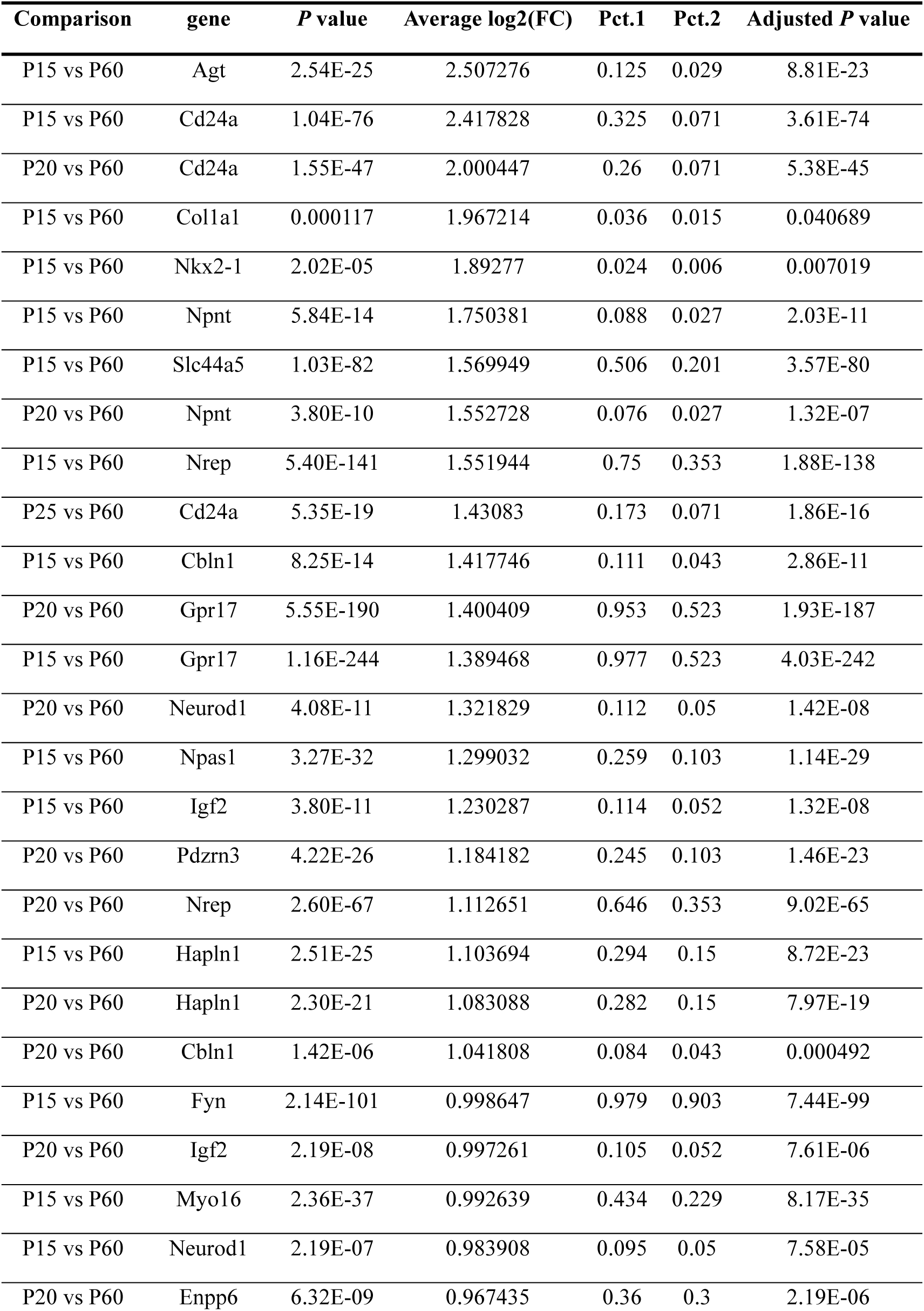
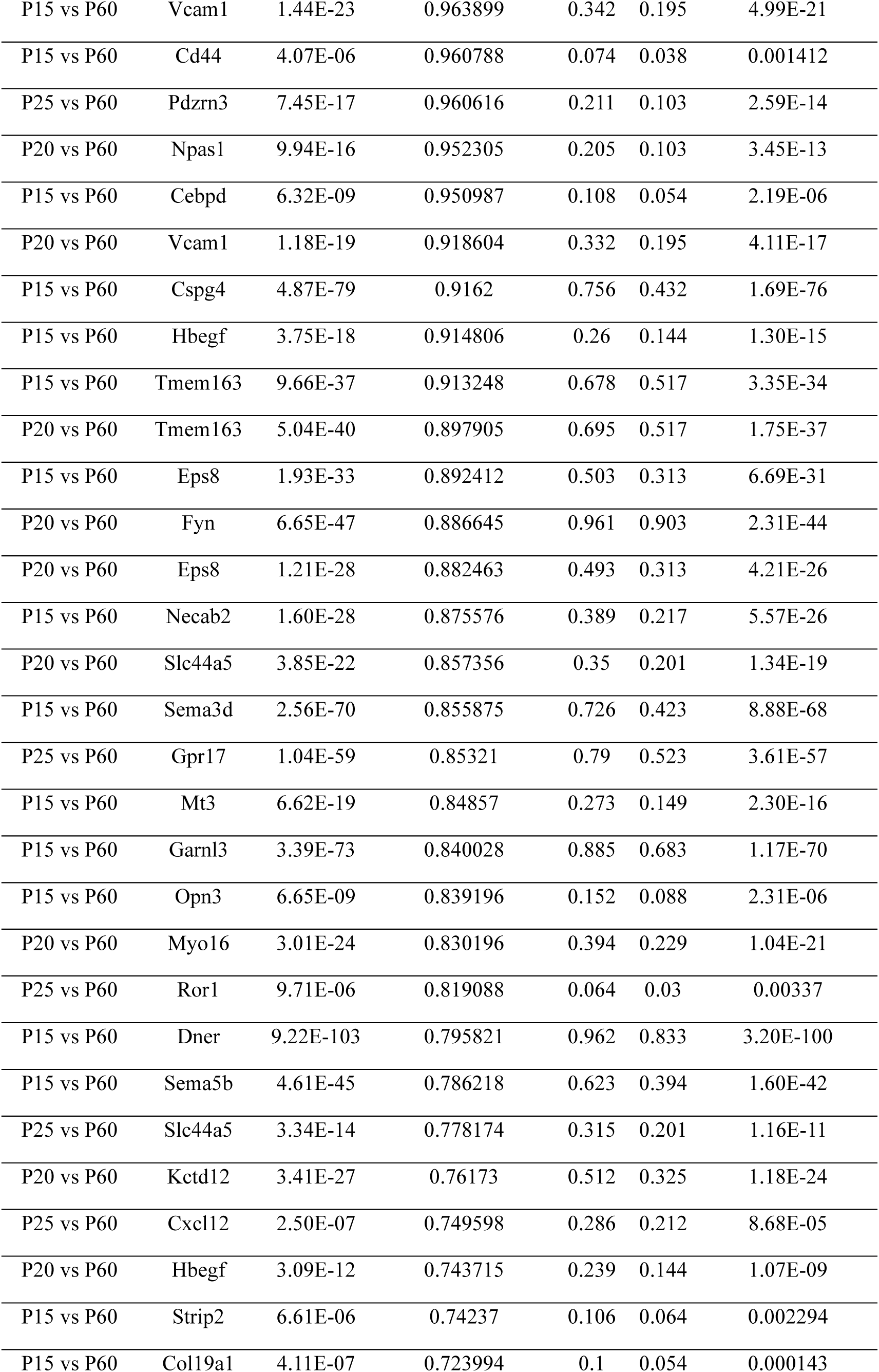
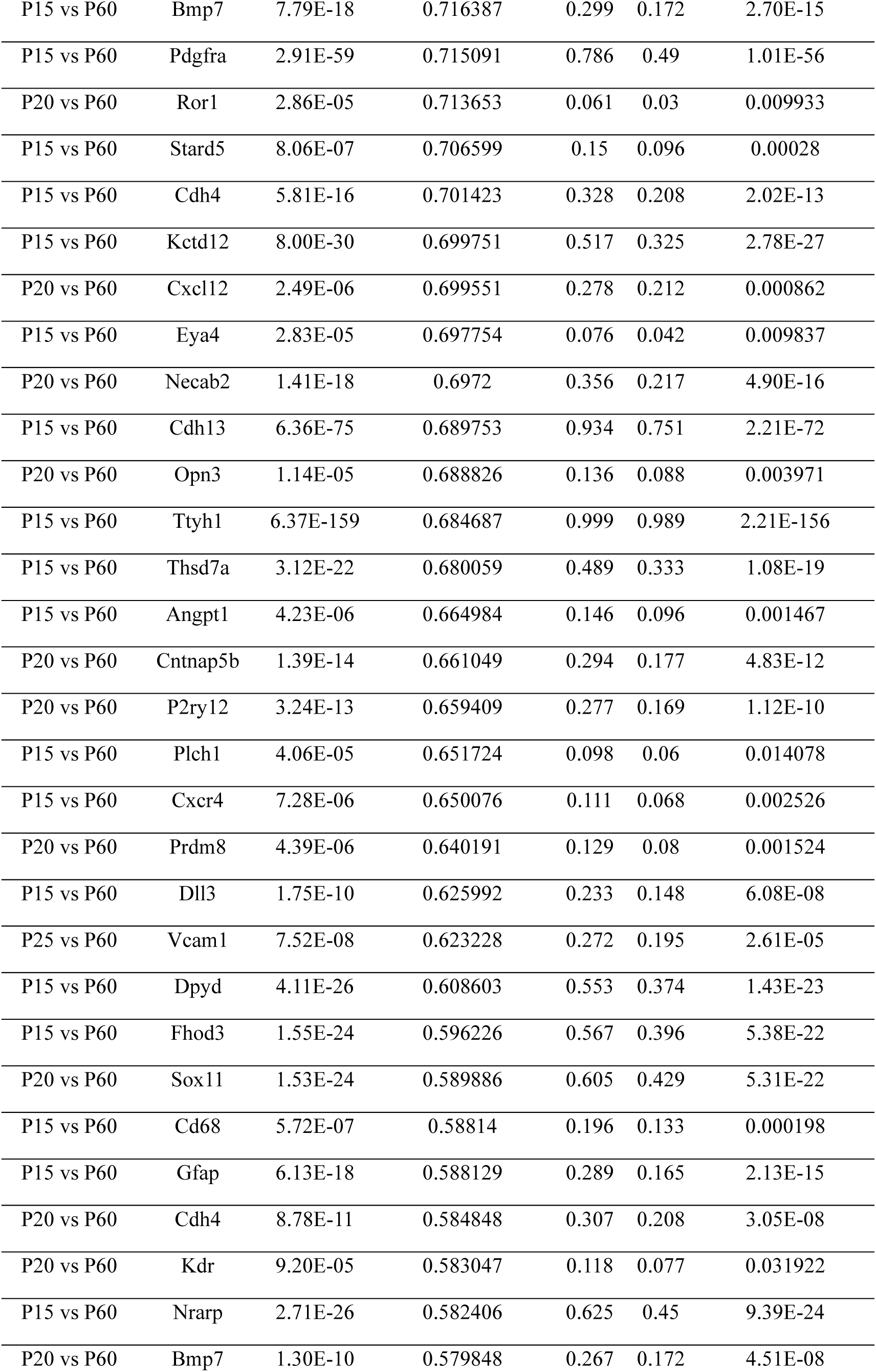
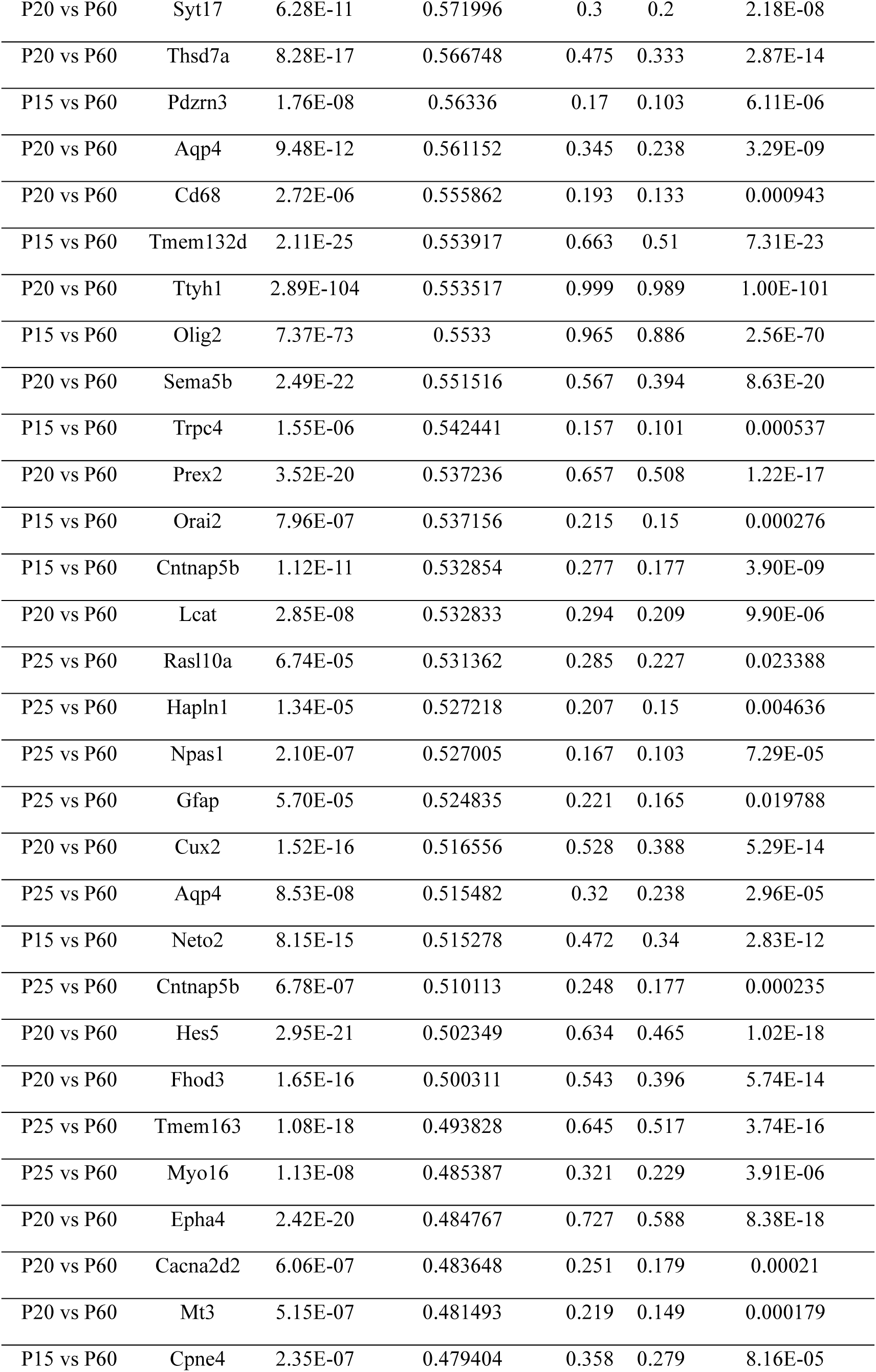
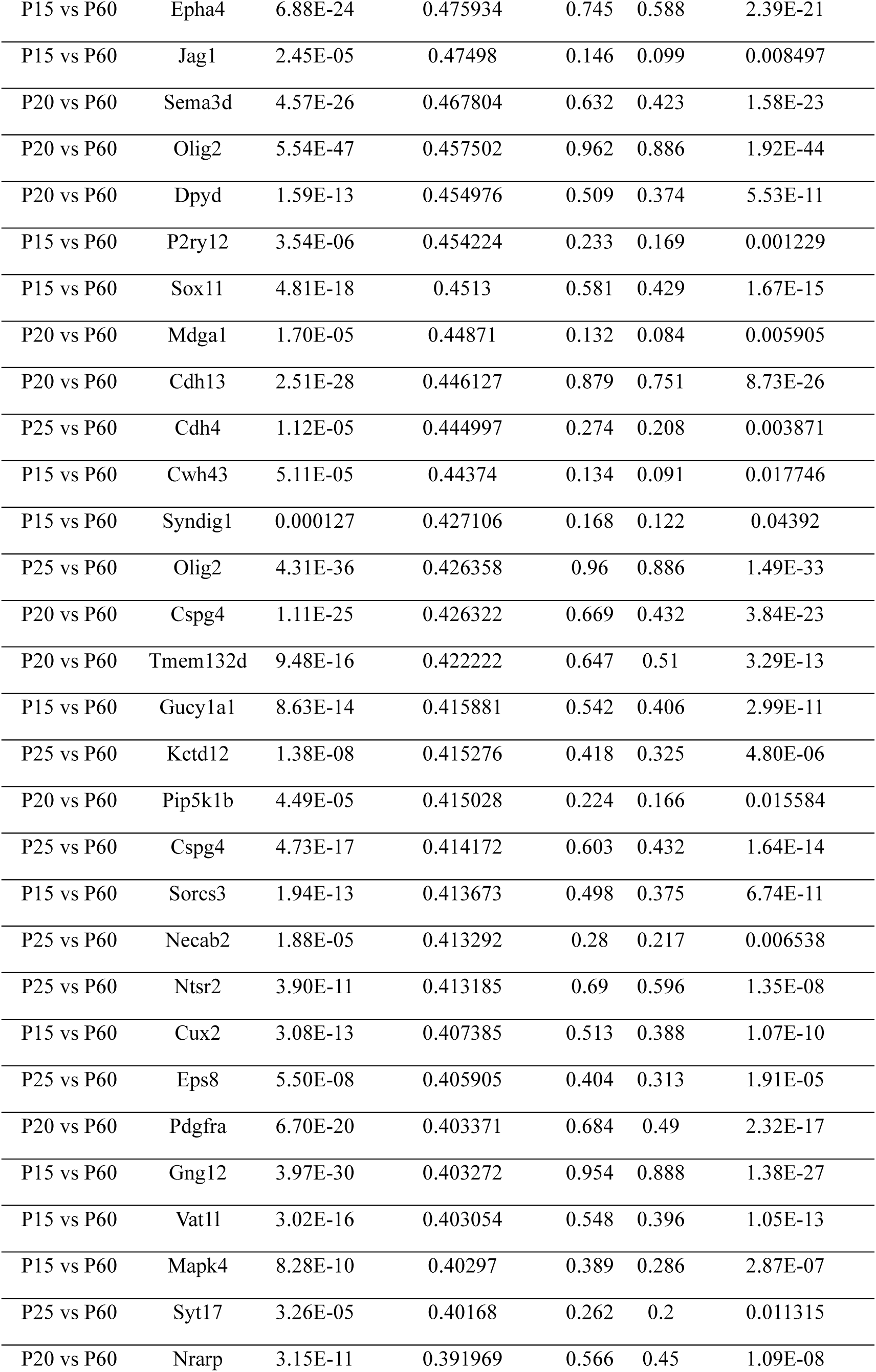
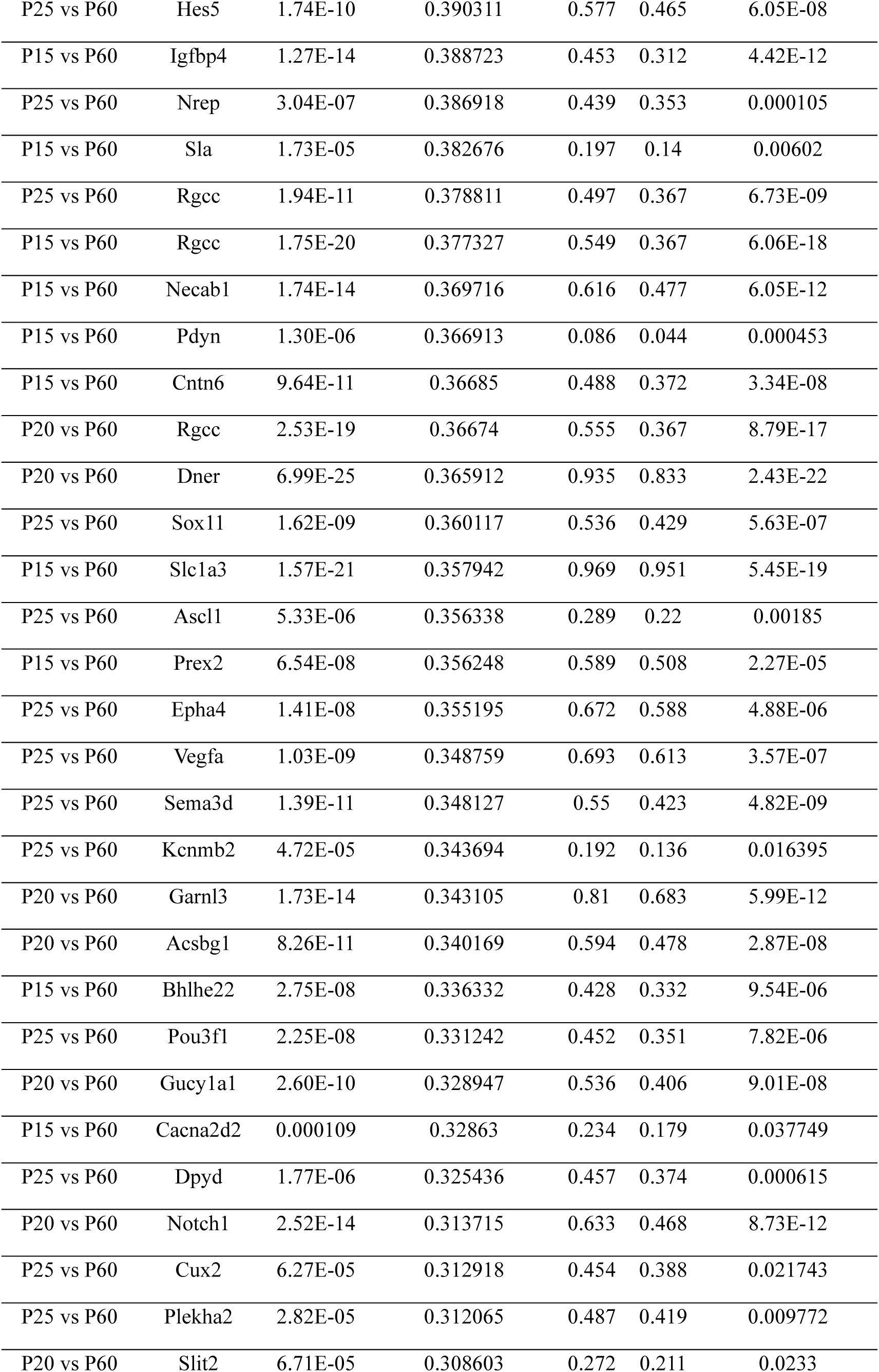
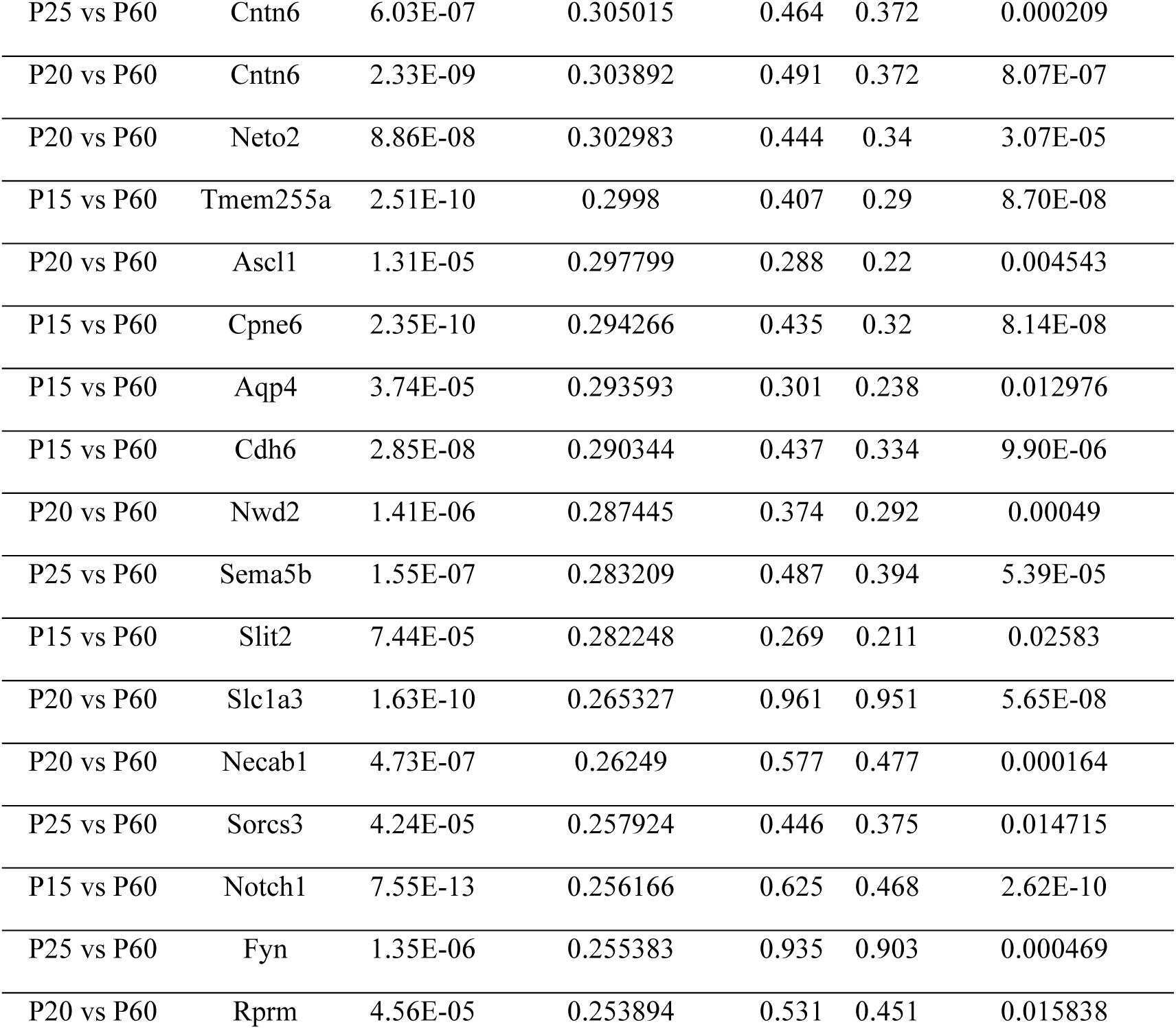
List of upregulated genes (enriched in younger ages) detected in differential gene expression analysis between the young ages (P15, P20, or P25) and adults (P60). (Pct.1: percentage of cells where the gene is detected in the younger age, Pct.2: percentage of cells where the gene is detected in in adult age, FC: Fold change)

**Extended Data Table 6.** List of downregulated genes (enriched in adult ages) detected in differential gene expression analysis between the young ages (P15, P20, or P25) and adults (P60). (Pct.1: percentage of cells where the gene is detected in the younger age, Pct.2: percentage of cells where the gene is detected in in adult age, FC: Fold change)

| Comparison | Gene | *P* value | Average log2(FC) | Pct.1 | Pct.2 | Adjusted *P* value |
| --- | --- | --- | --- | --- | --- | --- |
| P15 vs P60 | Adamtsl1 | 1.17E-116 | -3.35792 | 0.049 | 0.316 | 4.05E-114 |
| P15 vs P60 | Opalin | 1.02E-197 | -3.31271 | 0.069 | 0.478 | 3.55E-195 |
| P15 vs P60 | Igfbp6 | 8.90E-61 | -3.10569 | 0.031 | 0.186 | 3.09E-58 |
| P15 vs P60 | Arc | 1.03E-225 | -2.67737 | 0.296 | 0.731 | 3.59E-223 |
| P15 vs P60 | Osm | 2.66E-08 | -2.40228 | 0.005 | 0.025 | 9.22E-06 |
| P15 vs P60 | Chrm2 | 4.98E-43 | -2.10866 | 0.061 | 0.203 | 1.73E-40 |
| P15 vs P60 | Fos | 1.78E-30 | -2.00299 | 0.065 | 0.18 | 6.17E-28 |
| P25 vs P60 | Fos | 2.29E-24 | -1.94983 | 0.067 | 0.18 | 7.95E-22 |
| P20 vs P60 | Arc | 2.40E-128 | -1.94189 | 0.437 | 0.731 | 8.32E-126 |
| P15 vs P60 | Adamts2 | 1.65E-32 | -1.8695 | 0.072 | 0.196 | 5.72E-30 |
| P20 vs P60 | Thbs4 | 4.11E-08 | -1.73485 | 0.012 | 0.041 | 1.42E-05 |
| P25 vs P60 | Arc | 3.82E-95 | -1.65333 | 0.484 | 0.731 | 1.32E-92 |
| P20 vs P60 | Adamtsl1 | 2.76E-49 | -1.65298 | 0.115 | 0.316 | 9.59E-47 |
| P15 vs P60 | Rxfp1 | 4.48E-11 | -1.6309 | 0.039 | 0.089 | 1.55E-08 |
| P15 vs P60 | Clmn | 3.89E-103 | -1.61588 | 0.431 | 0.671 | 1.35E-100 |
| P15 vs P60 | Ccn2 | 2.07E-20 | -1.5896 | 0.157 | 0.266 | 7.19E-18 |
| P20 vs P60 | Opalin | 6.22E-82 | -1.58377 | 0.175 | 0.478 | 2.16E-79 |
| P15 vs P60 | Arhgap6 | 4.49E-10 | -1.57289 | 0.033 | 0.077 | 1.56E-07 |
| P15 vs P60 | Foxj1 | 5.54E-43 | -1.53473 | 0.093 | 0.257 | 1.92E-40 |
| P15 vs P60 | Gfra2 | 2.01E-130 | -1.51861 | 0.403 | 0.733 | 6.97E-128 |
| P20 vs P60 | Fos | 1.04E-15 | -1.47624 | 0.091 | 0.18 | 3.61E-13 |
| P15 vs P60 | Pkib | 1.94E-10 | -1.44397 | 0.034 | 0.08 | 6.72E-08 |
| P20 vs P60 | Chrm2 | 2.38E-21 | -1.35981 | 0.093 | 0.203 | 8.25E-19 |
| P15 vs P60 | Prox1 | 4.07E-59 | -1.34763 | 0.312 | 0.523 | 1.41E-56 |
| P15 vs P60 | Carmn | 2.07E-05 | -1.33235 | 0.017 | 0.038 | 0.007196 |
| P15 vs P60 | Mag | 5.16E-78 | -1.2762 | 0.308 | 0.57 | 1.79E-75 |
| P20 vs P60 | Clmn | 3.21E-62 | -1.24524 | 0.495 | 0.671 | 1.11E-59 |
| P15 vs P60 | Dpy19l1 | 1.40E-136 | -1.15519 | 0.839 | 0.939 | 4.87E-134 |
| P15 vs P60 | Zfp536 | 6.44E-58 | -1.13962 | 0.453 | 0.637 | 2.24E-55 |
| P15 vs P60 | Prph | 5.48E-17 | -1.13857 | 0.119 | 0.216 | 1.90E-14 |
| P20 vs P60 | Ccn2 | 2.59E-10 | -1.11553 | 0.185 | 0.266 | 8.98E-08 |
| P15 vs P60 | Rims3 | 2.55E-33 | -1.10359 | 0.304 | 0.459 | 8.86E-31 |
| P15 vs P60 | Deptor | 3.28E-12 | -1.08781 | 0.078 | 0.147 | 1.14E-09 |
| P15 vs P60 | Tanc1 | 5.87E-32 | -1.05172 | 0.256 | 0.418 | 2.04E-29 |
| P20 vs P60 | Igfbp6 | 1.37E-10 | -1.02898 | 0.112 | 0.186 | 4.76E-08 |
| P15 vs P60 | Rmst | 3.60E-26 | -1.00485 | 0.225 | 0.369 | 1.25E-23 |
| P15 vs P60 | Lifr | 1.54E-40 | -0.99386 | 0.307 | 0.497 | 5.34E-38 |
| P15 vs P60 | Cntnap4 | 7.51E-16 | -0.97744 | 0.277 | 0.369 | 2.61E-13 |
| P15 vs P60 | Dkk3 | 9.39E-106 | -0.97384 | 0.74 | 0.911 | 3.26E-103 |
| P20 vs P60 | Vwc2l | 1.47E-11 | -0.97304 | 0.08 | 0.153 | 5.09E-09 |
| P15 vs P60 | Gjc3 | 1.01E-129 | -0.97112 | 0.729 | 0.917 | 3.51E-127 |
| P15 vs P60 | Ndst3 | 2.86E-17 | -0.94567 | 0.196 | 0.305 | 9.91E-15 |
| P20 vs P60 | Prph | 7.79E-11 | -0.91331 | 0.136 | 0.216 | 2.70E-08 |
| P15 vs P60 | Cabp7 | 1.15E-27 | -0.90935 | 0.311 | 0.462 | 3.99E-25 |
| P15 vs P60 | Hs3st2 | 4.73E-24 | -0.90665 | 0.372 | 0.495 | 1.64E-21 |
| P15 vs P60 | Ly6a | 7.45E-05 | -0.89605 | 0.064 | 0.098 | 0.025838 |
| P15 vs P60 | Mbp | 3.98E-66 | -0.89107 | 0.808 | 0.906 | 1.38E-63 |
| P20 vs P60 | Zfp536 | 1.34E-36 | -0.88442 | 0.483 | 0.637 | 4.66E-34 |
| P20 vs P60 | Dkk3 | 8.19E-86 | -0.88264 | 0.742 | 0.911 | 2.84E-83 |
| P15 vs P60 | Igflr | 1.21E-42 | -0.8782 | 0.498 | 0.65 | 4.21E-40 |
| P15 vs P60 | Rbp4 | 7.58E-08 | -0.86738 | 0.11 | 0.168 | 2.63E-05 |
| P20 vs P60 | Igflr | 4.76E-39 | -0.84321 | 0.492 | 0.65 | 1.65E-36 |
| P15 vs P60 | Arhgap25 | 8.61E-11 | -0.83624 | 0.161 | 0.239 | 2.99E-08 |
| P20 vs P60 | Rims3 | 5.74E-18 | -0.83123 | 0.351 | 0.459 | 1.99E-15 |
| P15 vs P60 | Vwc2l | 2.12E-08 | -0.81213 | 0.094 | 0.153 | 7.36E-06 |
| P20 vs P60 | Tanc1 | 2.05E-17 | -0.79482 | 0.299 | 0.418 | 7.13E-15 |
| P20 vs P60 | Gfra2 | 4.89E-42 | -0.77749 | 0.571 | 0.733 | 1.70E-39 |
| P20 vs P60 | Prox1 | 1.40E-28 | -0.77658 | 0.359 | 0.523 | 4.87E-26 |
| P25 vs P60 | Vwc2l | 3.20E-07 | -0.75962 | 0.095 | 0.153 | 0.000111 |
| P15 vs P60 | Ppp1r1b | 3.59E-13 | -0.75713 | 0.215 | 0.312 | 1.25E-10 |
| P15 vs P60 | Aif1 | 2.13E-05 | -0.74959 | 0.054 | 0.089 | 0.007395 |
| P20 vs P60 | Lifr | 1.10E-19 | -0.71704 | 0.374 | 0.497 | 3.80E-17 |
| P20 vs P60 | Mag | 6.32E-39 | -0.71083 | 0.335 | 0.57 | 2.19E-36 |
| P20 vs P60 | Foxp2 | 2.78E-09 | -0.69758 | 0.316 | 0.389 | 9.66E-07 |
| P20 vs P60 | Dpy19l1 | 5.77E-55 | -0.69254 | 0.897 | 0.939 | 2.00E-52 |
| P15 vs P60 | Foxp2 | 2.63E-08 | -0.69195 | 0.328 | 0.389 | 9.13E-06 |
| P15 vs P60 | Tmem119 | 4.21E-11 | -0.68008 | 0.164 | 0.248 | 1.46E-08 |
| P20 vs P60 | Foxj1 | 1.10E-11 | -0.67485 | 0.164 | 0.257 | 3.82E-09 |
| P20 vs P60 | Cntnap4 | 3.16E-08 | -0.67462 | 0.305 | 0.369 | 1.10E-05 |
| P25 vs P60 | Gadd45a | 1.46E-05 | -0.67009 | 0.204 | 0.261 | 0.005055 |
| P15 vs P60 | Rspo2 | 4.69E-10 | -0.66925 | 0.217 | 0.298 | 1.63E-07 |
| P20 vs P60 | Gjc3 | 1.90E-76 | -0.66365 | 0.735 | 0.917 | 6.58E-74 |
| P15 vs P60 | Cdh9 | 1.17E-05 | -0.66056 | 0.134 | 0.183 | 0.004063 |
| P25 vs P60 | Nr4a1 | 1.19E-15 | -0.64562 | 0.563 | 0.653 | 4.13E-13 |
| P25 vs P60 | Cntnap4 | 3.05E-08 | -0.64394 | 0.298 | 0.369 | 1.06E-05 |
| P20 vs P60 | Arhgap25 | 1.03E-07 | -0.64145 | 0.169 | 0.239 | 3.57E-05 |
| P25 vs P60 | Bdnf | 7.74E-06 | -0.63267 | 0.13 | 0.186 | 0.002685 |
| P15 vs P60 | Sema3e | 2.51E-07 | -0.63027 | 0.159 | 0.222 | 8.71E-05 |
| P25 vs P60 | Tanc1 | 6.44E-13 | -0.62478 | 0.309 | 0.418 | 2.23E-10 |
| P20 vs P60 | Ndst3 | 9.30E-08 | -0.62179 | 0.232 | 0.305 | 3.23E-05 |
| P15 vs P60 | Rasl10a | 2.38E-05 | -0.60754 | 0.18 | 0.227 | 0.008252 |
| P15 vs P60 | Fign | 7.03E-07 | -0.59679 | 0.162 | 0.224 | 0.000244 |
| P15 vs P60 | Nrp2 | 2.70E-06 | -0.59674 | 0.162 | 0.218 | 0.000938 |
| P20 vs P60 | Rmst | 3.55E-08 | -0.58667 | 0.298 | 0.369 | 1.23E-05 |
| P20 vs P60 | Mbp | 2.42E-26 | -0.5774 | 0.883 | 0.906 | 8.41E-24 |
| P20 vs P60 | Ppp1r1b | 1.24E-06 | -0.57054 | 0.247 | 0.312 | 0.000429 |
| P25 vs P60 | Adamtsl1 | 4.93E-08 | -0.56942 | 0.236 | 0.316 | 1.71E-05 |
| P20 vs P60 | Nr4a1 | 2.38E-09 | -0.54868 | 0.626 | 0.653 | 8.25E-07 |
| P25 vs P60 | Opalin | 5.63E-18 | -0.54718 | 0.323 | 0.478 | 1.95E-15 |
| P25 vs P60 | Grasp | 3.67E-08 | -0.53944 | 0.341 | 0.421 | 1.27E-05 |
| P20 vs P60 | Tmem119 | 2.40E-05 | -0.52254 | 0.192 | 0.248 | 0.008328 |
| P15 vs P60 | Hat1 | 3.47E-08 | -0.51974 | 0.363 | 0.426 | 1.20E-05 |
| P25 vs P60 | Ndst3 | 7.02E-07 | -0.51713 | 0.232 | 0.305 | 0.000243 |
| P20 vs P60 | Hs3st2 | 4.40E-07 | -0.50489 | 0.444 | 0.495 | 0.000153 |
| P25 vs P60 | Igflr | 5.48E-15 | -0.49299 | 0.57 | 0.65 | 1.90E-12 |
| P25 vs P60 | Rims3 | 1.10E-07 | -0.48257 | 0.387 | 0.459 | 3.81E-05 |
| P15 vs P60 | Bhlhe40 | 3.56E-09 | -0.45136 | 0.443 | 0.517 | 1.23E-06 |
| P20 vs P60 | Cobll1 | 4.46E-11 | -0.44881 | 0.455 | 0.542 | 1.55E-08 |
| P25 vs P60 | Dkk3 | 7.60E-27 | -0.42777 | 0.83 | 0.911 | 2.64E-24 |
| P25 vs P60 | Clmn | 1.64E-10 | -0.41957 | 0.631 | 0.671 | 5.68E-08 |
| P15 vs P60 | Gsta4 | 3.26E-07 | -0.41716 | 0.347 | 0.412 | 0.000113 |
| P25 vs P60 | Gfra2 | 1.44E-13 | -0.4126 | 0.656 | 0.733 | 5.01E-11 |
| P15 vs P60 | Slc38a1 | 2.83E-15 | -0.41114 | 0.61 | 0.7 | 9.81E-13 |
| P20 vs P60 | Nxph3 | 3.74E-09 | -0.40347 | 0.422 | 0.507 | 1.30E-06 |
| P20 vs P60 | Gsta4 | 9.59E-06 | -0.39426 | 0.356 | 0.412 | 0.003329 |
| P25 vs P60 | Lifr | 8.15E-07 | -0.38277 | 0.431 | 0.497 | 0.000283 |
| P15 vs P60 | Vegfa | 8.77E-05 | -0.37386 | 0.598 | 0.613 | 0.030425 |
| P20 vs P60 | Bhlhe40 | 3.29E-05 | -0.35569 | 0.464 | 0.517 | 0.011429 |
| P25 vs P60 | Zfp536 | 1.79E-08 | -0.35106 | 0.559 | 0.637 | 6.22E-06 |
| P15 vs P60 | Satb2 | 0.000112 | -0.34435 | 0.4 | 0.443 | 0.03899 |
| P15 vs P60 | Nxph3 | 5.45E-10 | -0.34408 | 0.416 | 0.507 | 1.89E-07 |
| P15 vs P60 | Ntsr2 | 2.44E-06 | -0.32188 | 0.558 | 0.596 | 0.000847 |
| P15 vs P60 | Il6st | 1.58E-07 | -0.29822 | 0.645 | 0.68 | 5.49E-05 |
| P25 vs P60 | Mbp | 2.33E-10 | -0.29693 | 0.883 | 0.906 | 8.10E-08 |
| P25 vs P60 | Cdh20 | 4.68E-07 | -0.29216 | 0.508 | 0.579 | 0.000162 |
| P20 vs P60 | Arhgef28 | 4.54E-09 | -0.26649 | 0.524 | 0.612 | 1.57E-06 |
| P25 vs P60 | Dpy1911 | 1.79E-10 | -0.25473 | 0.91 | 0.939 | 6.22E-08 |

**Extended Data Table 7.**
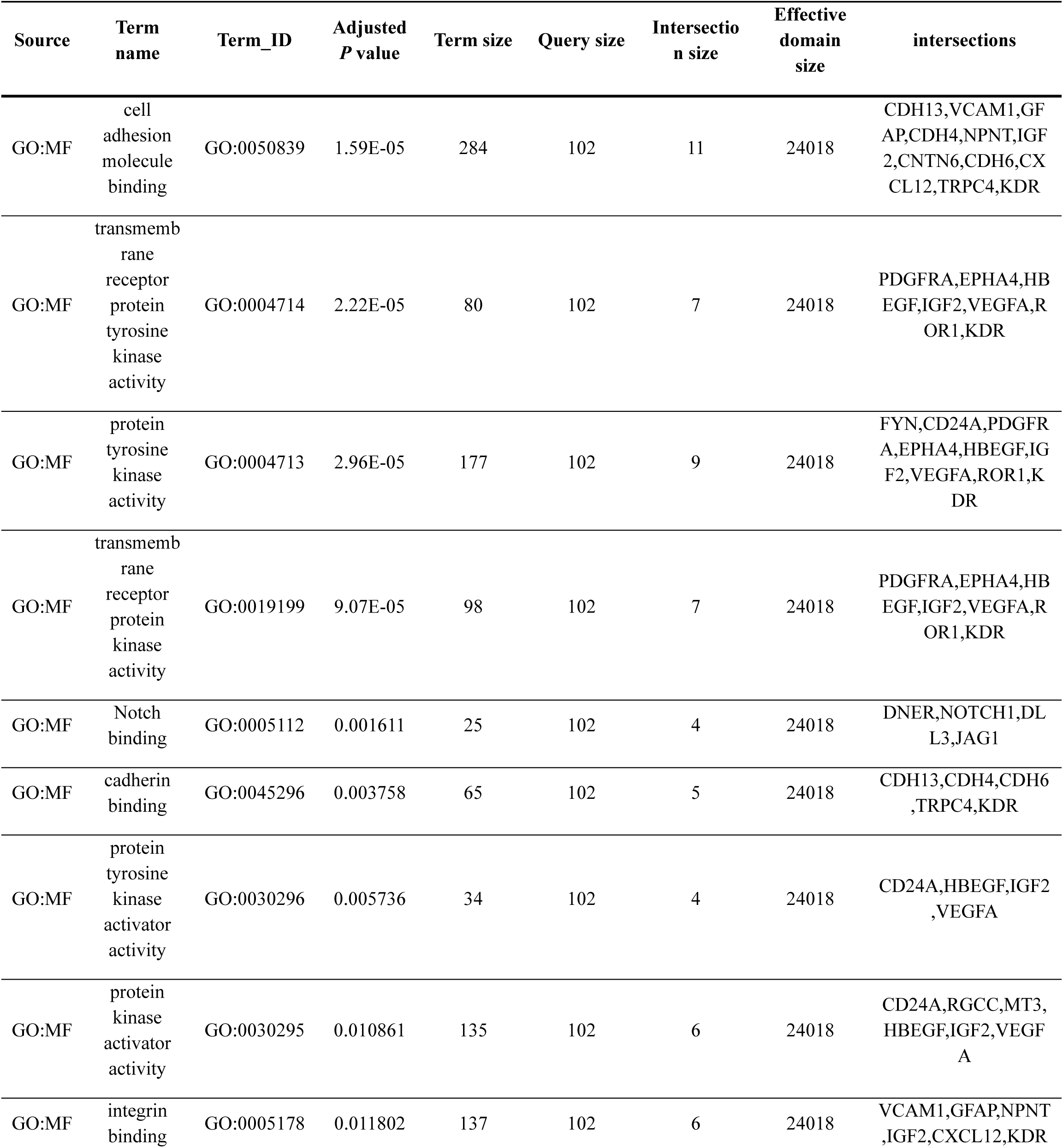
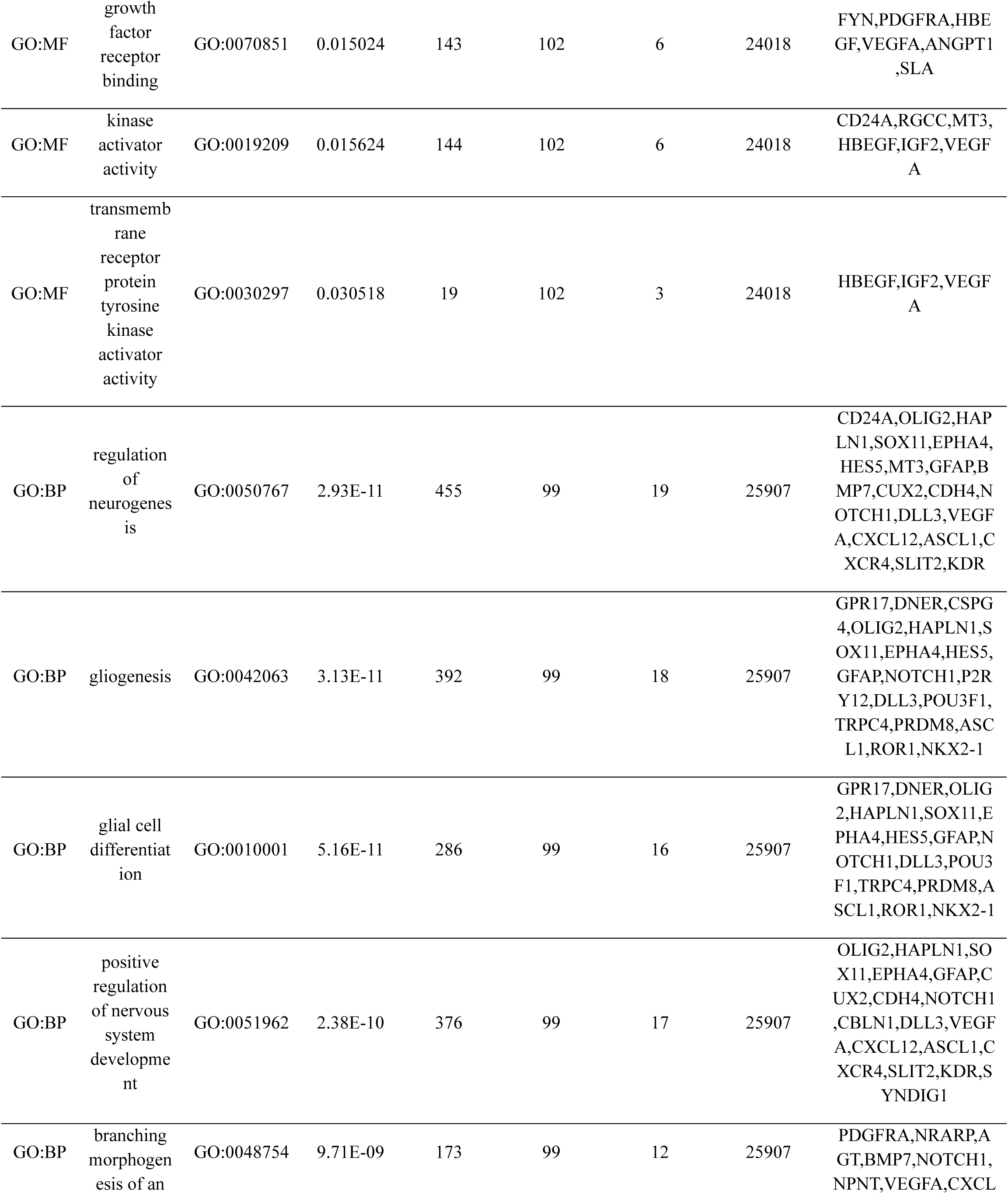
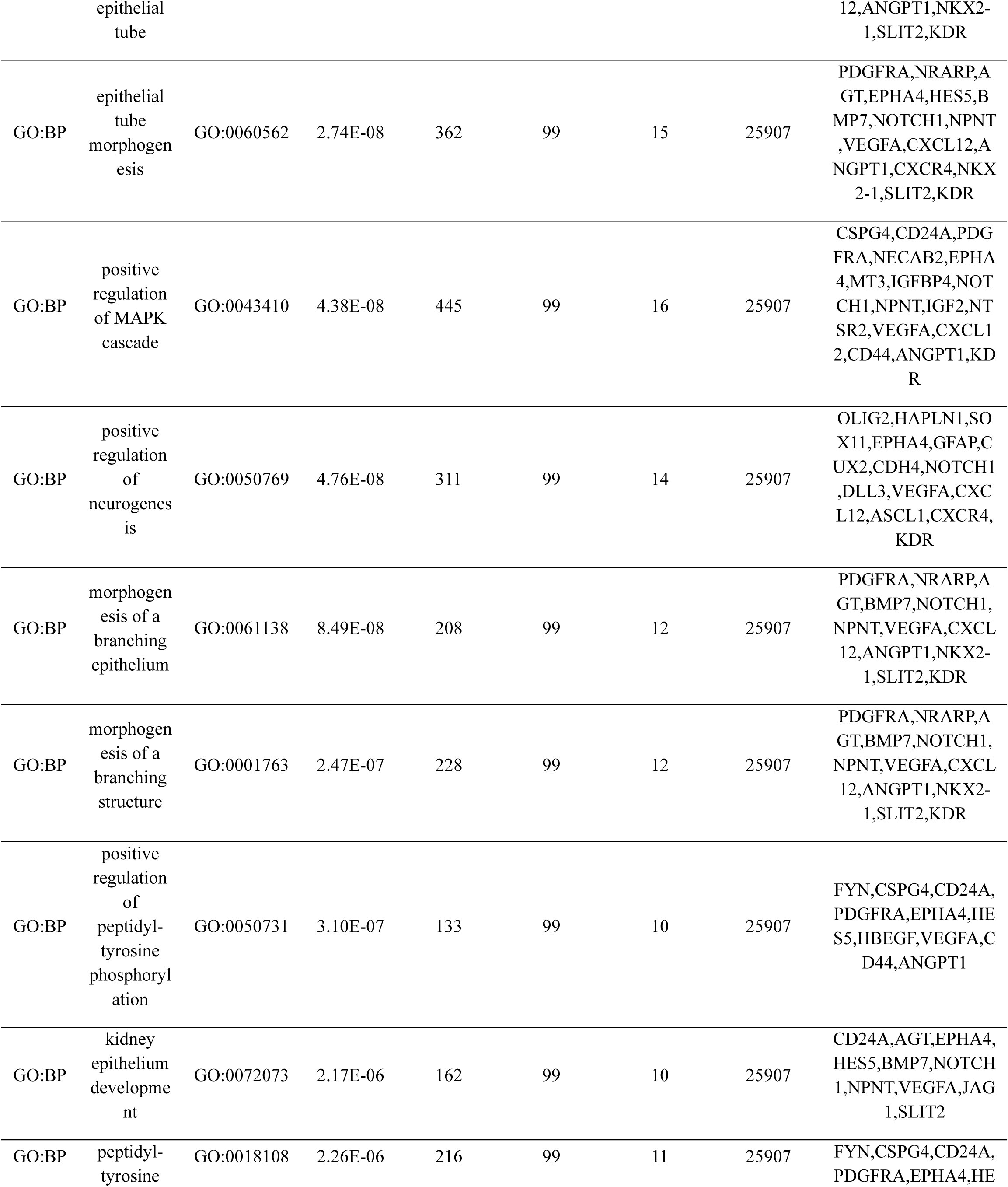
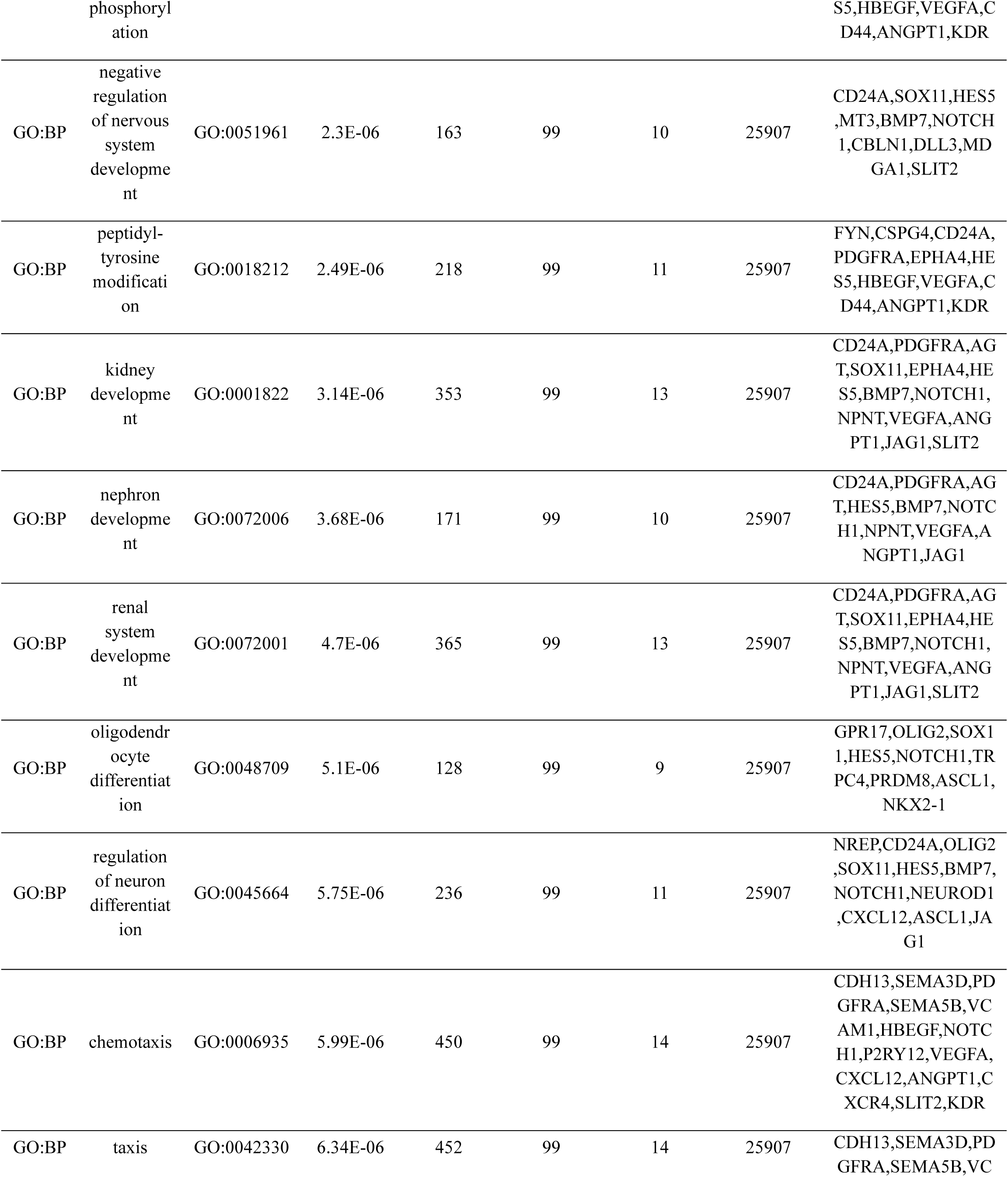
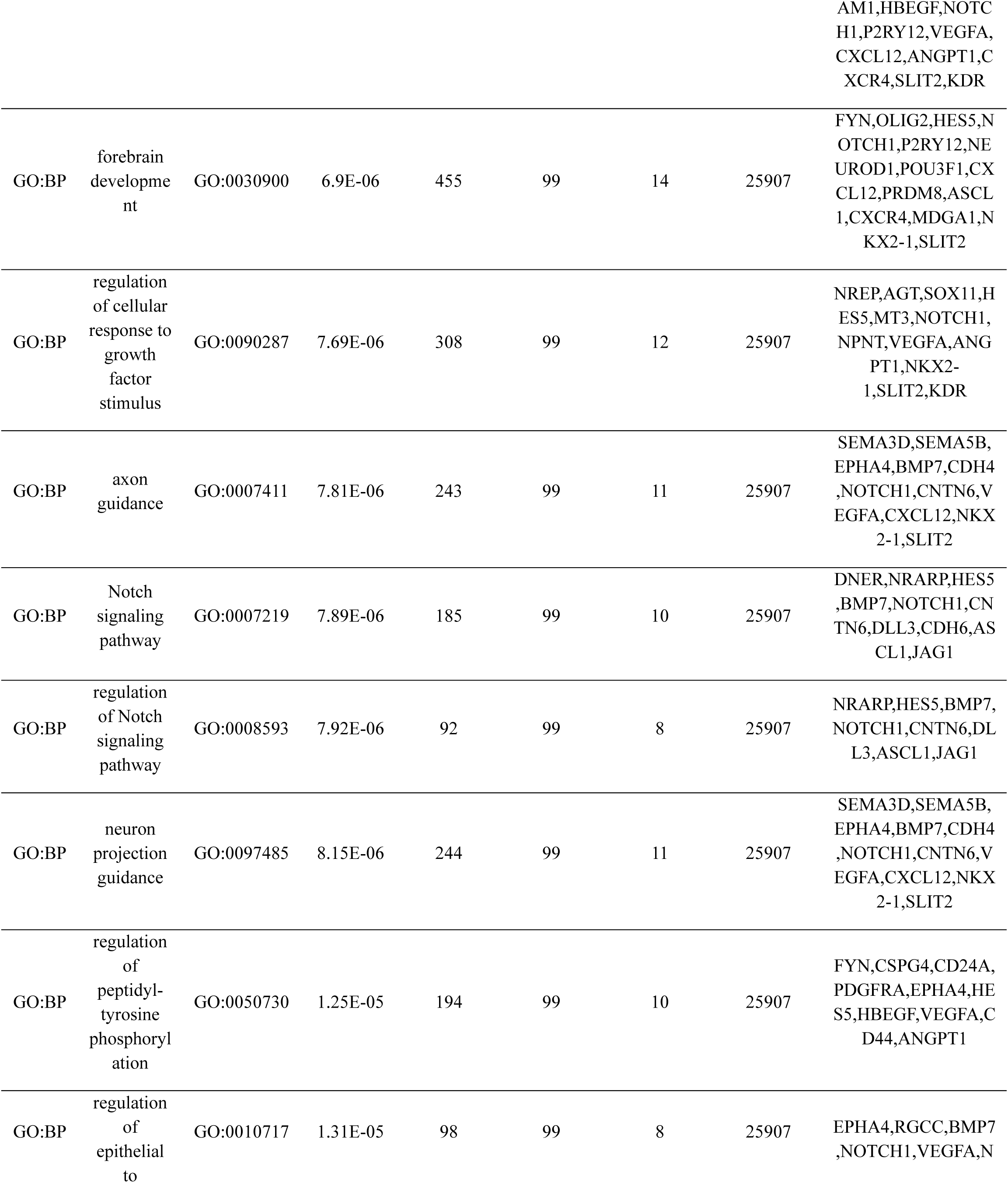
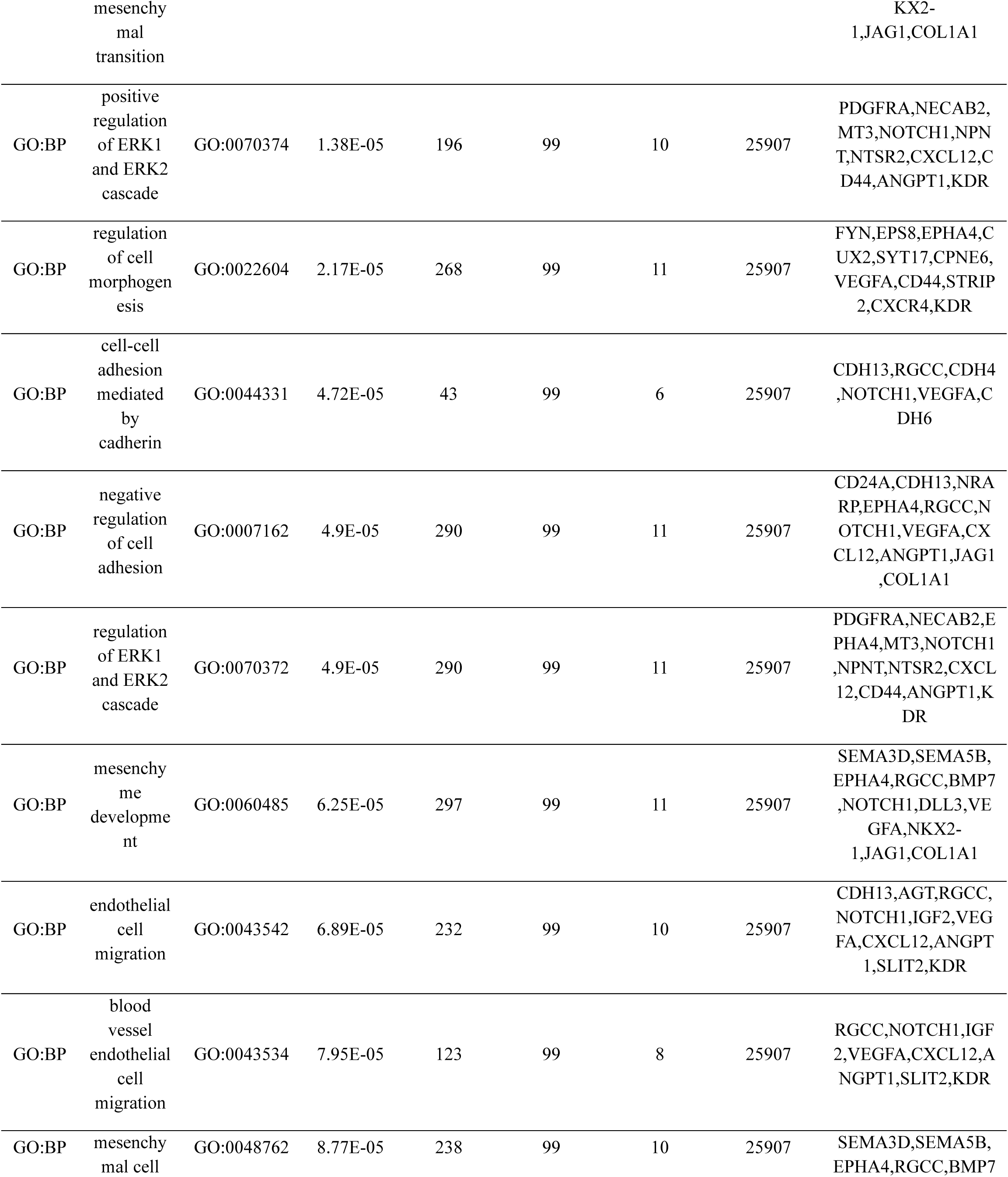
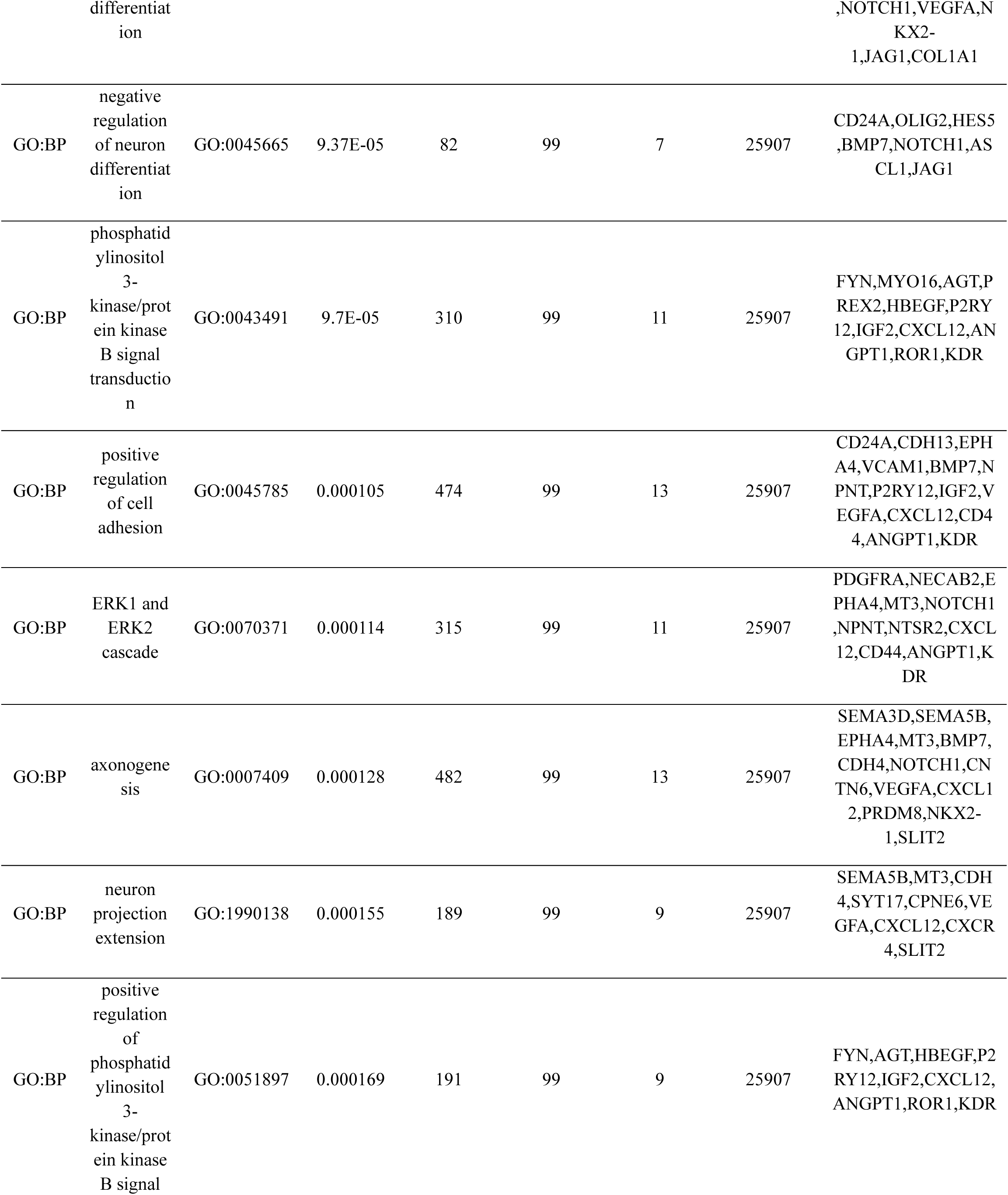
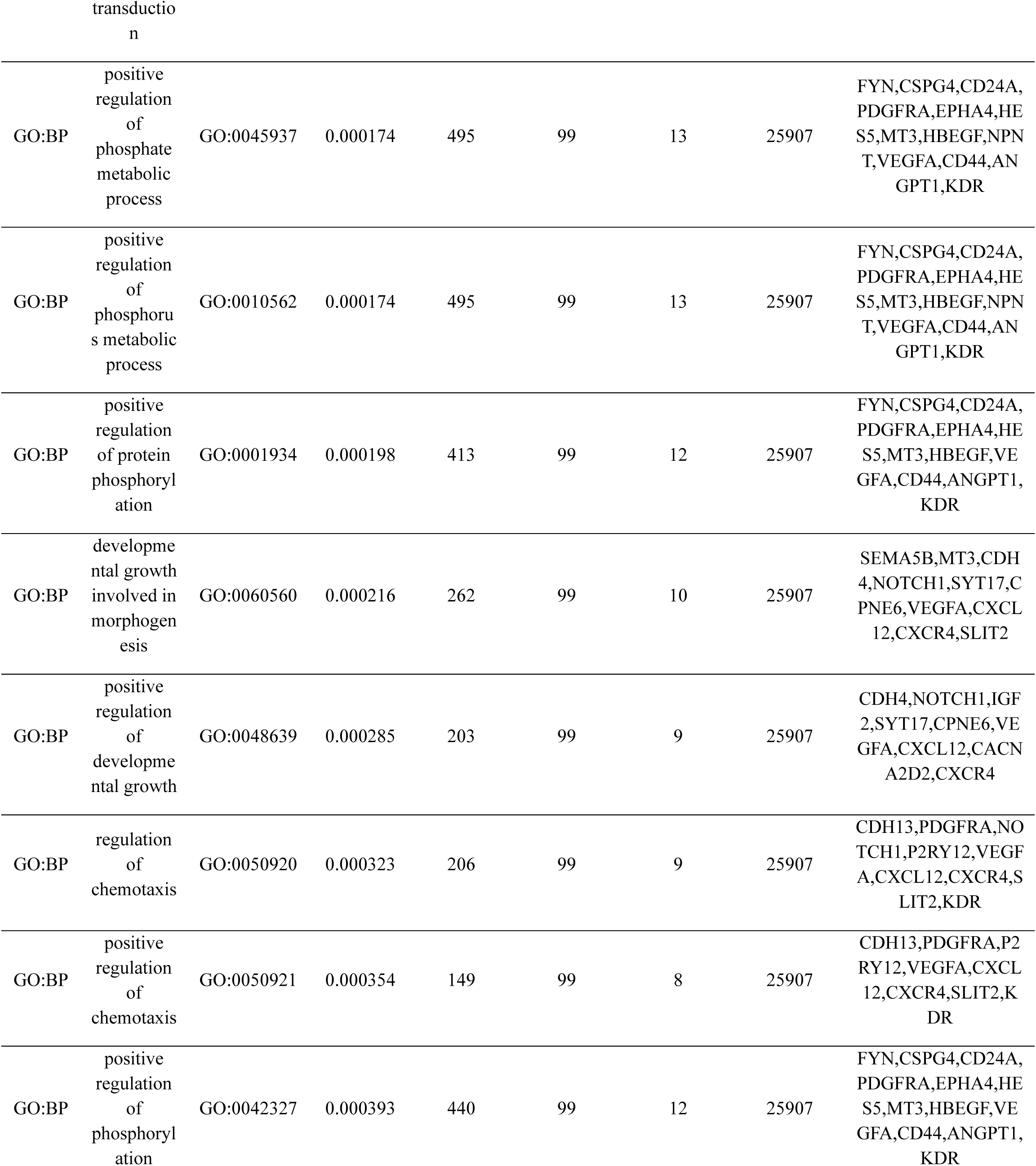
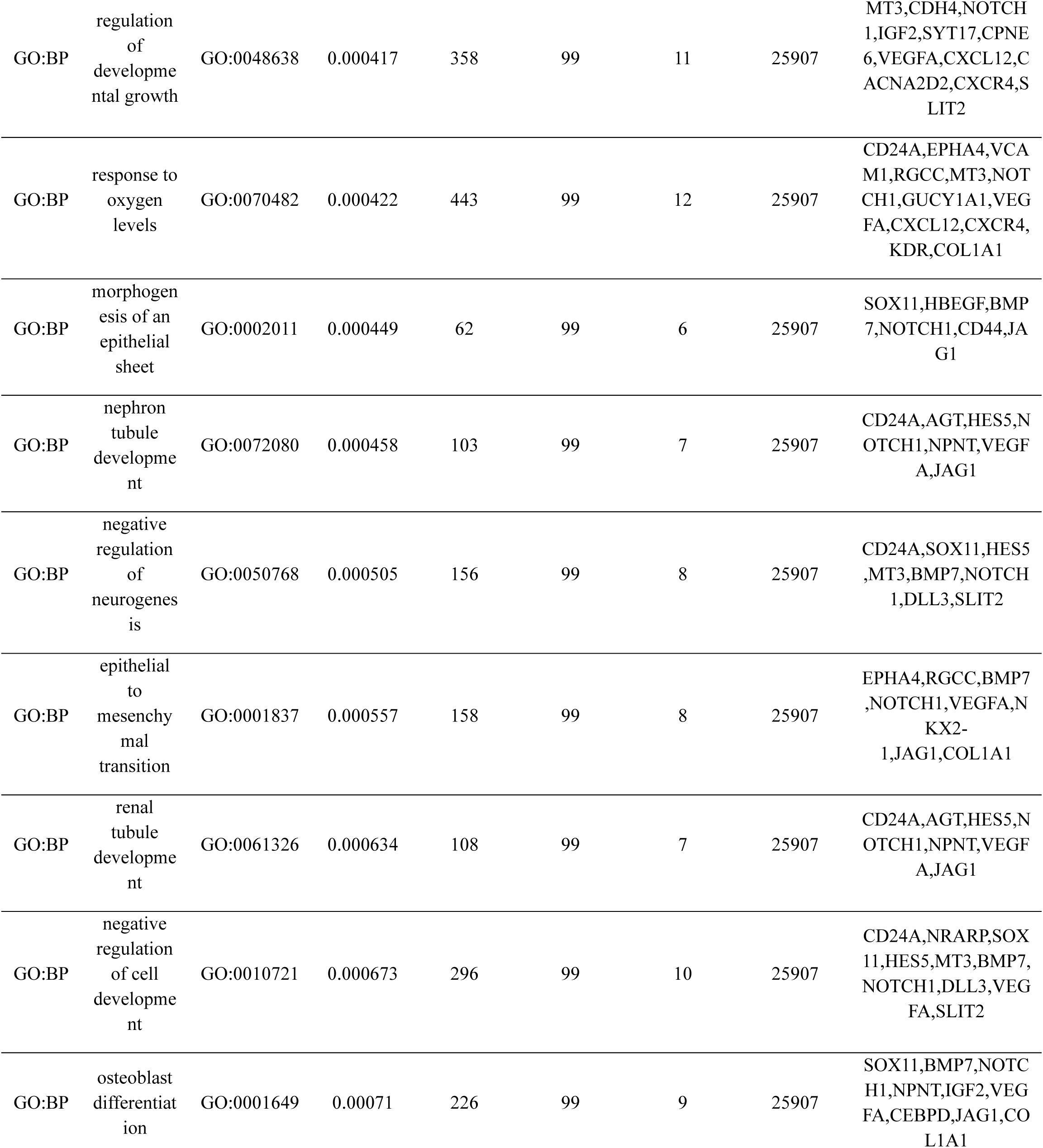
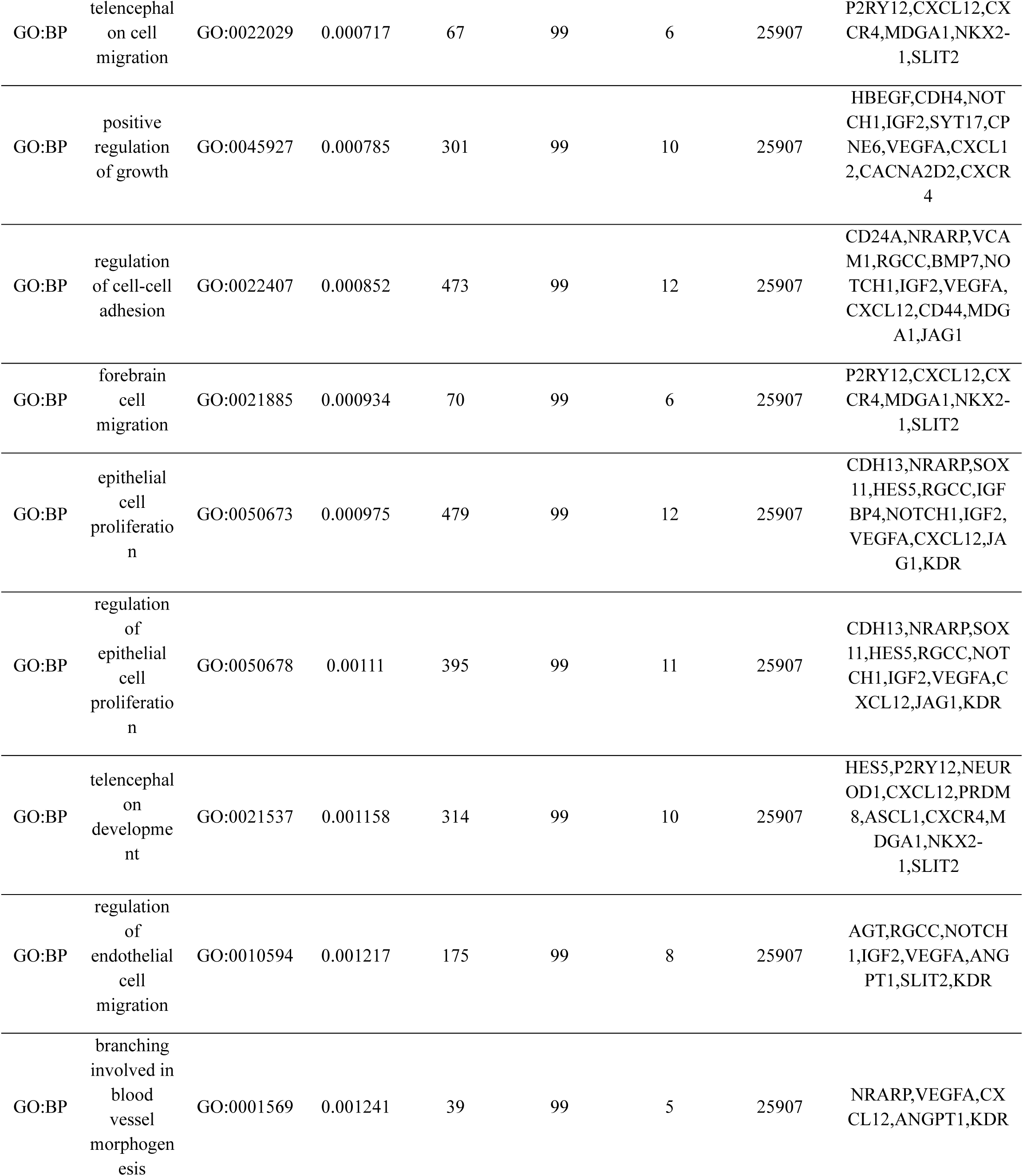
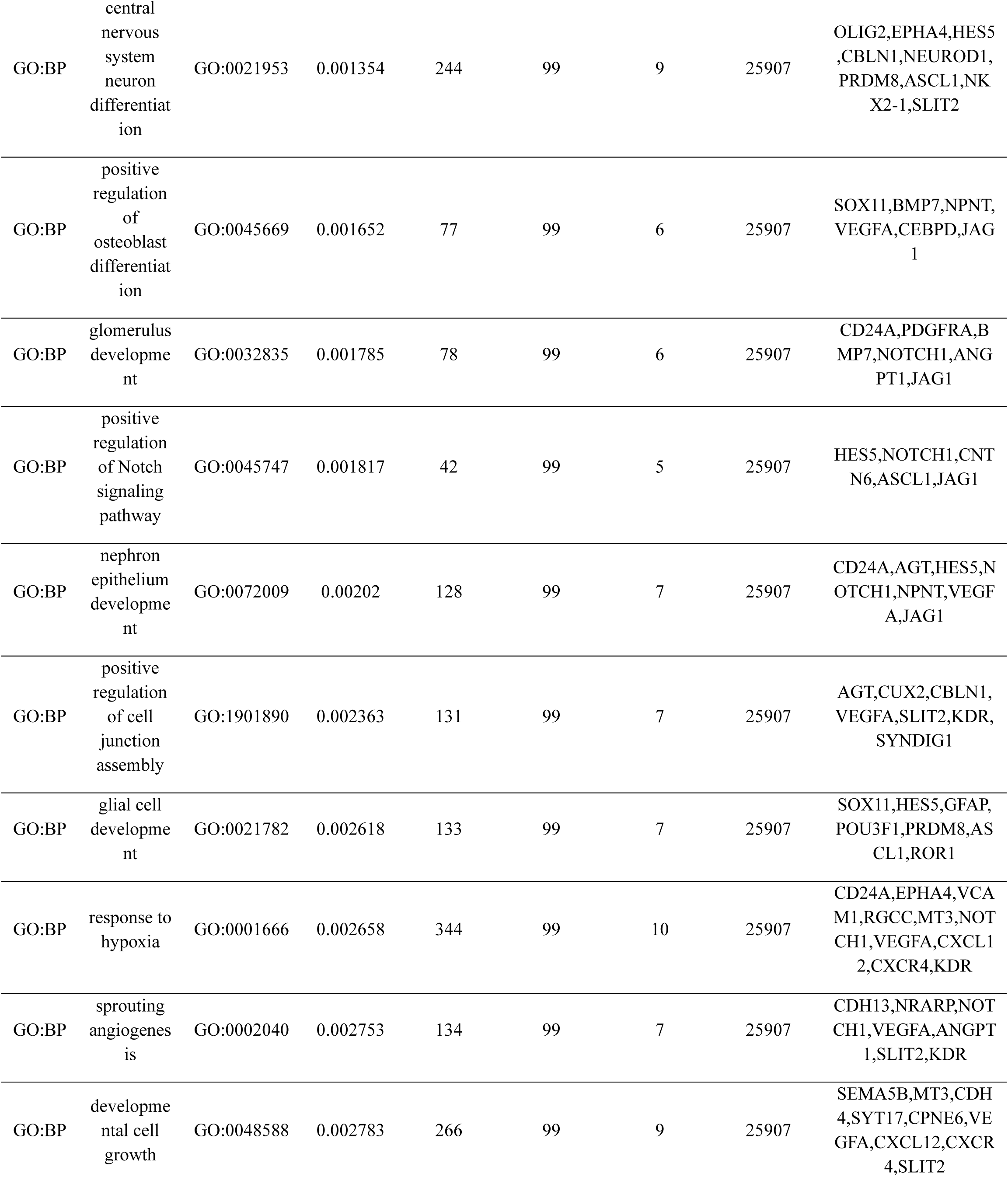
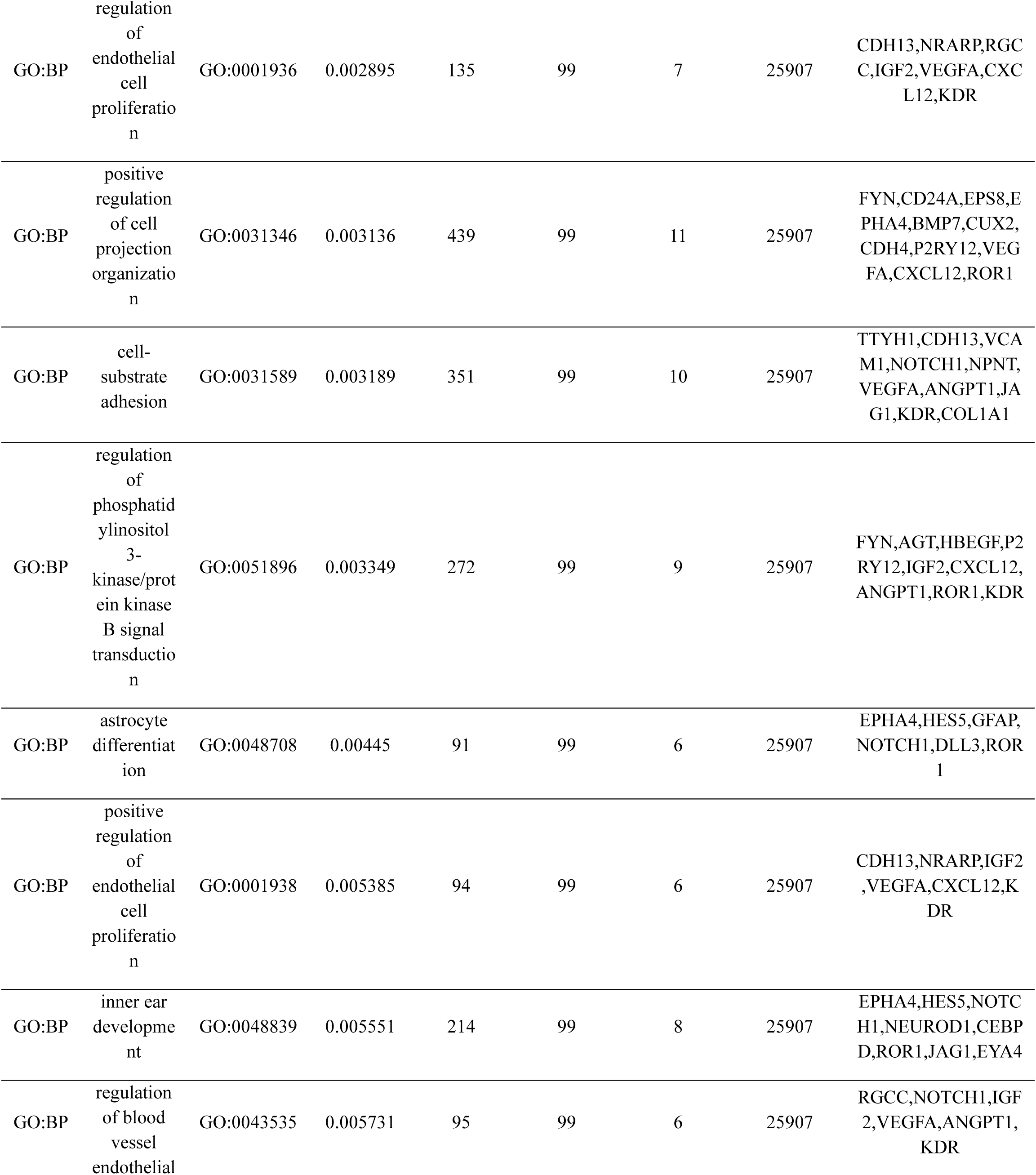
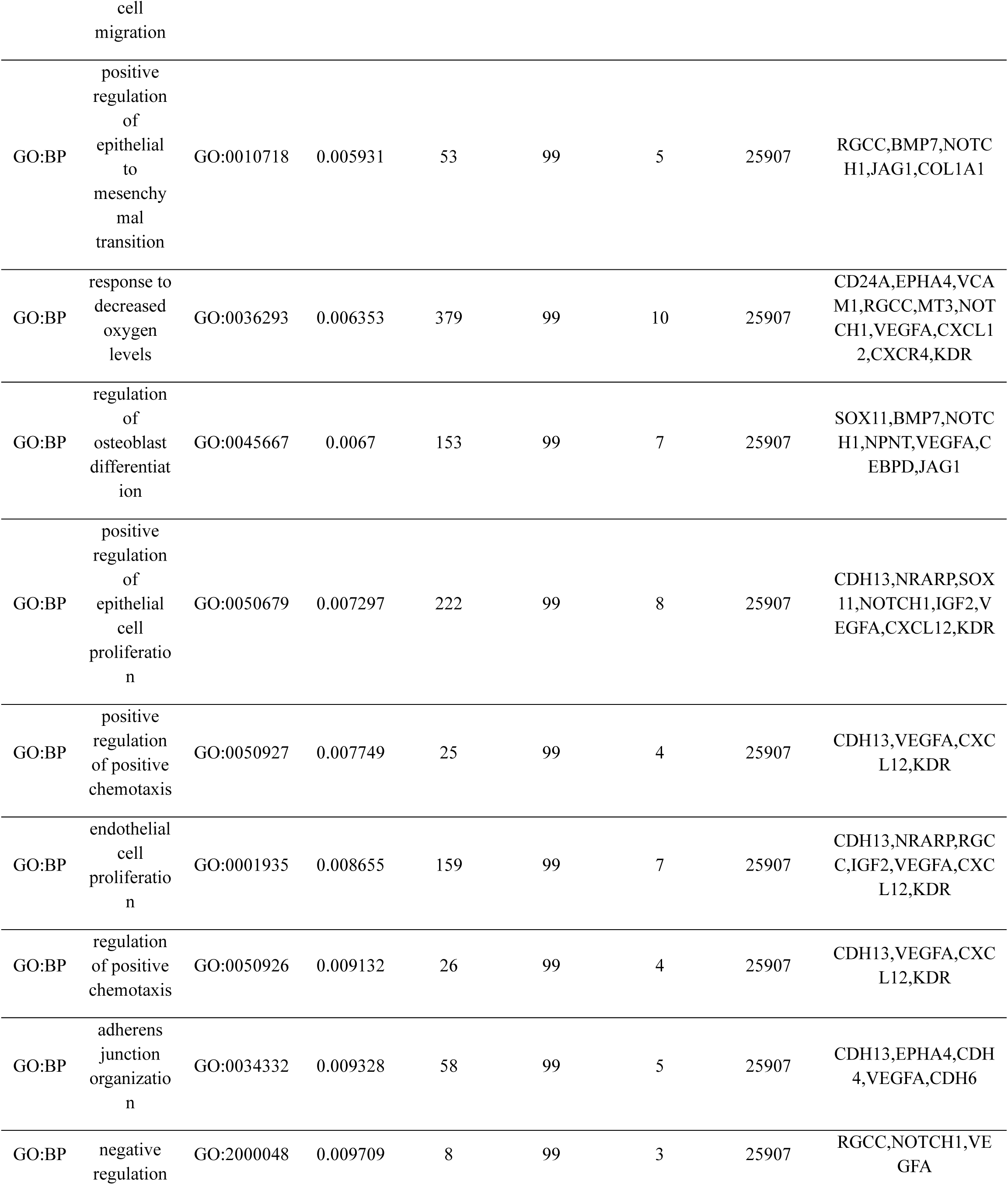
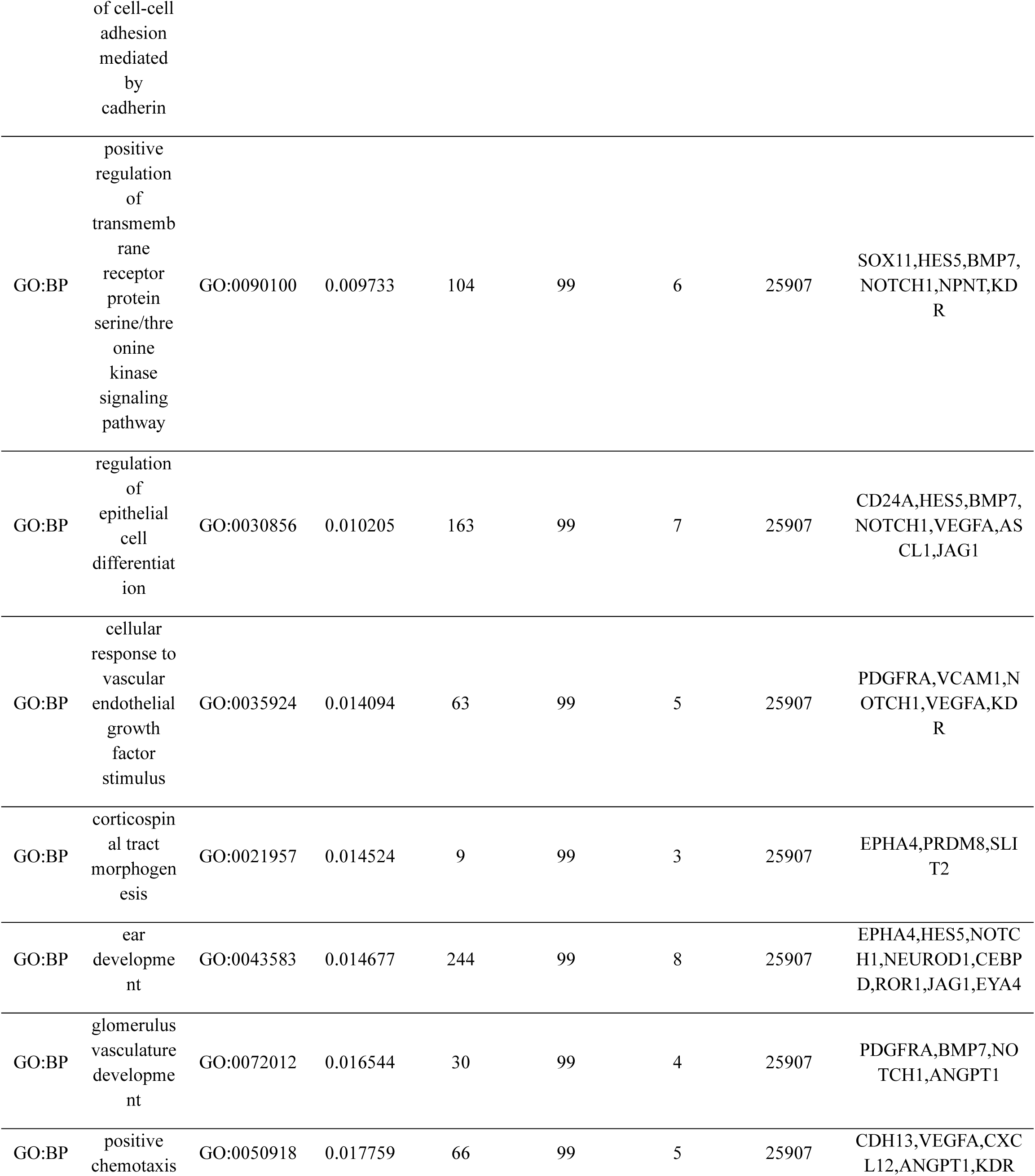
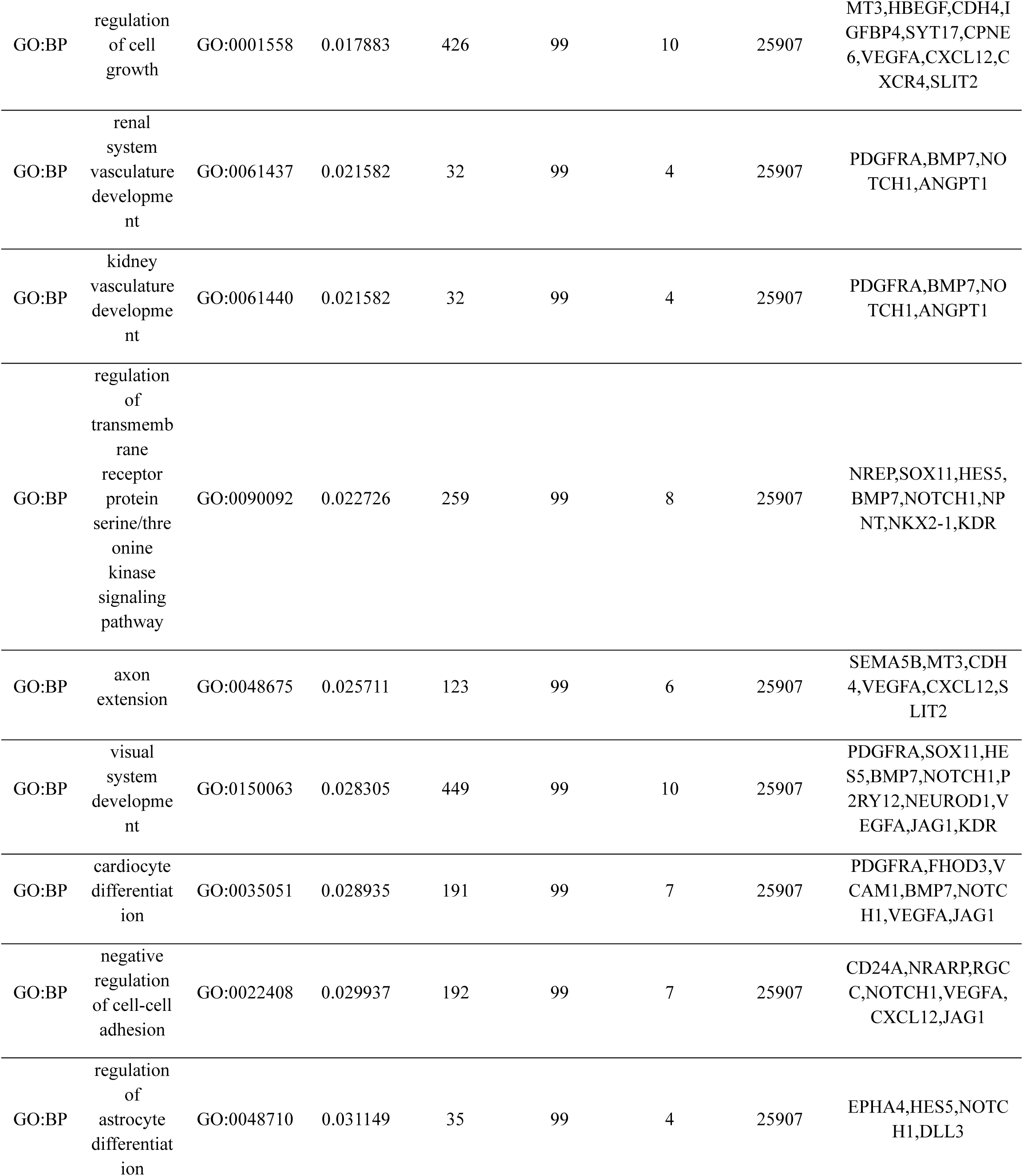
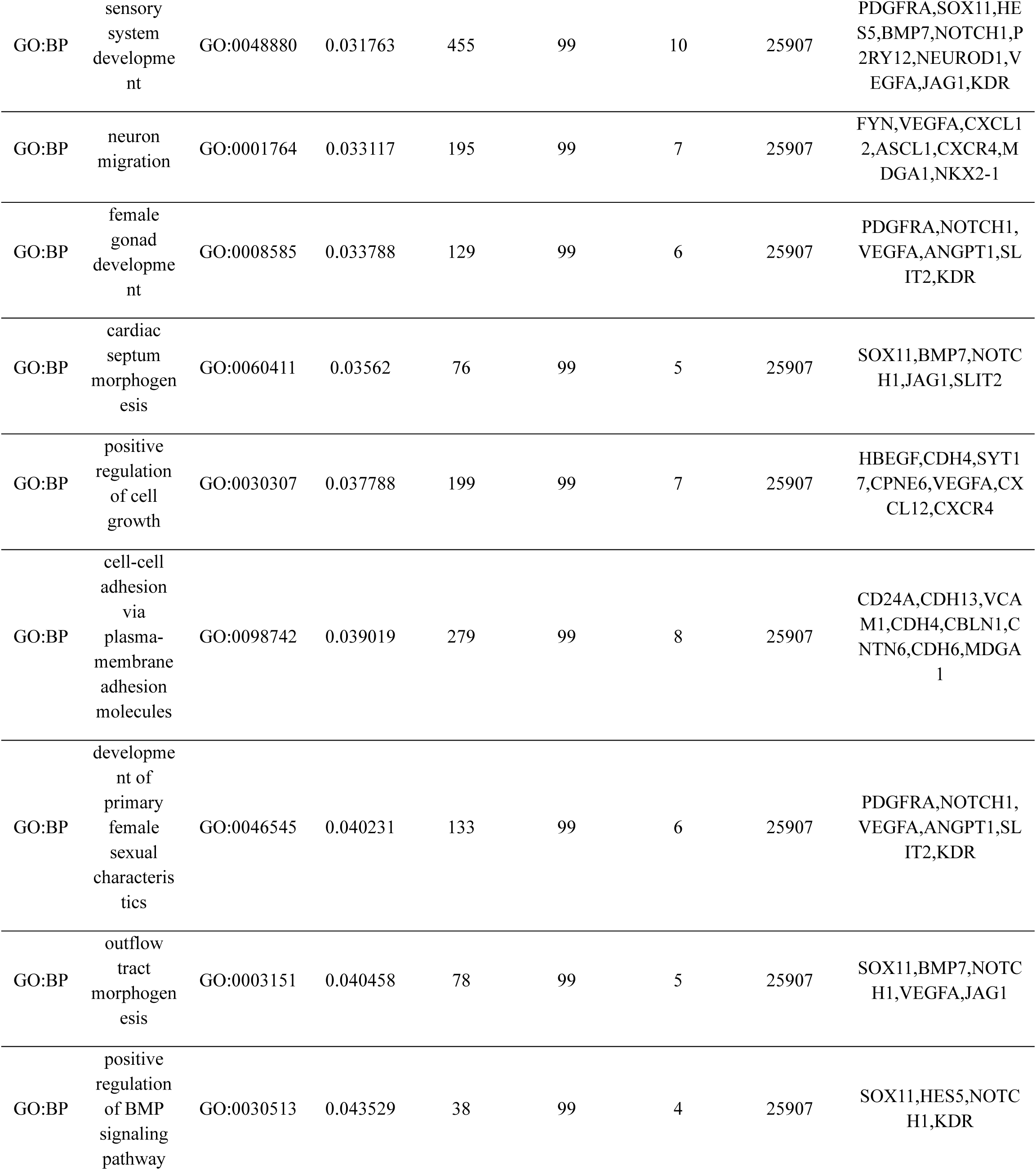
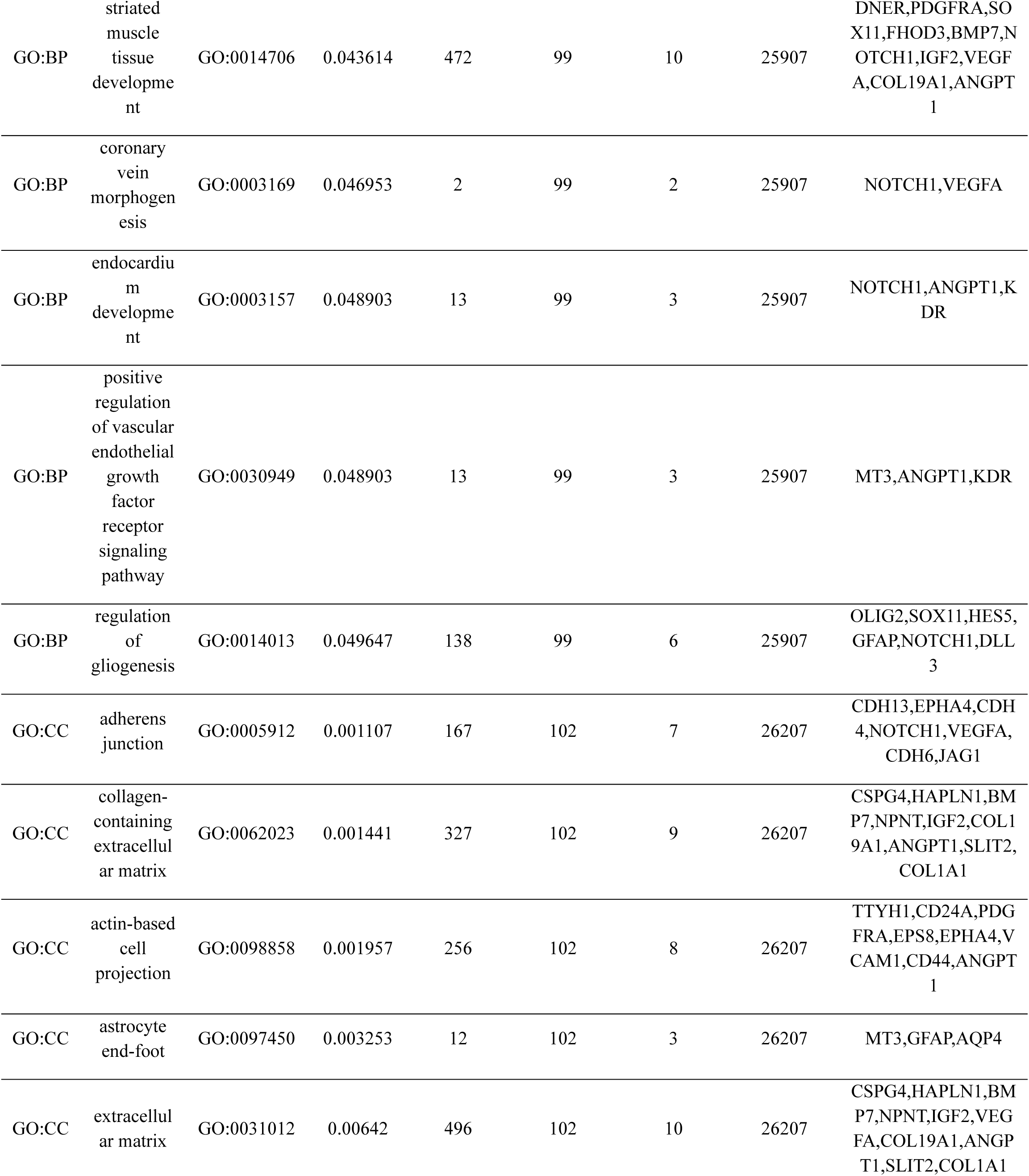
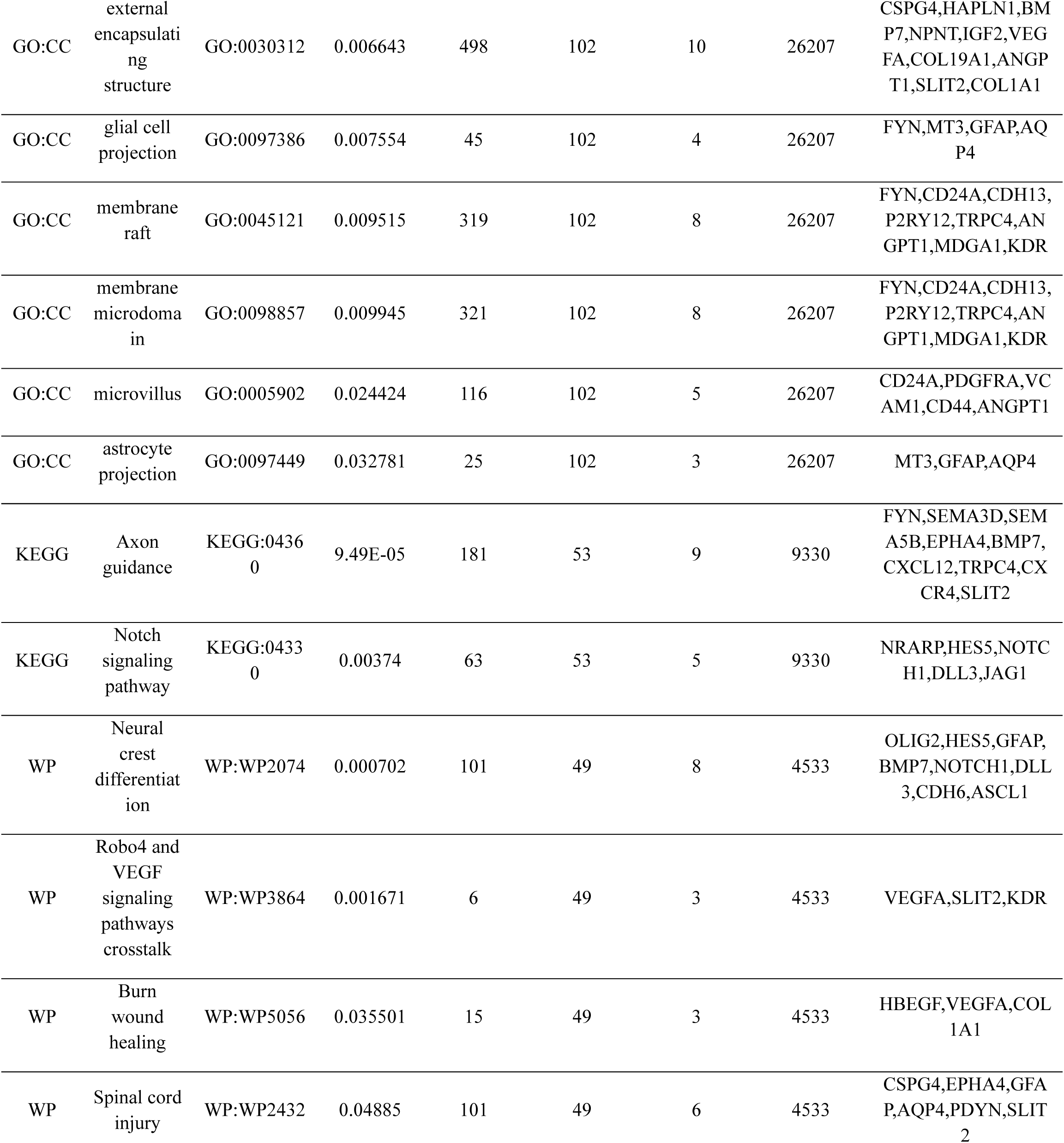
Gene Ontology enrichment analysis on upregulated genes (enriched in younger ages).

**Extended Data Table 8.**
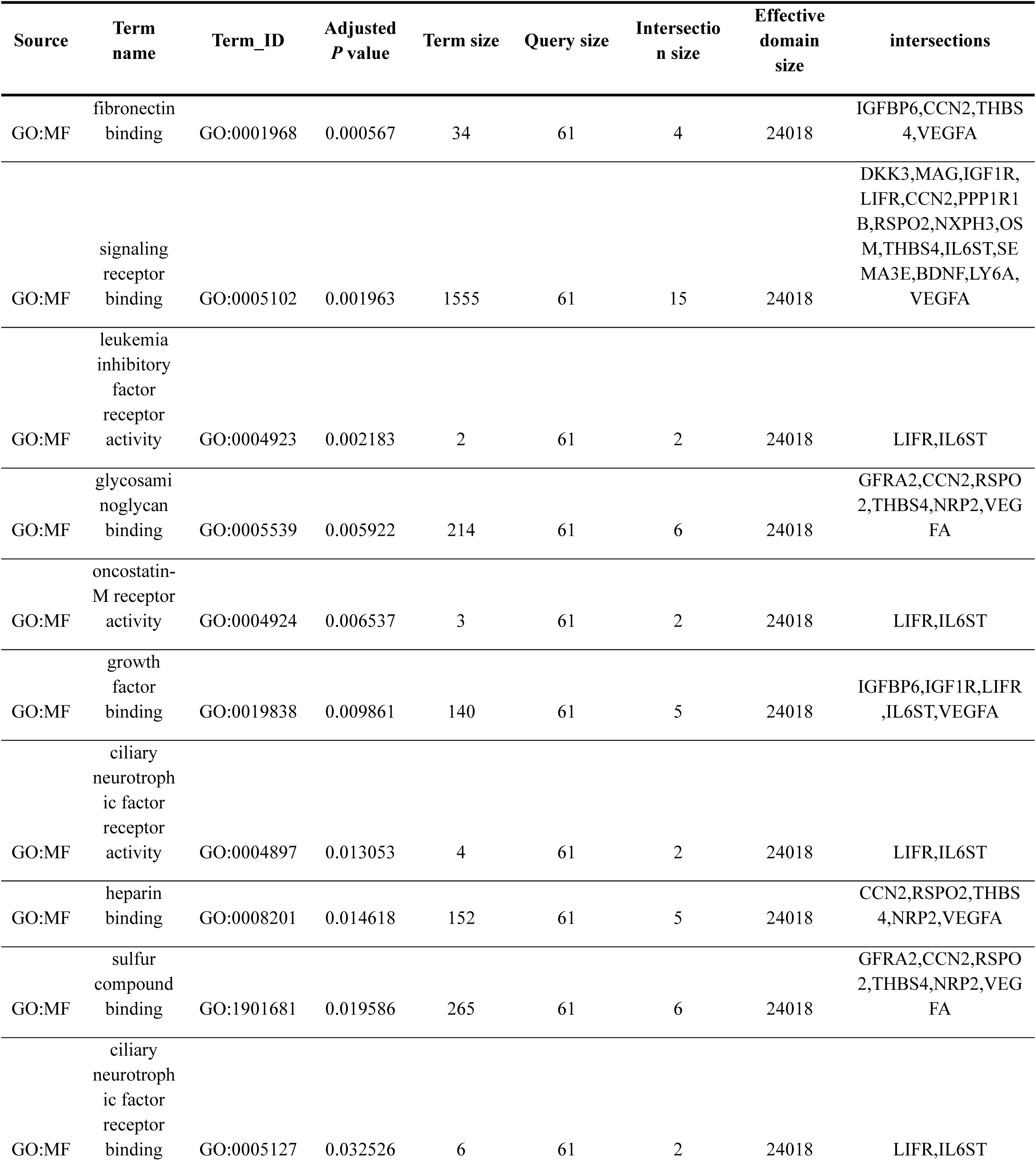
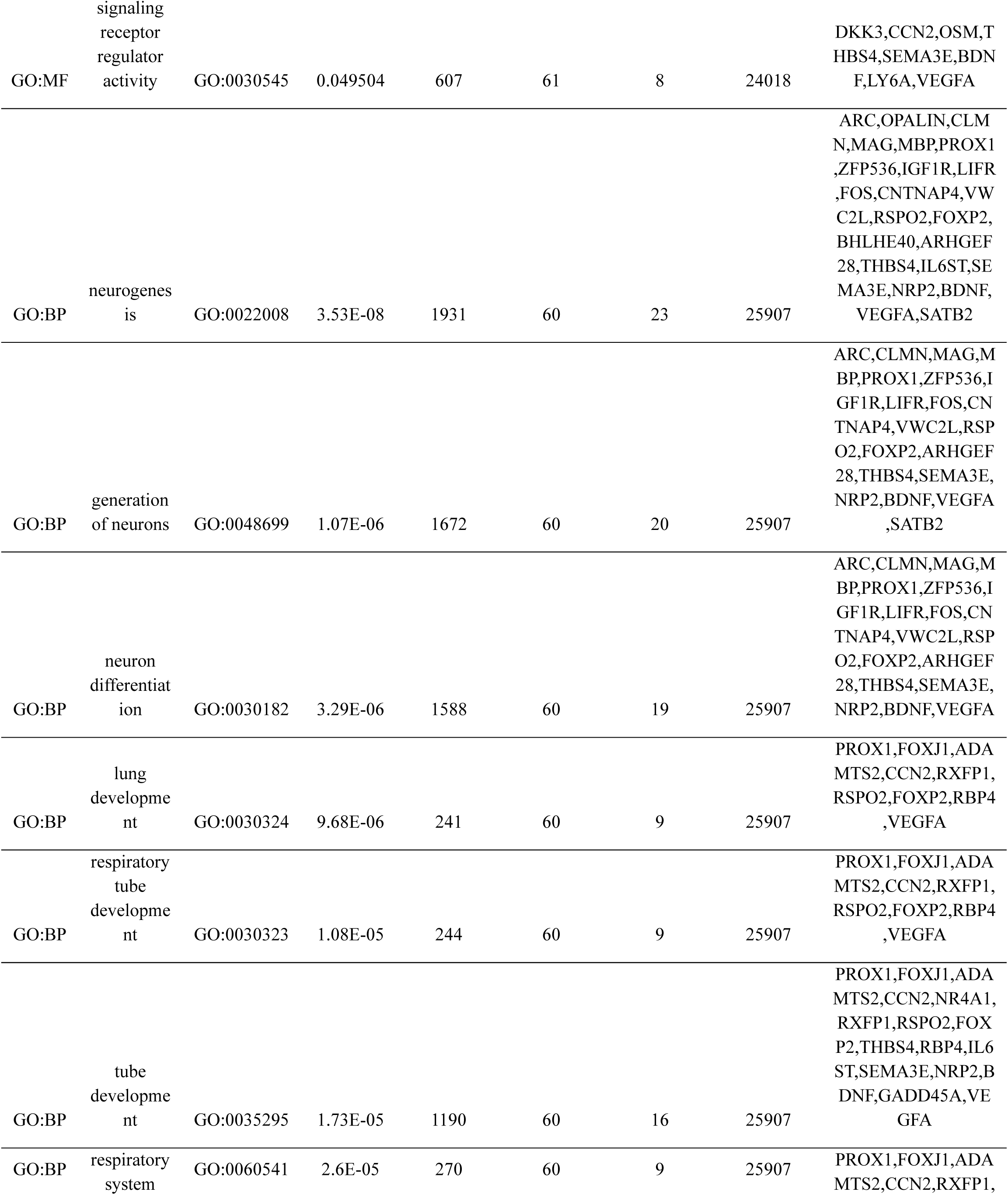
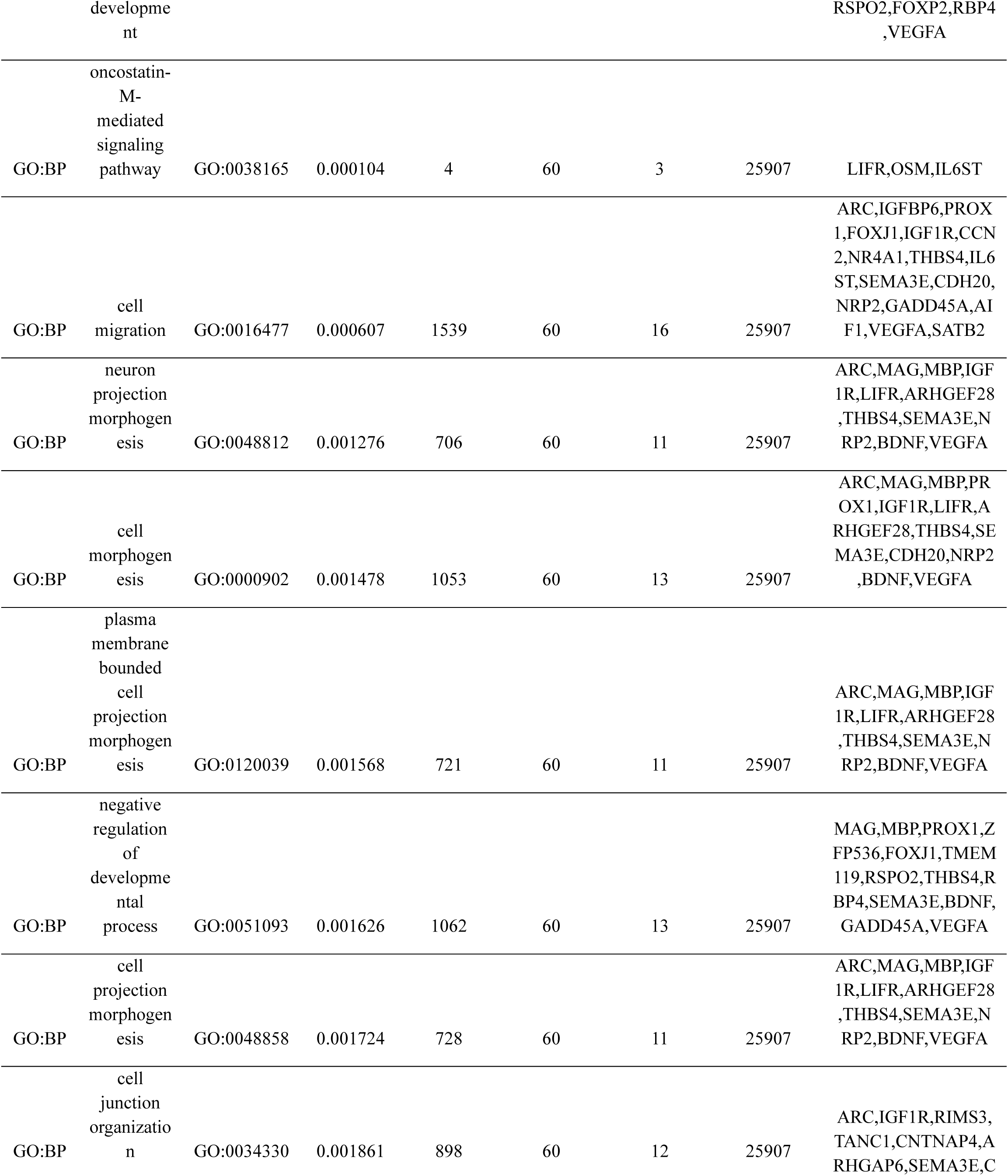
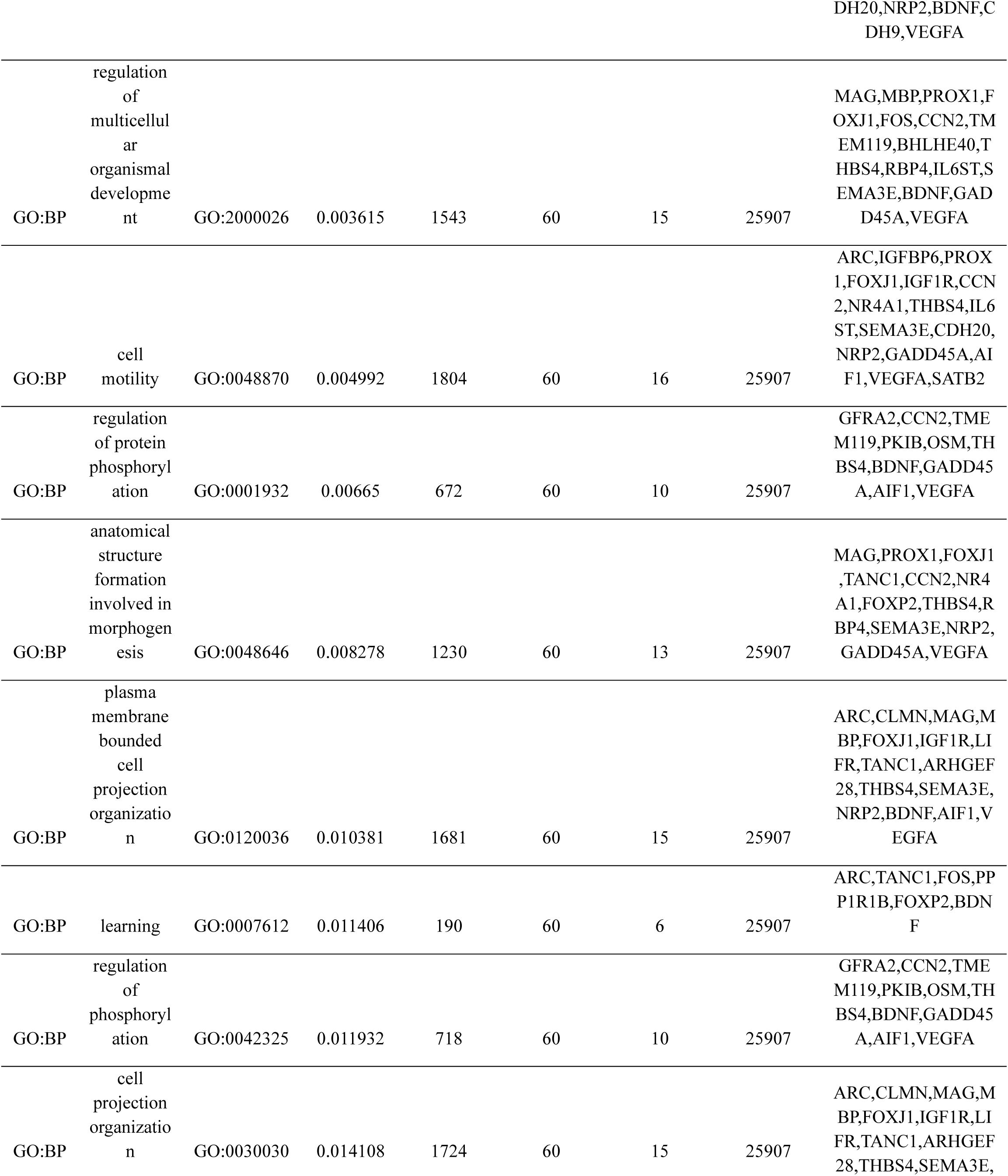
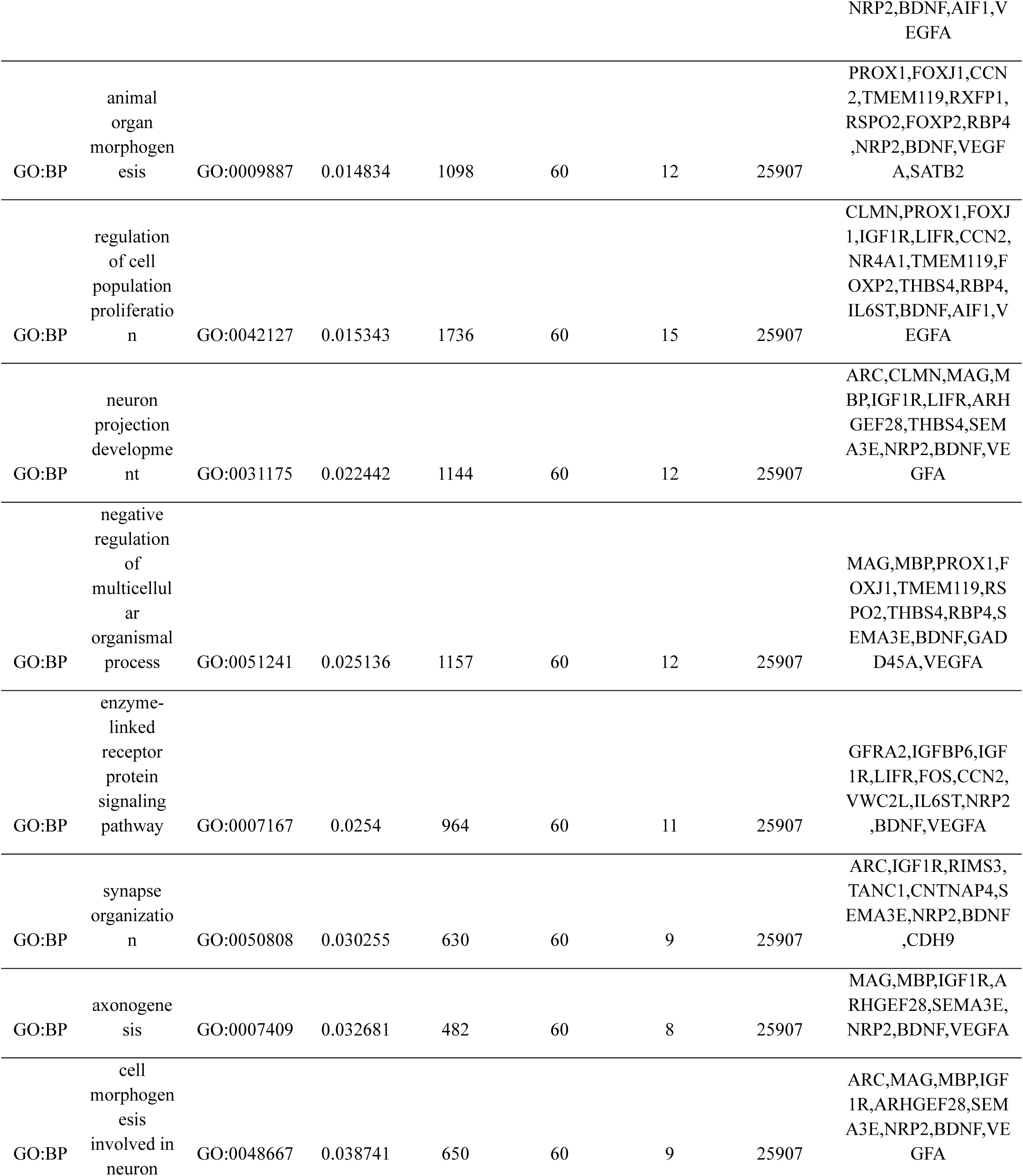
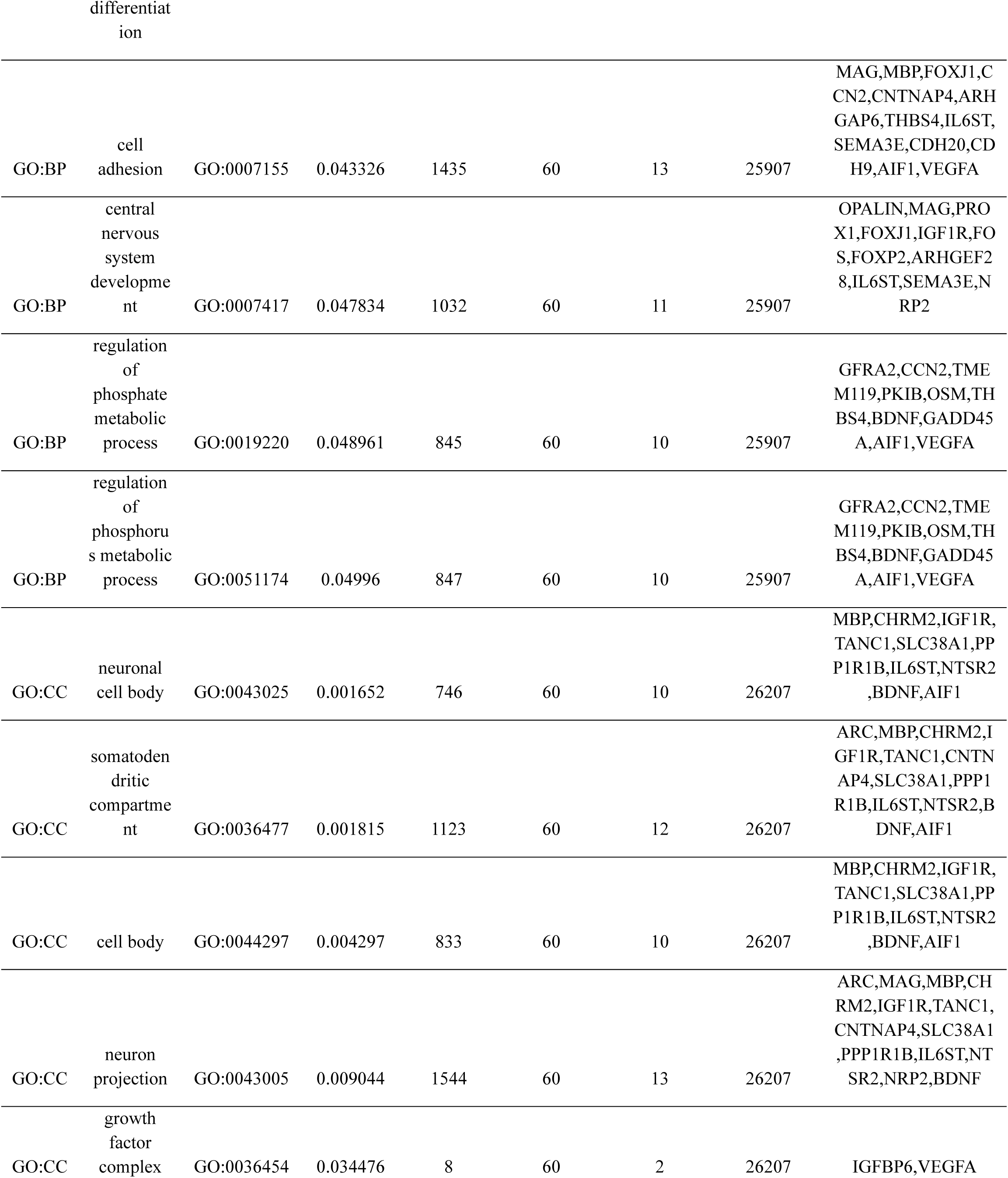
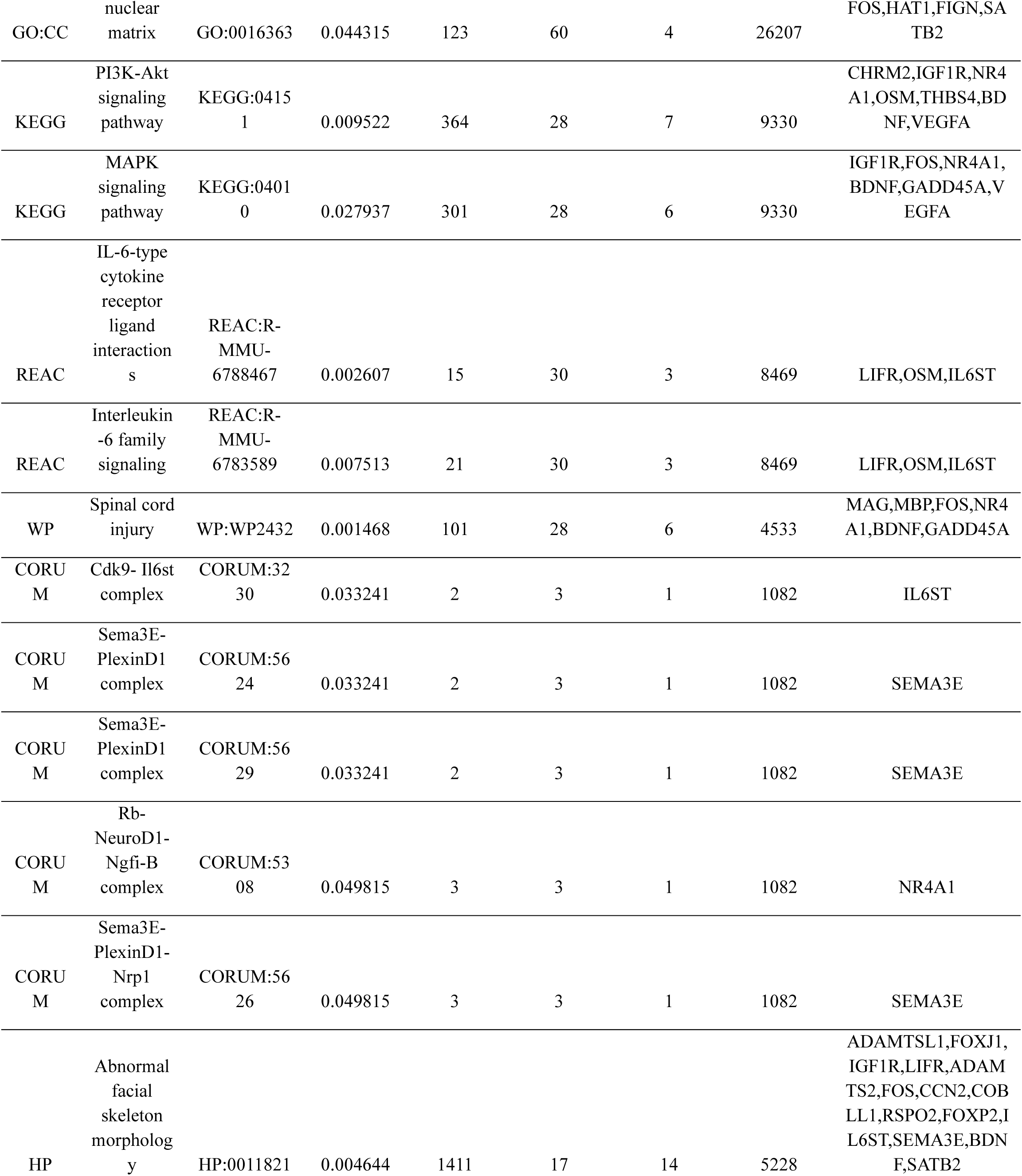
Gene Ontology enrichment analysis on downregulated genes (enriched in older ages).

## Notes

### Competing Interest Statement

The authors have declared no competing interest.

## REFERENCES

1 Tulving, E. Episodic memory: From mind to brain. Annual review of psychology 53, 1–25 (2002).

2 Rubin, D. C. The distribution of early childhood memories. Memory 8, 265–269 (2000).

3 Rubin, D. C. & Schulkind, M. D. The distribution of autobiographical memories across the lifespan. Memory & cognition 25, 859–866 (1997).

4 Bauer, P. J. What do infants recall of their lives? Memory for specific events by one-to two-year-olds. American psychologist 51, 29 (1996).

5 Freud, S. Childhood and concealing memories. (1914).

6 Bauer, P. J. A complementary processes account of the development of childhood amnesia and a personal past. Psychological Review 122, 204 (2015).

7 Wang, Q. Infantile amnesia reconsidered: A cross-cultural analysis. Memory 11, 65–80 (2003).

8 Hayne, H. & Jack, F. Childhood amnesia. Wiley Interdisciplinary Reviews: Cognitive Science 2, 136–145 (2011).

9 Howe, M. L. & Courage, M. L. On resolving the enigma of infantile amnesia. Psychological bulletin 113, 305 (1993).

10 Campbell, B. A. & Campbell, E. H. Retention and extinction of learned fear in infant and adult rats. Journal of comparative and physiological psychology 55, 1 (1962).

11 Travaglia, A., Bisaz, R., Sweet, E. S., Blitzer, R. D. & Alberini, C. M. Infantile amnesia reflects a developmental critical period for hippocampal learning. Nature neuroscience 19, 1225–1233 (2016).

12 Akers, K. G. et al. Hippocampal neurogenesis regulates forgetting during adulthood and infancy. Science 344, 598–602 (2014).

13 Power, S. D. et al. Immune activation state modulates infant engram expression across development. Science Advances 9, eadg9921 (2023).

14 Alberini, C. M. & Travaglia, A. Infantile amnesia: a critical period of learning to learn and remember. Journal of Neuroscience 37, 5783–5795 (2017).

15 Campbell, B. A. & Spear, N. E. Ontogeny of memory. Psychological review 79, 215 (1972).

16 Josselyn, S. A. & Frankland, P. W. Infantile amnesia: a neurogenic hypothesis. Learning & Memory 19, 423–433 (2012).

17 Ramsaran, A. I., Schlichting, M. L. & Frankland, P. W. The ontogeny of memory persistence and specificity. Developmental Cognitive Neuroscience 36, 100591 (2019).

18 Bevandić, J. et al. Episodic memory development: Bridging animal and human research. Neuron 112, 1060–1080 (2024).

19 Bauer, P. J., Wenner, J. A., Dropik, P. L., Wewerka, S. S. & Howe, M. L. Parameters of remembering and forgetting in the transition from infancy to early childhood. Monographs of the Society for Research in Child Development, i-213 (2000).

20 Keresztes, A., Ngo, C. T., Lindenberger, U., Werkle-Bergner, M. & Newcombe, N. S. Hippocampal maturation drives memory from generalization to specificity. Trends in Cognitive Sciences 22, 676–686 (2018).

21 Yates, T. S. et al. Hippocampal encoding of memories in human infants. Science 387, 1316–1320 (2025).

22 Akers, K. G., Arruda-Carvalho, M., Josselyn, S. A. & Frankland, P. W. Ontogeny of contextual fear memory formation, specificity, and persistence in mice. Learning & memory 19, 598–604 (2012).

23 Klune, C. B., Jin, B. & DeNardo, L. A. Linking mPFC circuit maturation to the developmental regulation of emotional memory and cognitive flexibility. Elife 10, e64567 (2021).

24 Frankland, P. W. & Bontempi, B. The organization of recent and remote memories. Nature reviews neuroscience 6, 119–130 (2005).

25 Gilboa, A. & Moscovitch, M. No consolidation without representation: Correspondence between neural and psychological representations in recent and remote memory. Neuron 109, 2239–2255 (2021).

26 Allen, T. A. & Fortin, N. J. The evolution of episodic memory. Proceedings of the National Academy of Sciences 110, 10379–10386 (2013).

27 Fanselow, M. S. Conditional and unconditional components of post-shock freezing. The Pavlovian journal of biological science: Official Journal of the Pavlovian 15, 177–182 (1980).

28 Bonnici, H. M. et al. Detecting representations of recent and remote autobiographical memories in vmPFC and hippocampus. Journal of Neuroscience 32, 16982–16991 (2012).

29 DeNardo, L. A. et al. Temporal evolution of cortical ensembles promoting remote memory retrieval. Nature neuroscience 22, 460–469 (2019).

30 Golbabaei, A., Josselyn, S. A. & Frankland, P. W. PV-dependent reorganization of prelimbic cortex sub-engrams during systems consolidation. Neuron (2025).

31 Kitamura, T. et al. Engrams and circuits crucial for systems consolidation of a memory. Science 356, 73–78 (2017).

32 McCormick, C., Barry, D. N., Jafarian, A., Barnes, G. R. & Maguire, E. A. vmPFC drives hippocampal processing during autobiographical memory recall regardless of remoteness. Cerebral Cortex 30, 5972–5987 (2020).

33 Roth, B. L. DREADDs for neuroscientists. Neuron 89, 683–694 (2016).

34 Chen, M. B., Jiang, X., Quake, S. R. & Südhof, T. C. Persistent transcriptional programmes are associated with remote memory. Nature 587, 437–442 (2020).

35 Bergles, D. E. & Richardson, W. D. Oligodendrocyte development and plasticity. Cold Spring Harbor perspectives in biology 8, a020453 (2016).

36 Hilscher, M. M. et al. Spatial and temporal heterogeneity in the lineage progression of fine oligodendrocyte subtypes. BMC biology 20, 122 (2022).

37 Xiao, J. et al. Brain-derived neurotrophic factor promotes central nervous system myelination via a direct effect upon oligodendrocytes. Neurosignals 18, 186–202 (2011).

38 Vondran, M. W., Clinton-Luke, P., Honeywell, J. Z. & Dreyfus, C. F. BDNF+/− mice exhibit deficits in oligodendrocyte lineage cells of the basal forebrain. Glia 58, 848–856 (2010).

39 Wang, S. et al. Notch receptor activation inhibits oligodendrocyte differentiation. Neuron 21, 63–75 (1998).

40 Wong, A. W., Xiao, J., Kemper, D., Kilpatrick, T. J. & Murray, S. S. Oligodendroglial expression of TrkB independently regulates myelination and progenitor cell proliferation. Journal of Neuroscience 33, 4947–4957 (2013).

41 Fletcher, J. L. et al. Targeting TrkB with a brain-derived neurotrophic factor mimetic promotes myelin repair in the brain. Journal of Neuroscience 38, 7088–7099 (2018).

42 Monje, M. Myelin plasticity and nervous system function. Annual review of neuroscience 41, 61–76 (2018).

43 Jia, M., Travaglia, A., Pollonini, G., Fedele, G. & Alberini, C. M. Developmental changes in plasticity, synaptic, glia, and connectivity protein levels in rat medial prefrontal cortex. Learning & Memory 25, 533–543 (2018).

44 Nagappan, G. et al. Control of extracellular cleavage of ProBDNF by high frequency neuronal activity. Proceedings of the National Academy of Sciences 106, 1267–1272 (2009).

45 Barde, Y.-A. The physiopathology of brain-derived neurotrophic factor. Physiological Reviews (2025).

46 Korrell, K. V. et al. Differential effect on myelination through abolition of activity-dependent synaptic vesicle release or reduction of overall electrical activity of selected cortical projections in the mouse. Journal of Anatomy 235, 452–467 (2019).

47. Alshehri, B., Pagnin, M., Lee, J., Petratos, S. & Richardson, S. (2020).

48 Aranmolate, A., Tse, N. & Colognato, H. Myelination is delayed during postnatal brain development in the mdx mouse model of Duchenne muscular dystrophy. BMC neuroscience 18, 63 (2017).

49 Lang, J. et al. Adenomatous polyposis coli regulates oligodendroglial development. Journal of Neuroscience 33, 3113–3130 (2013).

50 Fletcher, J. L., Makowiecki, K., Cullen, C. L. & Young, K. M. in Seminars in cell & developmental biology. 14–23 (Elsevier).

51 Hughes, E. G., Orthmann-Murphy, J. L., Langseth, A. J. & Bergles, D. E. Myelin remodeling through experience-dependent oligodendrogenesis in the adult somatosensory cortex. Nature neuroscience 21, 696–706 (2018).

52 Donato, F., Jacobsen, R. I., Moser, M.-B. & Moser, E. I. Stellate cells drive maturation of the entorhinal-hippocampal circuit. Science 355, eaai8178 (2017).

53 Pang, P. T. et al. Cleavage of proBDNF by tPA/plasmin is essential for long-term hippocampal plasticity. Science 306, 487–491 (2004).

54 Todd, D. et al. A monoclonal antibody TrkB receptor agonist as a potential therapeutic for Huntington’s disease. PloS one **G**, e87923 (2014).

55 Hensch, T. K. Critical period plasticity in local cortical circuits. Nature reviews neuroscience 6, 877–888 (2005).

56 Xin, W. et al. Oligodendrocytes and myelin limit neuronal plasticity in visual cortex. Nature 633, 856–863 (2024).

57 Makinodan, M., Rosen, K. M., Ito, S. & Corfas, G. A critical period for social experience–dependent oligodendrocyte maturation and myelination. science 337, 1357–1360 (2012).

58 McGee, A. W., Yang, Y., Fischer, Q. S., Daw, N. W. & Strittmatter, S. M. Experience-driven plasticity of visual cortex limited by myelin and Nogo receptor. Science 309, 2222–2226 (2005).

59 Talidou, A., Frankland, P. W., Mabbott, D. & Lefebvre, J. Homeostatic coordination and up-regulation of neural activity by activity-dependent myelination. Nature Computational Science 2, 665–676 (2022).

60 Atwal, J. K. et al. PirB is a functional receptor for myelin inhibitors of axonal regeneration. Science 322, 967–970 (2008).

61 Cafferty, W. B., Duffy, P., Huebner, E. & Strittmatter, S. M. MAG and OMgp synergize with Nogo-A to restrict axonal growth and neurological recovery after spinal cord trauma. Journal of Neuroscience 30, 6825–6837 (2010).

62 Syken, J., GrandPre, T., Kanold, P. O. & Shatz, C. J. PirB restricts ocular-dominance plasticity in visual cortex. science 313, 1795–1800 (2006).

63 Guskjolen, A. et al. Recovery of “lost” infant memories in mice. Current Biology 28, 2283–2290. e2283 (2018).

64 Lahr, M., Imhof, F., Mauro, L., Ulmer, T. & Donato, F. The Reinstatement of a Forgotten Infantile Memory. bioRxiv, 2025.2009. 2027.678956 (2025).

65 Ryan, T. J. & Frankland, P. W. Forgetting as a form of adaptive engram cell plasticity. Nature Reviews Neuroscience 23, 173–186 (2022).

66 Emery, B. et al. Myelin gene regulatory factor is a critical transcriptional regulator required for CNS myelination. Cell 138, 172–185 (2009).

67 Steadman, P. E. et al. Disruption of oligodendrogenesis impairs memory consolidation in adult mice. Neuron 105, 150–164. e156 (2020).

68 Ramsaran, A. I. et al. A shift in the mechanisms controlling hippocampal engram formation during brain maturation. Science 380, 543–551 (2023).

69 Willis, A., et al. Single cell approaches define neural stem cell niches and identify microglial ligands that can enhance precursor-mediated oligodendrogenesis. Cell Reports 44 (2025).

70 Janesick, A. et al. High resolution mapping of the tumor microenvironment using integrated single-cell, spatial and in situ analysis. Nature communications 14, 8353 (2023).

71 Kolberg, L. et al. g: Profiler—interoperable web service for functional enrichment analysis and gene identifier mapping (2023 update). Nucleic acids research 51, W207–W212 (2023).

72 Fukuchi, M. et al. Neuromodulatory effect of Gαs-or Gαq-coupled G-protein-coupled receptor on NMDA receptor selectively activates the NMDA receptor/Ca2+/calcineurin/cAMP response element-binding protein-regulated transcriptional coactivator 1 pathway to effectively induce brain-derived neurotrophic factor expression in neurons. Journal of Neuroscience 35, 5606–5624 (2015).

73 Fukuchi, M. et al. Visualization of activity-regulated BDNF expression in the living mouse brain using non-invasive near-infrared bioluminescence imaging. Molecular brain 13, 122 (2020).

74 Ko, S. Y. et al. Systems consolidation reorganizes hippocampal engram circuitry. Nature, 1–9 (2025).

